# Cell shape regulates subcellular organelle location to control early Ca^2+^ signal dynamics in Vascular Smooth Muscle Cells

**DOI:** 10.1101/161950

**Authors:** R. C. Calizo, M. K. Bell, A. Ron, M. Hu, S. Bhattacharya, N. J. Wong, W.G.M. Janssen, G. Perumal, P. Pederson, S. Scarlata, J. Hone, E. U. Azeloglu, P. Rangamani, R. Iyengar

## Abstract

The shape of the cell is connected to its function; however, we do not fully understand underlying mechanisms by which global shape regulates a cell’s functional capabilities. Using theory, experiments and simulation, we investigated how physiologically relevant cell shape changes affect subcellular organization, and consequently intracellular signaling, to control information flow needed for phenotypic function. Vascular smooth muscle cells going from a proliferative and motile circular shape to a contractile fusiform shape show changes in the location of the sarcoplasmic reticulum, inter-organelle distances and differential distribution of receptors in the plasma membrane. These factors together lead to the modulation of signals transduced by the M_3_ muscarinic receptor/G_q_/PLCβ pathway at the plasma membrane, amplifying Ca^2+^ dynamics in the cytoplasm and the nucleus resulting in phenotypic changes, as determined by increased activity of myosin light chain kinase in the cytoplasm and enhanced nuclear localization of the transcription factor NFAT. Taken together, our observations show a systems level phenomenon whereby global cell shape affects subcellular organization to modulate signaling that enables phenotypic changes.

## INTRODUCTION

Cells utilize receptors on the plasma membrane to transduce a range of extracellular signals to regulate function in the cytoplasm and the nucleus ^1^. Reaction kinetics of the biochemical interactions that comprise the signaling networks regulate the temporal dynamics of activation and inactivation of signaling components and effectors ^2^. However, information flow within cells is not just temporally regulated, it is also spatially regulated by the shape of the cell ^3–5^ and surface-to-volume ratio at the plasma membrane ^6,7^. An additional layer of complexity is conferred by the spatial transfer of information by signaling reactions that occur within or at intracellular organelles. Recently, studies have shown that signaling at the endosomes plays an important role in prolonging cAMP dynamics through GPCRs ^8^ and in EGFR dynamics ^9^. An important compartmental regulation of organelle-based signaling is Ca^2+^ dynamics, since endoplasmic/sarcoplasmic reticulum is a regulatable Ca^2+^ store in cells. Ca^2+^ is a ubiquitous signaling molecule that controls many cellular functions including fertilization, proliferation, migration and cell death ^10–13^. Ca^2+^ is able to participate in controlling this diverse array of functions due to the precise control of Ca^2+^ concentration within the cell. In vascular smooth muscle cells (VSMC), Ca^2+^ regulates both contractility and gene expression through IP_3_-mediated Ca^2+^ release by IP_3_R receptors located on the membrane of the sarcoplasmic reticulum (SR) and through Ca^2+^ influx at the plasma membrane ^14–16^. Ca^2+^-calmodulin activates myosin light chain kinase (MLCK), which phosphorylates the light chain of myosin, initiating contraction ^17^. Ca^2+^ also activates protein kinases and phosphatases that regulate transcription regulators that define the phenotypic status of VSMC ^18^. Ca^2+^ activates calcineurin, which dephosphorylates the transcription factor nuclear factor of activated T-cells (NFAT) in the cytoplasm, resulting in its nuclear accumulation and expression of NFAT-regulated genes ^19^. Ca^2+^ also activates calmodulin kinase II (CaMKII), that phosphorylates the transcription factor serum response factor (SRF) ^20^ which, as a complex with myocardin, controls the expression of proteins necessary for the contractile function of VSMC ^21^.

VSMC in the medial layer of the walls of blood vessels are not terminally differentiated and can undergo phenotypic transitions during injury and disease states ^22–24^. VSMC shape and function are closely related; increasing elongation or aspect ratio (AR, defined as the ratio of the short axis to long axis) is correlated with differentiation and contractility ^25,26^. This fusiform shape represents the normal physiological state of contractile VSMCs ^27^. Previous studies have shown that changes in cell shape are associated with major changes in subcellular organization, which may affect cell physiology and phenotype including contractility, proliferation and differentiation states ^28–30^. Recent experiments have shown that local microdomains Ca^2+^ are possible because of hindered diffusion of IP_3_ ^31^ and differential subcellular distribution of IP_3_R ^32^. However, the mechanisms underlying the relationship between global cell shape and changes in cell physiology such as contractility, remain poorly understood.

Based on the observations that cell shape and Ca^2+^ signaling closely regulate the contractile phenotype of differentiated VSMCs, we hypothesized that cell shape regulates organelle location, including the relative distances between plasma membrane, endoplasmic/sarcoplasmic reticulum (ER/SR) and the nucleus, to modulate cellular functions. We sought to answer the question -- does cell-shape-dependent organelle localization drive spatial control of information flow and affect cellular function? The answer to this question is important for understanding how mechano-chemical relationships control cell shape, signaling, and phenotype over multiple length scales. We used 3D biochips to culture VSMCs in different shapes and found that different cell shapes have different PM-SR (plasma membrane - sarcoplasmic reticulum) distances. We then developed theoretical and computational models to represent the spatio-temporal dynamics of IP_3_/Ca^2+^ transients mediated by Muscarinic Receptor 3 (M_3_R)/Phospholipase Cβ (PLCβ) pathway. Activation of M_3_R mediates contractility in VSMC by activating PLCβ resulting in phosphoinositide hydrolysis and IP_3_ mediated Ca^2+^ release from the SR ^33,34^. We tested the effect of cell shape on VSMC contractility and found an unexpected modulation of organelle location as a function of cell shape and that this change in organelle location results in signal amplification in the cytoplasm and nucleus.

## METHODS

### Cell culture

A10 cells, which are VSMC from thoracic/medial layer of rat aortas, were obtained from American Type Culture Collection (CRL-1476). A10 cells were maintained in Dulbecco’s modified eagle’s medium (DMEM, Gibco), supplemented with 10% Fetal Bovine Serum, 1% penicillin/streptomycin, at 37 °C and 5% CO_2_. Cells were transfected using Neon Transfection System (Life Technologies) according to manufacturer’s instructions. Briefly, 5 × 10^5^ cells were electroporated with 1 µg DNA in suspension buffer, with the following electroporation settings: 1400 V, 2 pulses, 20 ms pulses each. Cells were then suspended in DMEM supplemented with 10% FBS and then allowed to adhere on 3D biochips. Forty-eight hours post-transfection, cells were imaged using Hanks Balanced Salt Solution (HBSS) supplemented with CaCl_2_, MgCl_2_ and 10 mM HEPES.

### Fabrication of 3D biochips

Patterned surfaces were fabricated by conventional photolithography using SU8 photoresist ^35^. Complete details are provided in ^86^ (see Figure 5 in this reference for details on ellipsoid shape). Briefly, cover glass slides were cleaned by sonication in isopropanol and deionized water for 15 minutes and baked at 110 °C overnight and photolithography was subsequently performed using standard vacuum hard-contact mode. Before plating of cells onto 3D biochips, microfabricated surfaces were washed with 50 μg/mL gentamicin and then incubated with 0.5% Pluronic for at least 3 hours. The 3D biochips were then washed with PBS and were seeded with cells.

### Immunofluorescence of cells in 3D biochips

Cells were seeded onto the 3D biochips and were allowed to adhere and comply with the patterns for at least 24 hours. After assay treatments, cells were fixed with 4% paraformaldehyde (Electron Microscopy Sciences) for 15 minutes at room temperature, washed with PBS, permeabilized with 0.2% saponin for 30 minutes and blocked with 4% normal goat serum doped with 0.05% saponin for 1 hour. Cells were then incubated overnight with primary antibodies (sources and catalog numbers shown in table below) that had been diluted in blocking solution at 4 °C. Cells were washed with PBS and samples were incubated with secondary antibodies (Alexa 488, Alexa 568 and/or Alexa 647) for 1 hour at room temperature. For organelle staining, cells were counter-stained with Actin Green and DAPI counterstains in addition to the secondary antibodies. Cells were then imaged on a Zeiss LSM 880 confocal microscope equipped with 63x 1.4NA oil immersion objective. Same acquisition settings were applied across different conditions that were compared (laser power, gain settings, magnification, zoom, pixel size and slice thickness). For quantitative immunofluorescence of M_3_R, a Z-stack of 30-40 slices using a slice thickness of 0.5 μm were obtained for each cell. Z-stack datasets were then pared down to 21-22 slices encompassing the entire height of the cell (mean cell height ~ 10 μm). Alignment, registration and cropping were performed to ensure each image had the same x-y dimensions (circular cells = 253 × 246 pixels, elliptical cells (AR 1:10) = 106 × 512 pixels). Per condition, images of cells obtained from the same z-plane (3.0 μm from the confocal slice corresponding to the bottom region of the plasma membrane), were averaged to obtain the averaged distribution of M_3_R in different regions of the cell. For immunofluorescence of transcription factors, multi-channel images consisting of DAPI, Actin Green (Invitrogen), and primary antibodies for NFAT, SRF or myocardin were aligned and stitched using the ZEN 2014 software. Image analysis and quantification was performed using ImageJ scripts. Briefly, nuclei were segmented in the DAPI channel. Corresponding cytosol and whole cell objects were outlined utilizing the contrast enhanced phalloidin channel to define cell boundaries. Nuclear-to-cytoplasmic transcription factor ratio was defined as the ratio of the mean transcription factor intensity colocalizing with the nuclear object divided by the mean intensity of the corresponding cytosol object. All measurements were exported directly to csv files and were subsequently analyzed using MATLAB to generate plots.

| **Antibody** | **Dilution** | **Source** |
| --- | --- | --- |
| Calnexin | IF 1:100 | Abcam, cat # ab22595 |
| Protein Disulfide Isomerase (PDI) (C81H6) | IF 1:100 | Cell Signaling, cat # 3501 |
| Nogo-A/Reticulon-4 | IF 1:100 | Cell Signaling, cat # ab47085 |
| α-tubulin | IF 1:100 | Cell Signaling, cat # 2144 |
| AIF mitochondrial marker (D39D2) | IF 1:100 | Cell Signaling, cat # 5318 |
| EEA1 – Early endosome marker (C45B10) | IF 1:100 | Cell Signaling, cat # 3288 |
| RCAS1 (D2B6N) Golgi marker | IF 1:100 | Cell Signaling, cat # 9091 |
| Muscarinic Acetylcholine Receptor 3 (M<sub>3</sub>R) | IF 1:100 | Abcam, cat # ab126168 |
| NFATc1 antibody | IF 1:100 | Abcam, cat # ab2722 |
| SRF antibody | IF 1:100 | Cell Signaling, cat # 4261 |
| Myocardin | IF 1:100 | Abcam, cat # 22073 |

### Airyscan Imaging of Live Cells

VSMC conforming in the 3D biochips were simultaneously labeled with 1 μM CellMask Plasma Membrane tracker (Life Technologies), 1 μM CellMask ER marker (BODIPY TR Glibenclamide), in HBSS buffer supplemented with 1% Pyruvate, 1% HEPES and 1 mM Trolox, for 5 minutes at room temperature. Images were acquired using Zeiss LSM 880 using Airyscan super-resolution imaging equipped with 63× 1.4 Plan-Apochromat Oil objective lens at 30ºC. Z-stacks with an interval of 0.15 μm were collected for the entire cell height which approximated 10-12 μm. Z-stack analyses and other post-acquisition processing were performed on ZEN Black software (Carl Zeiss).

### Calcium Measurements

VSMC were seeded on 3D biochips. Calcium measurements in 3D biochips were performed as previously described with modifications ^36^. Briefly, cells in 3D biochips were serum-starved for 12 hours and loaded with 5 μM of calcium green (dissolved in DMSO) for 30 minutes at room temperature, with Hanks Balanced Salt solution, (HBSS) supplemented with CaCl_2_, MgCl_2_ and 10 mM HEPES. Calcium Green was imaged using Zeiss 510 equipped with 40x Apochromat objective at acquisition frame rate of 4 fps (250 ms acquisition time), and Calcium Green was excited using Argon ion laser 488 at low transmittivity (1%) to prevent photobleaching. Image stacks acquired were then imported into Fiji/ImageJ. Background subtraction was performed on the time stacks by using a rolling ball radius of 50 pixels. Cytoplasm and nuclear regions of interest (ROI) were chosen by performing a maximum intensity projection of the time-stack and specifying a 5 μm radius circle within the the nuclear and cytoplasmic regions. To convert intensity values to Ca^2+^ concentration, modified Grynkiewicz equation was used, defined as: 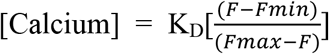. Where *F*_*min*_ is the average fluorescence intensity of the ROI after addition of 100 μM BAPTA AM, *F*_*max*_ is the average fluorescence intensity of the ROI after addition of 0.100 μM A23187. Integrated Ca^2+^ was obtained using the trapz() function in MATLAB.

### FRET imaging

MLCK-FRET plasmid is a kind gift from Dr. James T. Stull (University of Texas Southwestern Medical Center). The MLCK-FRET plasmid is a calmodulin-binding based sensor, where calmodulin binding sequence is flanked with eCFP and eYFP and exhibits decreased FRET upon binding with calmodulin ^19,37^. Cells expressing MLCK-FRET were imaged using Zeiss LSM 880 (Carl Zeiss, Jena, Germany), at 37 °C incubator, fitted with Plan-Apochromat 20x, equipped with 458 nm and 514 nm Argon ion laser lines for excitation of eCFP and eYFP respectively. Incident excitation light was split using an MBS 458 nm/514 nm beam splitter and collected on a 32-spectral array GaAsp detector. The fluorescence emission was collected from 463-520 nm (ECFP), 544-620 nm (FRET channel and eYFP channel). Intensity based ratiometric FRET were obtained using custom-written scripts in ImageJ and MATLAB. Since MLCK-FRET is a single-chain construct, decrease in FRET, and increase in MLCK binding to calmodulin, was expressed as the ratio of emission intensity at 520 nm/emission intensity at 510 nm normalized at the basal levels.

### NFAT imaging

HA-NFAT1(4-460)-GFP was a gift from Anjana Rao (Addgene plasmid # 11107). Patterned cells expressing NFAT-GFP was imaged using Zeiss LSM 880, using Argon ion laser 488 nm, as described above and 63X 1.4 NA oil objective, with an acquisition rate of 1 frame every 10 seconds. Time series image stacks were analyzed using ImageJ. Regions of interest of identical size were drawn in the cytoplasmic and nuclear regions of interest and the ratios of these intensities were computed over time.

### Electron Microscopy

3D biochips containing fixed A10 cells were embedded in Embed 812 resin (Electron Microscopy Sciences (EMS), Hatfield, PA) using the following protocol. Cells were rinsed in 200mM phosphate buffer (PB), osmicated with 1% osmium tetroxide/PB, washed with distilled water (dH20), and *en bloc* stained with aqueous 2% uranyl acetate, washed with dH2O and dehydrated via increasing ethanol (ETOH) series /distilled water (25%, 60%, 75%, 95% and 100% ETOH). Cells were further dehydrated using propylene oxide (PO), and embedded using ascending PO:EPON resin concentrations (2:1, 1:1, 1:2, pure). Prior to solidification, the coverslips were placed on 1”X 3” microscope slides, and multiple open ended embedding capsules with a 1 × 1 cm face (EMS) were placed on the coverslips covering the areas of interest. The resin was then polymerized in a vacuum oven at 65 °C for 8–12 hours. After the first layer was solidified, the capsule was topped off with more resin and put back in the oven for another 8–12 hours. Capsules containing cells within 3D biochips were removed from coverslips using previously described methods ^38^. Briefly, to separate the block from the coverslip, a hot plate was heated to 60°C and the microscope slide was placed directly on a pre-heated hot plate for exactly 3 minutes and 30 seconds. The slide was removed from the hot plate and the capsules carefully dislodged free from the coverslips. Once separated, the block face retains the cells within the 3D biochips. The block was coarsely trimmed with a double-edged razor blade, and a Diatome cryotrim 45° mesa-trimming knife (EMS) was used to finely trim the block. Using as large a block face as possible, 70 nm ultrathin sections were cut from the block surface using an ultra-thin diamond knife (EMS), and a Leica EM UC7 ultramicrotome (Buffalo Grove, IL). All sections coming off the block face were collected. Sections were collected using a Perfect Loop (EMS) and transferred to a 2 × 1 mm formvar-carbon coated reinforced slot grid (EMS). The sample was dried on the grid and transferred to Hiraoka Staining Mats (EMS) for heavy metal staining. Grids were stained with 3% uranyl acetate in water for 40 minutes, washed and stained with Reynold’s lead citrate for 3 minutes, washed and allowed to dry. Electron microscopy images were taken using a Hitachi 7000 Electron Microscope (Hitachi High Technologies America, Inc.) equipped with an AMT Advantage CCD camera. Cells were viewed at low-magnification to identify areas of interest (cell tip versus cell body) before high magnification imaging. Images were transferred to Adobe Photoshop CS3 (version 10), and adjusted for brightness and contrast. Measurement of plasma membrane to ER distances from electron microscopy images were performed blindly. Briefly, sample information from images were removed and images were saved with a randomized filename. Image contrast was further enhanced using ImageJ using contrast-limited adaptive histogram equalization (CLAHE). Only images with discernible smooth ER closely apposed to the plasma membrane were analyzed and distances were measured at optimal xy orientations at 50 nm intervals using ImageJ. Data was graphed using MATLAB. For 3D reconstruction of ER-PM distances, 3D serial block face scanning electron microscopy was performed following the above protocol with the following modifications: prior to sample embedding, high-contrast tissue pre-staining was performed using 2% Osmium tetroxide, 2.5% potassium ferricyanide, 1% thiocarbohydrazide, and 1% uranyl acetate in cacodylic buffer. Blocks were serial sectioned and imaged at 1600X magnification using a 3View Gatan system coupled with a Zeiss Gemini FE scanning electron microscope (Carl Zeiss Microscopy, LLC). Final tiled pixel frame store of 32k-by-32k with a 650 μm field of view lead to lateral and axial resolutions of 20 and 75 nm, respectively. ER-PM distances were manually segmented for each axial image by selecting dyads perpendicular to the cell surface in a blinded fashion.

### Statistics

Results are presented as mean ± standard error of the mean from at least three independent experiments. Normality was determined using Shapiro-Wilk test using a p-value ≥ 0.05. If the distribution was normal, a two-tailed Student’s t-test was performed. For datasets with non-normal distribution, two-tailed Mann-Whitney test was used. P < 0.05 was considered statistically significant.

### Model Development

We formulated a phenomenological model to study the role of PM-SR distances. The complete derivation and mathematical solution is given in the Supplementary Information. Complete details of the simulations using finite-element methods in COMSOL and using finite-volume methods in *Virtual Cell* are given in the Supplementary Information.

### Data Availability

Data supporting the findings of this study are available within the paper and its supplementary information.

## RESULTS

### Reaction-diffusion model predicts that the distance between the PM and SR membrane affects signaling dynamics in the cytoplasmic volume

We tested the hypothesis that a change in membrane curvature and distance between two membranes affects signaling dynamics using a phenomenological reaction-diffusion model in COMSOL (Fig. 1a). A signaling molecule of interest, C_A_, is produced at the PM with an on-rate k_on_ (μm/s) and binds to a receptor located at the SR membrane, with a rate k_off_(μm/s) and is free to diffuse in the sandwiched cytoplasmic space and is degraded by a degrading enzyme with a rate k_deg_ (1/s). This model essentially captures the lifecycle of a second messenger such as IP_3_ that is produced at the PM through PIP_2_ hydrolysis by phospholipases, binds to inositol 3 phosphate receptor (IP_3_R) channel at the SR membrane and is degraded in the cytoplasm by 1,4,5-trisphosphate-5-phosphatase^32^. These events can be mathematically represented by the following system of reaction-diffusion equations. The dynamics of C_A_ in the cytoplasm are governed by diffusion and degradation and is given by,

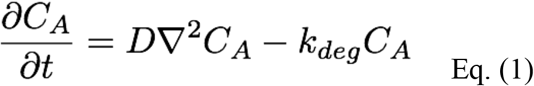

Here, D is the diffusion coefficient of C_A_(μm^2^/s), and C_A_ is the concentration of C_A_ (μM). The boundary condition at the PM is a balance between the rate of production of C_A_ at the membrane and the diffusive flux from the membrane to the cellular interior and is given by,

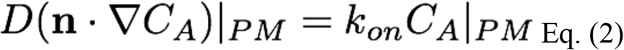

Here, n is the normal vector to the surface and ∇ represents the gradient operator. At the SR membrane, similarly, we can write the boundary condition for the consumption of C_A_ as the balance of diffusive flux to the SR and consumption rate at the SR.

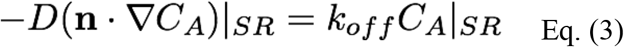

**Figure 1.**
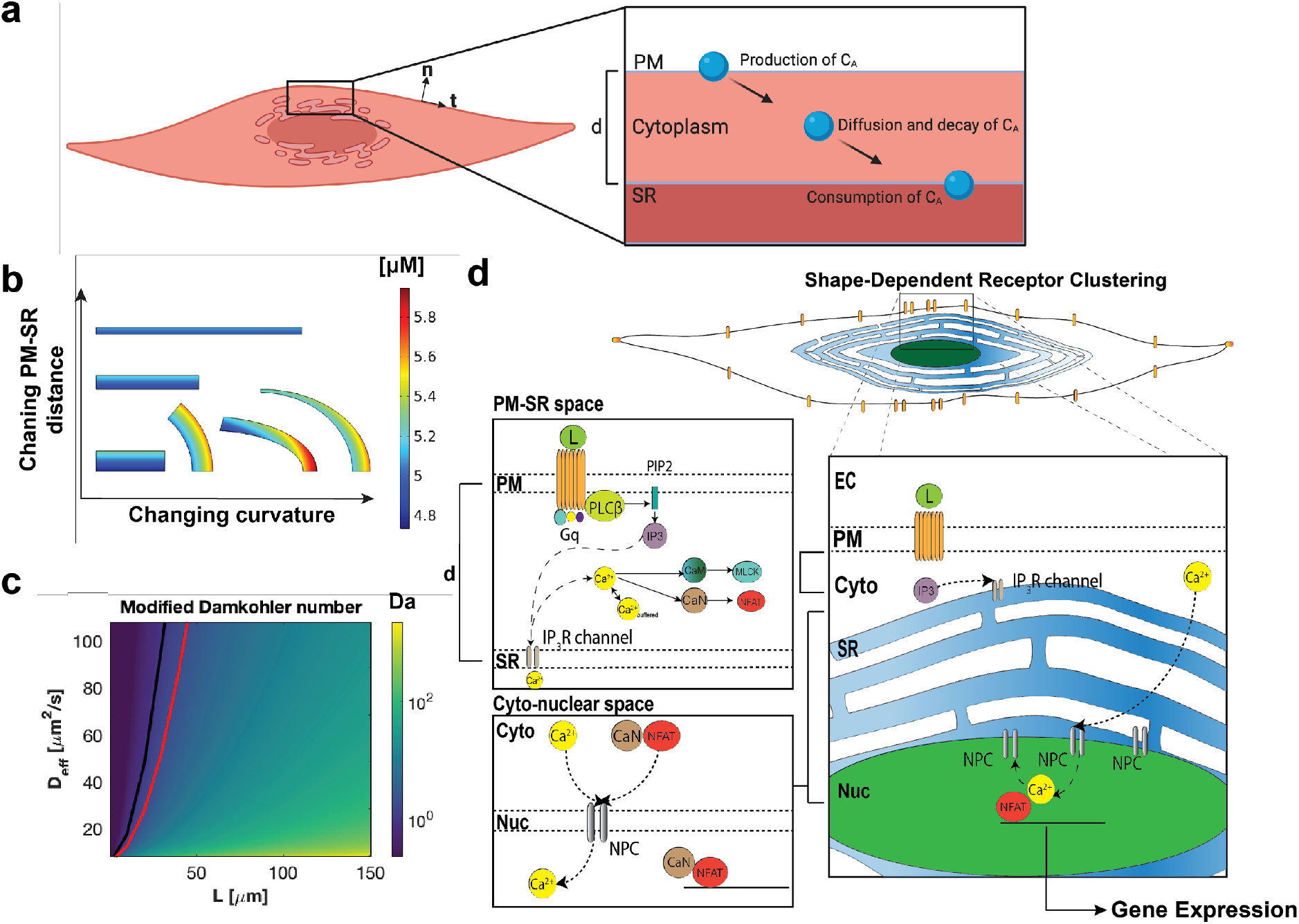
Phenomenological model describing the dynamics of a signaling molecule produced at a membrane and consumed at an organelle membrane compartment as a function of curvature and PM-SR distance. (a) Dynamics of a signaling molecule can be described by its production, diffusion, degradation in the cytoplasmic space, and consumption at an intracellular organelle membrane. Both curvature (denoted as t and n) and PM-SR distance differences (denoted as d) can play important roles in signaling dynamics. The interplay between sources, sinks, and boundary fluxes determines signaling dynamics. Schematic created with Biorender.com. (b) Six different geometries with the same area were used to test how curvature of the membrane and distance between the plasma membrane (PM) and the inner membrane, a sarcoplasmic reticulum (SR) in this case, affect the spatio-temporal dynamics of the signaling molecule ‘C_A_’. Reaction-diffusion modeling in COMSOL predicts that curvature and distance between the PM and SR will both affect the spatial distribution of C_A_. Here we show transient snapshots at 50 ms for a toy model of C_A_ as defined in Eqs. 1-3 where k_deg_ = 0.5 [1/s], k_on_ = k_off_ = 7.5e-10 [μm/s], D_Ca_ = 1 [μm^2^/s], and Ca_cytoIC_ = 5 [μM] is the initial concentration of calcium everywhere in the cytosol. The top surface for the rectangles and rightmost surface for the curved sections is the PM as the emission surface, and the bottom surface for the rectangles and leftmost surface for the curved sections is the SR membrane as the absorption surface. The concentration ranges from ~4.7 to 5.9 μM. (c) The modified Damkohler number for different values of PM-SR distance, ‘L’, and effective diffusion coefficient, ‘D_eff_.’ k is taken as 0.1 1/s in this example. For physiological distances shown in this study, the dynamics of A are dominated by diffusion. The threshold between reaction-dominated and diffusion-dominated regimes is defined at Da = 1, shown as a black line. If a system has diffusion-trapping, for the same diffusion coefficient, the system can shift regimes due to geometric and kinetic trapping factors (compare black line to red line). Therefore it is clear that considering effective diffusion could shift the system between a diffusion-dominated to reaction-dominated system, particularly for small crowded intracellular regions. d) Biochemical events that govern Ca^2+^ signaling in VSMC including GPCR-G_αq_ cycling on the membrane, PLCβ activation, IP_3_ production and IP_3_R-mediated Ca^2+^ release from SR.

We note that C_A_ = 0 is the trivial solution to this system, so we assume a nonzero initial condition. For illustrative purposes, we solved these equations using finite element methods on six geometries (1) a control rectangle (constant distance, zero curvature) (2) two additional rectangles with different PM-SR distances, (constant distance, zero curvature), (3) a circular sector (constant distance, constant curvature), and (4) an elliptical sector, (constant distance, varying curvature), and (5) a elliptical sector (varying distance, varying curvature) (Fig. 1b). In cases (1-3), the gradient of C_A_ is only along the normal direction (Fig. 1b). Cases (4-5), where curvature varies and both curvature and PM-SR distance vary, results in two-dimensional gradients. The curvature of the membrane and the PM-SR distance will affect both the production and consumption of C_A_ at the PM and SR respectively. Hence, C_A_ varies both in the radial and angular directions, indicating that curvature and varying distances between the two membranes amplifies signaling gradients. Even though the observations here are in 2D, the effect of curvature and distance holds true in 3D as has been demonstrated by us and others ^3,39,40^.

Dimensional analysis of Eq. 1 produces the Damkohler number (*K*_*deg*_*L*^2^/*D*), a nondimensional number relating the contribution of reaction rate to diffusive transport, which identifies the system as either diffusion- or reaction-dominated ^41^. Considering the model system, species C_A_ travels a distance, L (d in Fig. 1), from the PM to the SR during which time it can be degraded at some rate k_deg_. However, the cell cytoplasm is a dense mixture of proteins, ions, and other cellular components that can interact with and trap species C_A_ in various direct or indirect ways ^42,43^. This trapping, whether through binding or diffusion barriers, produces an effective diffusion coefficient that can be written as 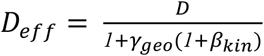, where *γ*_*geo*_ represents the geometric effects of traps and *β*_*kin*_ represents kinetics of binding at traps if applicable ^42^. Inserting this effective diffusion coefficient into the Damkholer number provides a more realistic analysis of the species C_A_, accounting for diffusion trapping, consistent with approaches from statistical physics. Diffusion-trapping can modify the effects of diffusion however, altering the dynamics of the system. In Fig. 1c, the black line denotes Da = 1, the usual threshold between reaction-dominated and diffusion-dominated regimes. If one considers two cases for the same diffusion coefficient, one without diffusion-trapping (black line) and one with diffusion-trapping (red line), it is clear that considering effective diffusion can shift the system between reaction- and diffusion-dominated regimes, particularly in crowded intracellular regions.

From this simple phenomenological model, we predict that second messenger signaling pathways such as receptor/phospholipase C/IP_3_, where signaling occurs between PM and SR, will be impacted by both cell shape and distance between the PM and the SR. This prediction raises the following questions: (1) does cell shape affect PM-SR distance? (2) Does changing PM-SR distance affect intracellular signaling dynamics? And (3) what is the impact of changing the distances between organelles on VSMC contractility through MCLK activity?

We asked whether changing the PM-SR distances can impact the dynamics of IP_3_ signaling. Since it is currently not possible to experimentally manipulate PM-SR distances with precise control in cells, we used numerical models of IP_3_ and Ca^2+^ in the context of VSMC to study the relationship between PM-SR distances and IP_3_ dynamics. This model is composed of a system of multi-compartmental partial differential equations, representing the reaction-diffusion of cytoplasmic species in the volume, coupled with boundary fluxes at the membrane. The reactions capture the biochemical interactions from the ligand binding and activation of M_3_R to IP_3_ production by PLC-mediated hydrolysis of PIP_2_ release from the SR (Fig. 1d, subset of Supplementary Table 1–2, R1-R16). The simulations were conducted in the commercially available finite element software COMSOL to enable investigation of small PM-SR distances ranging from 50-150 nm.

In order to investigate how PM-SR distances affect IP_3_ dynamics, we constructed a cylindrical geometry to represent a portion of the cell and the ER as discs (Fig. 2b). This idealized geometry enables us to study reaction-diffusion in 3D using COMSOL while gaining insight into the stacked ultrastructure of the SR. We varied the following parameters: (a) PM-SR distance, (b) diffusion constant of IP_3_, and (c) IP_3_ degradation rate. These three parameters govern the production, diffusion, and consumption of IP_3_. Our results can be summarized as follows – across all parameter variations, shorter distances between PM-SR gave rise to high IP_3_ gradients (Fig 2b i, iv, and vii, Supplementary Fig 7, Supplementary Fig 11). Increasing the diffusion coefficient resulted in rapid dissipation of the gradient and increasing IP_3_ degradation rate resulting in a lower concentration of IP_3_, as expected. We found that the shortest distance between the PM-SR (50 nm), with the smallest diffusion coefficient (10 μm^2^/s), and largest IP_3_ degradation rate (0.0625 s^−1^) (Fig. 2c vii, top row) had the strongest IP_3_ gradient at 5s. Increasing the PM-SR distance to 100 nm altered the extent of the observed IP_3_ gradient at 5s (Fig. 2c i, bottom row) but did not alter this gradient very much at later times. From these simulations, we predict that changing the distances between organelles can alter IP_3_/Ca^2+^ signaling dynamics and potentially VSMC contractility as measured by MLCK activity. We used VSMCs grown on 3D biochips to test these predictions.

**Figure 2.**
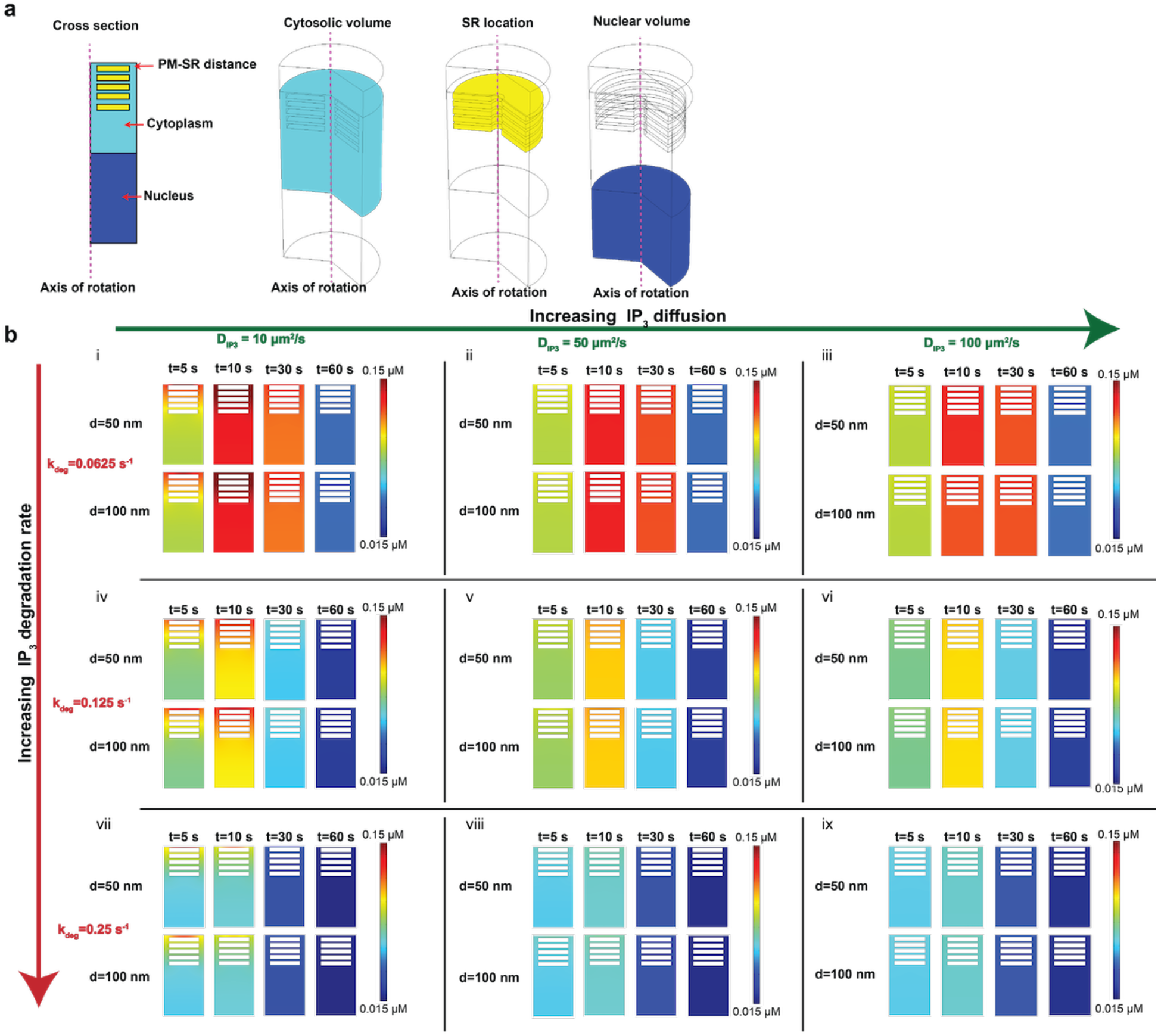
Small length scale simulations in COMSOL show a critical relationship between IP_3_ and PM-SR distance. (a) Cross-section and geometries representing different compartments governing biochemical events. We model a small part of the cell as a cylinder. The cytosolic volume is shown in cyan, the SR modeled as cylindrical disks shown in yellow and the nucleus is shown in blue and the axis of symmetry is shown in pink. The PM-SR distance, denoted as d in Fig 1d, is measured between the plasma membrane and the first SR disk. b) Effect of IP_3_ diffusion and degradation rate are shown for two different distances between the PM and the SR. Panels (i-iii) show the effect of increasing IP_3_ diffusion ((i) 10 μm^2^.s^−1^, (ii) 50 μm^2^.s^−1^, and (iii) 100 μm^2^.s^−1^) on IP_3_ concentration at 5s, 10s, 30s, and 60s for a degradation rate k_deg_= 0.0625 s^−1^ for two different values of d (50 nm and 100 nm). Panels (iv-vi) show similar calculations for k_deg_ =0.125 s^−1^ and panels (vii-ix) for k_deg_=0.25 s^−1^.

### Cell shape also affects cytoskeleton organization

We determined if changing the cell shape affects the organization of the cytoskeleton and the distribution of other organelles. In order to control the large scale cell shape, we used 3D biochips with the same surface area but increasing aspect ratio (AR, circles 1:1 to ellipses 1:8) (Fig. 3a). We investigated how cell shape affects cytoskeletal organization since the two are tightly interwoven ^44–46^. Actin stress fibers increasingly oriented themselves along the long axis of the cell as the aspect ratio increased (Fig. 3b), indicating that the cells were responding to the mechanical forces and tension exerted by the substrate ^47^. It has been shown in several cell types that nuclear shape is tightly coupled to cell shape ^48^. Nuclear aspect ratio increased with cellular aspect ratio (Fig. 3c-d) while nuclear size (area, in μm^2^) decreased with increasing cell aspect ratio (Fig. 3e). The nuclei became increasingly oriented along the major axis of the cell as the whole-cell aspect ratio increased, evidenced by polar graphs showing the orientation histograms of nuclei of cells in increasing cell AR (Fig. 3f). Circular shaped cells showed a random nuclear orientation whereas increasing the cellular aspect ratio progressively oriented the nucleus in the geometric center of the cell. PM-nuclear distance in the major axis of the cell increased with cell aspect ratio (Fig. 3h) while the PM-nuclear distance in the minor axis decreased with cell aspect ratio (Fig. 3i). These results indicate that in VSMC, cell elongation resulted in nuclear elongation, reduced nuclear size, and a decrease in the PM-nuclear distance in the minor axis of the cell (*i.e.* near the center of the cell). In addition to the nucleus, spatial arrangement of the microtubules also changed with aspect ratio. Microtubules became highly aligned and increasingly sparser in the cell tips compared to the cell body as the cell aspect ratio increased (Supplementary Fig. 1). Because cytoskeletal organization plays an important role in the distribution of many organelles ^49,50^, we visualized the effect of cell shape on the bulk distribution and location of the mitochondria (Supplementary Fig. 2), endosomes (Supplementary Fig. 3) and Golgi membrane (Supplementary Fig. 4). No clear trends were seen in the distribution of these organelles for different cell shapes. It has been reported that well-differentiated VSMCs have a characteristic central distribution of the endomembrane system ^25^. Because SR stores Ca^2+^, which controls both excitation-transcription coupling and contractility in VSMC ^51^, we focused on SR distribution as a function of cell shape, using calnexin (Fig. 3j), protein disulfide isomerase (Fig. 3k), reticulon-4 (Fig. 3l) and bodipy glibenclamide (Supplementary Fig. 5) as SR markers. All four markers show that in circular cells, the SR was spread uniformly throughout the cell; increased aspect ratio induced the SR to localize in the perinuclear region and become significantly sparser, and mainly tubular, in the cell tips (Fig. 3j-l, Supplementary Fig. 5). Upon close inspection of Airy scan images of these SR markers, we qualitatively observe that in circular cells, the SR appeared to be equidistant from the plasma membrane along the periphery, i.e. there were no angular variations of PM-SR distance, while elliptical cells show a large angular variation in the PM-SR distance (Fig. 3d-e bottom panels and Supplementary Fig. 5, inset). Thus, we show that cell shape affects not only cytoskeletal organization, but also affects nuclear shape and PM-nuclear distances, and organelle distribution.

**Figure 3.**
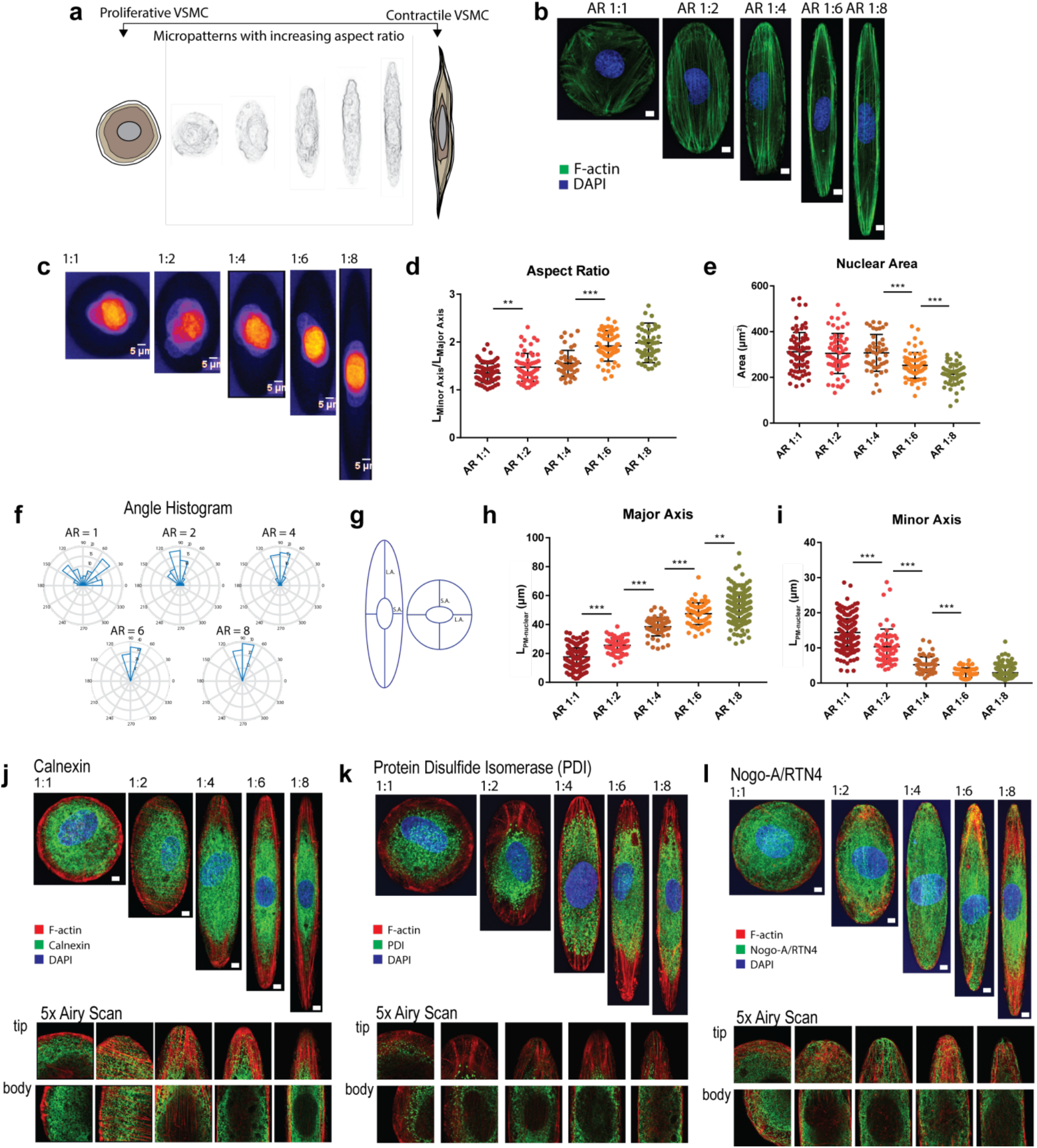
Cell shape impacts cytoskeletal organization and organelle location. Whole cell shape changes location and distribution of subcellular organelles in a systematic manner. (a) Biomimetic, microfabricated surfaces with graded aspect ratios fine-tune the shape of VSMC and its enclosed organelles. (b) Actin organization is dependent on cell shape. (c) Cell shape affects nuclear shape, size and distance to PM. Representative images of VSMC compliant in patterns with increasing AR stained for DAPI. Scatterplots of whole aspect ratio versus (d) nuclear aspect ratio (e) size (area μm^2^) (f) orientation with respect to the major axis of the cell (*N*_*AR1:1*_=*82, N*_*AR1:2*_=*60, N*_*AR1:4*_=*45, N*_*AR1:6*_=*65, N*_*AR1:8*_=*54*). (g) Relationship between whole cell shape and PM-nuclear distance was determined by segmenting cell and nuclear boundaries with Actin Green and DAPI respectively. Distances between nuclear and plasma membrane boundaries were measured along the long and short axes of the nucleus. (h) Increasing the cellular aspect ratio increases the distance between the nucleus and the plasma membrane in the major axis of the cell while (i) decreasing the PM-nuclear distance in the minor axis of the cell (**P ≤ 0.05, ***P ≤ 0.0001, two-tailed t-test). (j-l) SR distribution in VSMC seeded in AR 1:1 to 1:8 staining for SR membrane markers (panels below show 5x magnified Airy scan images of tip and body of images shown) (j) calnexin (k) protein disulfide isomerase (PDI) and (l) reticulon-4. Scale bars shown are 10 μm.

### Cell shape changes PM-SR distances in 3D

Two-dimensional images of peripheral SR membrane and plasma membrane indicate that cell shape alters SR abundance and PM-SR distance on the equatorial region (Fig. 3j-l). We investigated more closely whether PM-SR distances are altered by the global shape of cells in 3D. Confocal volume scans of VSMC cultured in circular and ellipse patterns show increased colocalization between calnexin and plasma membrane stains in the equatorial region of elongated cells, as opposed to the cell tips of elongated cells and circular cells, suggesting increased proximity of the two membranes at the equatorial region of elongated cells (Fig. 4a). Superresolution STED measurements reveal that SR puncta are closer to the cell cortex in the equatorial region of elongated cells (upper right panel, Fig. 4b), compared to both cell tips of elongated and circular cells (upper left and lower right panels, Fig. 4b). To quantitatively confirm that the PM-SR distance is shape dependent, we used transmission electron microscopy to visualize peripheral SR to PM distance (Supplementary Fig. 6). TEM showed that the cell periphery of circular cells and the cell body of elliptical cells showed long patches of smooth SR that were positioned close to the plasma membrane. PM-SR distance was indeed dependent on the shape of the cell: PM-SR distance in the cell body of elliptical cells was significantly smaller compared to circular cells (Fig. 4c). In the tips of elliptical cells, the SR membrane formed fewer contacts with the plasma membrane, and showed significantly higher PM-SR distance (Fig. 4d). 3D serial block face scanning electron microscopy reveals that the distance between puncta on the PM and SR is aspect ratio dependent (Fig. 4e left versus right). The difference between SR-PM distance in circular and elliptical cells was greatest within the first few microns of the basal surface, and it was progressive reduced towards the apical surface (Supplementary Fig. 20). However, we observed that for a given cell shape, the effect across was small when compared to differences between cell shapes (Fig. 4f). These results are consistent with the recent observation in neurons that PM-ER contacts are more extensive in the cell body compared to elongated projections such as dendrites and axons ^52^. While other groups have reported that cell shape can affect organelle location ^48,53^, here, we quantitatively determine PM-SR distances and the SR abundance along the juxtamembrane region with controlled cell shape variation.

**Figure 4.**
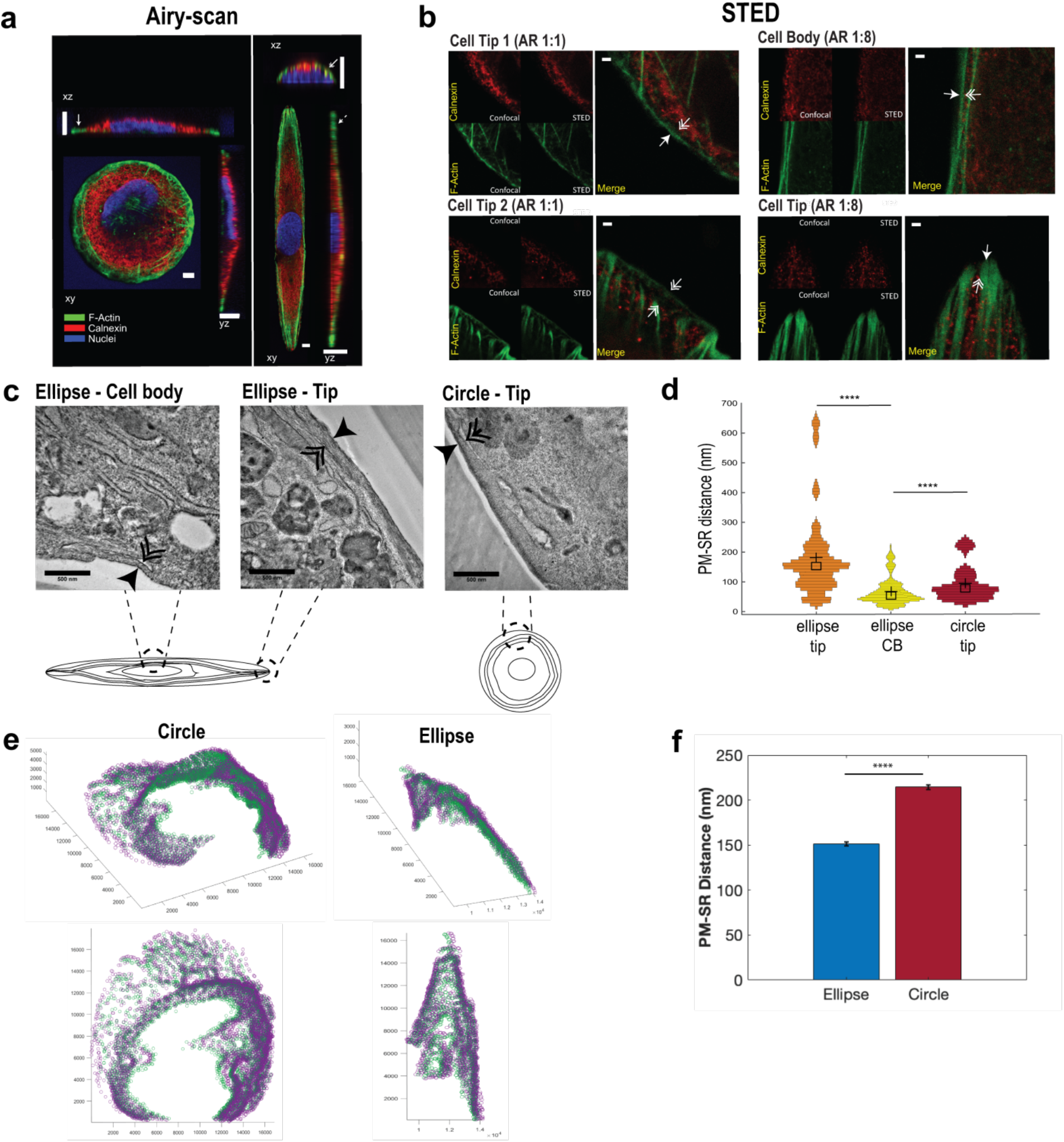
Shape impacts PM-SR distances in 3D. a) (Left panel) 3D confocal volume scans of VSMC cultured in circle and ellipse (Aspect ratio of 1:8) patterns showing F-Actin (green), ER membrane marker calnexin (red) and nuclei (blue). Increased co-localization of red and green pixels are seen in the xz (short axis) cross-section of elliptical cells (solid double arrow heads), while little co-localization is evident in tips of both circular and elliptical cells (solid single-arrow heads and dashed single-arrow heads, respectively). Scale bars shown are 10 μm. b) (Right panel) Super-resolution STED imaging of F-actin (green), calnexin (red), in cell tips of circular cells (AR 1:1) and in cell tip and cell body of elliptical cells (AR 1:8). Scale bars shown are 1 μm. c) Representative TEM micrographs sampled from corresponding regions in micropatterned cells (cartoon). Single-headed arrows indicate the plasma membrane and double-arrows indicate smooth, peripheral SR, apposed to the plasma membrane. Scale bars shown are 500 nm. (d) Distribution plots of PM-SR distances obtained from TEM images, mean and median are shown as crosses and squares respectively. (*N*_*ellipse tip*_=329, *N*_*cell body*_ = 254, *N*_*circle tip*_ = 115, ****P < 0.0001, two-tailed Mann-Whitney test). (e) In order to quantify the 3D distribution of PM-SR distances, spatial dyads were manually segmented in each axial image for representative circular and elliptical cells. Green points represent the SR segments while purple points represent the PM. (f) The average PM-SR distance and standard error mean (SEM) for a representative circular and elliptical cell was computed from the 3D serial block face scanning electron microscopy images (N_ellipse_ = 1692, N_circle_ = 5309, N refers to PM-SR distances taken within the representative cells, ****P < 0.0001, two sample t-test).

### Cell shape affects receptor activation on the membrane and intracellular calcium dynamics

We tested the model predictions that distance between PM and SR can affect the dynamics of Ca^2+^ mediated by the M_3_R/IP_3_/Ca^2+^ pathway ^14,54^ using computational and experimental methods. We stained for M_3_R in circular and elliptical cells under three different conditions - unstimulated, stimulated with carbachol, and stimulated with carbachol in the presence of hypertonic sucrose, which inhibits receptor endocytosis ^55–57^. We also constructed 3D spatial models of different aspect ratios in *Virtual Cell* (VCell) from carbachol stimulation to MLCK activation (see Supp for model details). We switched from COMSOL to *Virtual Cell* for the following reasons: *Virtual Cell* allows for experimental cell images to be imported and used as geometries for the computational simulations (see Supplementary Fig 10 and Material for more information on model geometries). This allows us to make closer comparisons between experiments and simulations. *Virtual Cell* is also computationally less expensive than COMSOL and is compatible for running simulations that involve whole cells. While we are unable to resolve the PM-SR distances in great detail in *Virtual Cell*, the benefits of conducting simulations in realistic geometries and for the entire signaling cascade were deemed significant to make the switch.

Experimental results show that in both the basal state and stimulated states, M_3_R was uniformly distributed on the plasma membrane of both circular and elliptical cells (Supplementary Fig. 8). Interestingly, in elliptical cells, when M_3_R was stimulated and endocytosis was inhibited, M_3_R accumulated in the cell body compared to the cell tips while there was no observable spatial asymmetry in the distribution of M_3_R in circular cells (Supplementary Fig. 8 and 12), consistent with our previous observations and modeling^3^. We then investigated the effect of shape on cytoplasmic Ca^2+^ dynamics upon stimulation of M_3_R in patterned VSMC (Fig. 5a-c). In the cytoplasmic region, circular and elliptical showed similar peak Ca^2+^ amplitudes. However, elliptical cells showed a slower rate of decrease in Ca^2+^ compared to circular cells, resulting in a reproducibly observable higher temporally integrated Ca^2+^ compared to circular cells, although the differences were near the threshold for statistical significance (p=0.057, two-tailed t-test n=14).

**Figure 5.**
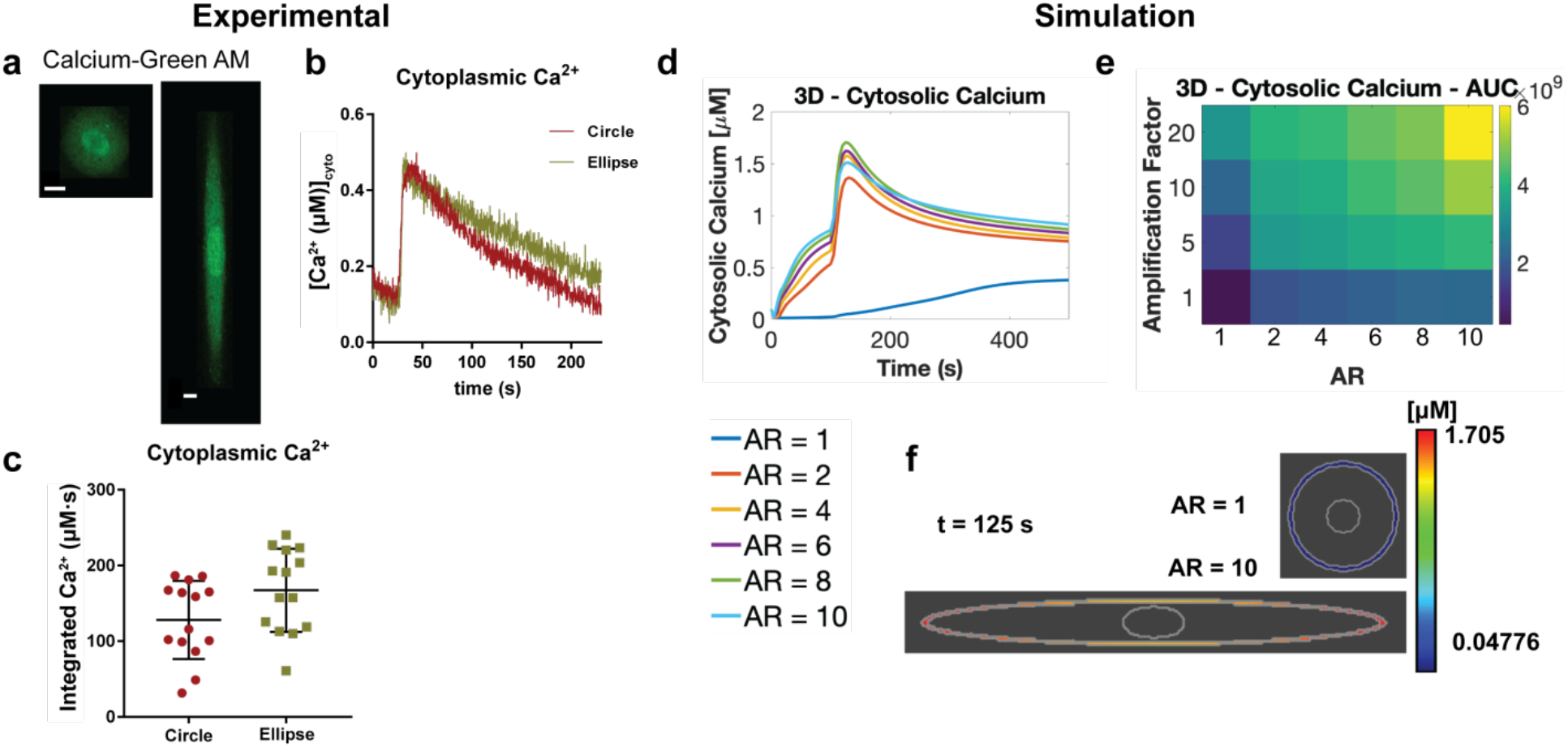
Cell aspect ratio affects cytosolic calcium levels. (a) Circular and elliptical cells were loaded with Calcium-Green AM dye, to measure Ca^2+^ signaling in response to 10 μM carbachol. Scale bars shown are 10 μm. (b) average time-course of cytoplasmic Ca^2+^ in response to CCh (circle, N=14, oval, N=14, stimulated at arrow indicated) (c) Scatterplot of cytoplasmic integrated Ca^2+^ (area-under-the-curve) per cell in circular and elliptical cells. Temporal (d) and AUC results (e) of cytosolic calcium for simulations of various aspect ratios and SR flux amplification factors in *Virtual Cell*. Inset below: legend for temporal simulation results showing the various aspect ratios, AR. (f) Cytosolic calcium spatial results for AR of 1 and 10 at 125 s.

Recent ultrastructural analyses have shown that the ER is a highly dynamic, tubular network that occupies roughly 10% of the cytosolic volume and extends from the nucleus to the cell periphery ^58^. Currently, geometries used in spatial models typically do not capture the complexity of the organelle ultrastructure ^52^, however, advancements in meshing algorithms and modeling pipelines pave the way for more complex geometries as model inputs ^59,60^. To represent the potential effect of the convoluted ultrastructure of the SR membrane, we introduce an SR flux amplification factor to our model. This amplification factor scales the SR flux term for IP_3_ activated calcium release from the SR, R17 in Supplementary Table 1^39^. This amplification factor provides some insight into how the spatial system would be affected by a modified flux term due to differences in SR geometry, specifically folds in the SR membrane, and allows us to explore the signaling consequences of varying the SR surface area. As a result, we can investigate the role of larger flux through the SR membrane without explicitly changing the SR model geometry (see Supplementary Material for more details). Model results (Fig 5d-f) show a similar trend of higher cytoplasmic Ca^2+^ in larger ARs compared to smaller AR. A parametric sensitivity analysis of the model reveals that the rate of Ca^2+^ release from the SR due to IP_3_R is a key parameter in the model (Supplementary Figs. 16–19). We investigate this geometric complexity through the aforementioned amplification factor. Increasing this amplification factor, which effectively captures a large surface area of the SR, also leads to increasing cytoplasmic Ca^2+^. Therefore, cell shape, SR surface area, and the PM-SR distance together influence short-timescale signaling events associated with the muscarinic receptor pathway.

### Cell shape affects downstream signaling dynamics

Since small changes in calcium signals can have large functional effects as a result of amplification by downstream signaling networks ^61^, we predicted that changes in cytoplasmic calcium levels can lead to additional downstream signaling changes. Our model predicts that higher ARs and larger shape factors will lead to large nuclear Ca^2+^ peaks and AUC (Area under the curve) (Fig. 6a,b,e). AUC is a common measure of signaling over time that quantifies accumulated signaling in the cell ^87^. We measured downstream effector activities in the cytoplasm and in the nucleus to verify these predictions experimentally. We first measured nuclear Ca^2+^ levels in circular and elliptical cells (Fig. 6). The differences in nuclear Ca^2+^ between circular and elliptical cells were distinct from cytoplasmic Ca^2+^ (Fig. 6c). In the nuclear region, the peak Ca^2+^ amplitudes of circular and elliptical cells were similar. However, there is a notable delay in the rate of nuclear Ca^2+^ increase to maximum in elliptical cells compared to circular cells, and the decay times in elliptical cells were slower as well, resulting in a significantly higher concentrations of temporally integrated Ca^2+^ in elliptical cells compared to circular cells (Fig. 6d). These results indicate that cell shape affects nuclear Ca^2+^ transients triggered by M_3_R/PLCβ and that such transients are more prolonged in elliptical cells. We similarly predicted elevated activated MLCK AUC and peaks for higher AR and shape factor (Fig. 6f,g,j) because MLCK is an immediate downstream effector of Ca^2+ 62,63^. We measured myosin light chain kinase (MLCK) activity using a CaM-sensor FRET probe ^37,64^ (Fig. 6h,i,k) to obtain estimates of actin cytoskeletal dynamics associated with contractility. Elliptical cells showed a higher degree of MLCK FRET probe localization on actin filaments (Fig. 6k) and higher maximal activation compared to circular cells (Fig. 6h,i), indicating that the shape-induced increase in cytoplasmic Ca^2+^ signal propagates and is amplified downstream through MLCK activation to control contractility. This is consistent with the previous finding that higher aspect ratio VSMC are more contractile ^29,65^. This observation supports the relationship between cell shape, Ca^2+^ signaling and contractility which has been demonstrated by previous studies^29,30^. Here, we show that geometric factors affect the relationship between upstream signaling and downstream transcription factor activation, providing a physical link between two chemical pathways.

**Figure 6.**
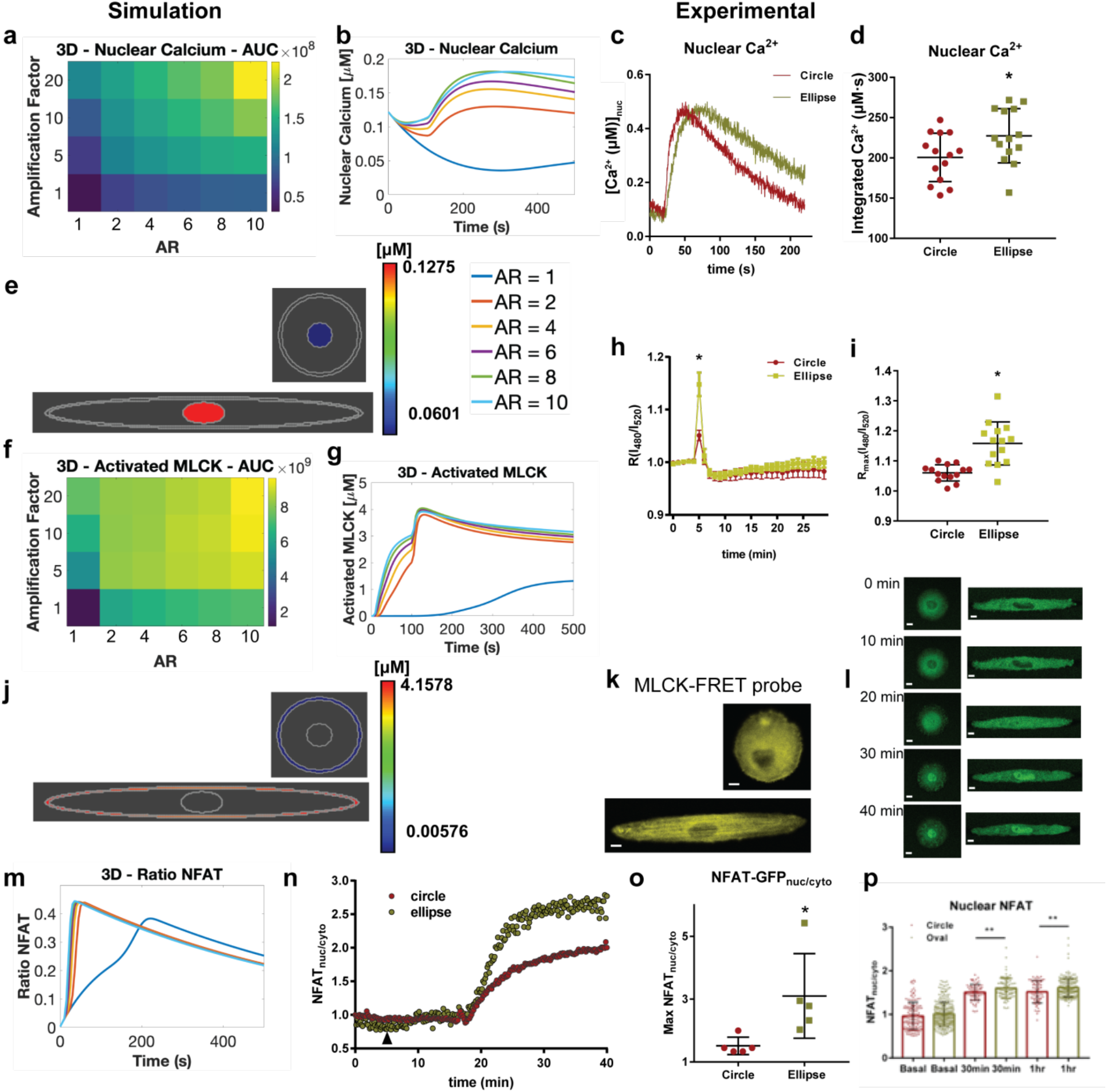
Downstream signaling effects of shape on nuclear Ca^2+^/MLCK/NFAT. (a) Simulation results in *Virtual Cell* for effects of SR amplification factor and AR on nuclear calcium AUC. (b) Temporal dynamics of nuclear calcium for models with various ARs. (c) Average time-course of nuclear Ca^2+^ in response to CCh (d) scatterplot of nuclear integrated Ca^2+^ (area-under-the-curve) in circular and elliptical VSMC. (*P = 0.035, two-tailed t-test). (e) Simulation spatial results of nuclear calcium at t = 125 s for AR = 1 (upper panel) and 10 (lower panel). Right inset: legend for all temporal simulation results (b, g, and l). (f) Effects of SR flux amplification factor and AR on activated MLCK AUC. (g) Temporal dynamics of activated MLCK for various ARs. (h) Temporal MLCK activation in circular and ellipse VSMC upon stimulation with carbachol, activation was measured by decrease in FRET and increase in R480/520 (i) scatterplot of maximal R(I_480_/I_520_) from individual cells expressing MLCK-FRET probe. (*P≤0.0001, two-tailed Mann Whitney test). (j) Spatial results of activated MLCK at t = 125 s for AR = 1 (upper panel) and 10 (lower panel). (k) Circle and elliptical cells expressing MLCK-FRET probe. (l) Representative VSMC complying to circle and elliptical micropatterns expressing NFAT-GFP, in basal and after stimulation with carbachol. (m) Model results for NFAT ratio for different ARs. (n) Representative NFAT-GFP time-course (o) scatterplot of maximal NFAT_nuc/cyt_ ratio in circle and elliptical cells. Max NFAT^circle^_nuc/cyto_=1.51±0.12, *N=5*, Max NFAT^ellipse^_nuc/cyto_=3.10±0.60, *N=5,* p-value=0.0325, two-tailed t-test (*P = 0.0325, two-tailed t-test, N=5 for each group). (p) Single cell quantification of endogenous NFAT translocation (NFAT_nuc/cyt_) in basal and M_3_R stimulated states, in circle and ellipse VSMC. Bar plots shown are mean±SD with corresponding dot plot overlays (at basal, *N_circle_=138, N_ellipse_=284*, at time = 30 minutes, *N*_*circle*_=*72, N*_*oval*_=*81*, **P = 0.024, at time = 1 hour, *N*_*circle*_=*69, N*_*oval*_=*193*, **P = 0.010, two-tailed Mann-Whitney test). Scalebars shown are 10 μm.

Increase in nuclear Ca^2+^ in elliptical cells is likely to impact the nucleo-cytoplasmic transport of NFAT, which exhibits a Ca^2+^/calcineurin dependent translocation and residence time in the nucleus ^19,66^. We predict that increased AR leads to a modest increase in rate of increase of NFAT ratio over 500s (Fig. 6m). We measured NFAT-GFP localization dynamics in live VSMCs in elliptical and circular 3D biochips in response to Gα_q_ activation through M_3_R stimulation (Fig. 6l,n,o,p). Elliptical cells exhibited greater NFAT-GFP nuclear localization compared to circular cells (Fig. 6n) and on average displayed higher maximal NFAT_nuc/cyto_ (Fig. 6o). We independently validated these differences in NFAT localization by immunofluorescent antibody-based visualization of NFAT1 (Fig. 6p, Supplementary Fig. 9a). At basal levels, NFAT_nuc/cyto_ were similar between circular and elliptical cells. However, elliptical cells displayed higher nuclear NFAT compared to circular cells at 30 minutes and 1 hour after stimulation, consistent with live-cell NFAT-GFP translocation results.

To determine whether the effects of Ca^2+^ were general, we studied the localization of SRF in the nucleus. Ca^2+^ also triggers the nuclear localization of SRF through nuclear Ca^2+^/CaMKIV ^20^ and actin dynamics ^67,68^. Elliptical VSMCs show increased nuclear SRF compared to circular cells at both basal and stimulated levels (Supplementary Fig. 9b). In contrast there was no difference in the localization of myocardin, a transcription factor known to be unaffected by Ca^2+^ dynamics, between circular and elliptical cells (Supplementary Fig. 9c). Myocardin has been shown to be constitutively active ^21,69^. Taken together, these results indicate that shape-induced modulation of Ca^2+^ signaling alters Ca^2+^ dependent physiological activities in both the cytoplasm and the nucleus.

### An integrated model of global cell shape and organelle location provides insight into curvature coupling of Ca^2+^ dynamics in VSMCs

We used spatial models constructed in both COMSOL and *Virtual Cell* to study the effects of cell curvature and PM-SR distance on calcium and downstream signaling dynamics. Our experimental observations find a qualitative association between cell aspect ratio and PM-SR distance (Fig. 4). Our model results predict that changing aspect ratio and PM-SR distance will modify signaling dynamics on both short- and long-timescales, and experimental observations validate these predictions (Figs. 5 and 6). We note that this qualitative trend of increased signaling AUC with increasing aspect ratio and increasing amplification factor was seen in both 2D and 3D models (Figs. 5–6 and Supp Fig. 13–15). Therefore, we conclude that cell shape and curvature regulates cell signaling in VSMCs, through changes in PM-SR distance. Briefly, it is important to note that currently there is some debate over the source of increased cytoplasmic calcium seen experimentally in VSMCs, with two competing hypotheses. The first explored in this paper is that cell shape and reduced PM-SR distances affects IP_3_ dynamics and triggers calcium release from the SR. The second is that a reduction in SR calcium triggers SOCE, Store Operated Ca^2+^ Entry, and extracellular Ca^2+^ influx through an interaction of STIM and ORAI channels ^70,71^. In both of these cases, PM-SR distance would play a key role in regulating cytosolic Ca^2+^, whether through IP_3_ dynamics or STIM-ORAI interaction. However, it remains experimentally difficult to differentiate between these two methods of Ca^2+^ increase. Furthermore, in our model, we do not see extreme amounts of SR calcium depletion and SR calcium depletion is believed to be necessary for SOCE (see Fig 3 in ^72^). As we are operating in this regime and due to the continued controversy and lack of experimental evidence, we have omitted SOCE from the model.

## DISCUSSION

One of the key features of intracellular signal flow is the spatial organization of information propagation. This feature is used in multiple cell types to closely regulate the temporal dynamics of second messengers such as calcium and cAMP. For example, in cardiac and skeletal muscle cells, T-tubules, which are tubular invaginations that penetrate into the center of the cell are enriched in L-type calcium channels ^73,74^. Mature dendritic spines in neurons have a specialized ER called the spine apparatus, which is thought to play a role in regulating calcium concentrations in spines ^75^. Here, we bring together several seemingly independent effects of global cell shape to provide an integrated view of how curvature affects organelle location as well as distribution of receptors in the plane of the membrane to modulate signal transduction and thus affect cellular function (Fig. 7). Shape and biochemical signaling are coupled together in a feedback loop to maintain phenotype: cell shape integrates external mechanical and chemical signals on the plasma membrane ^40^ while intracellular signaling cascades containing chemical information in the form of reaction kinetics, in turn regulate cell shape ^76,77^. We propose that shape-dependent endomembrane and nuclear organization serve as the critical link that connect these two components in the feedback loop. This intricate, non-linear coupling of geometric and chemical information can potentially lead to signal amplification to control phenotype.

**Figure 7.**
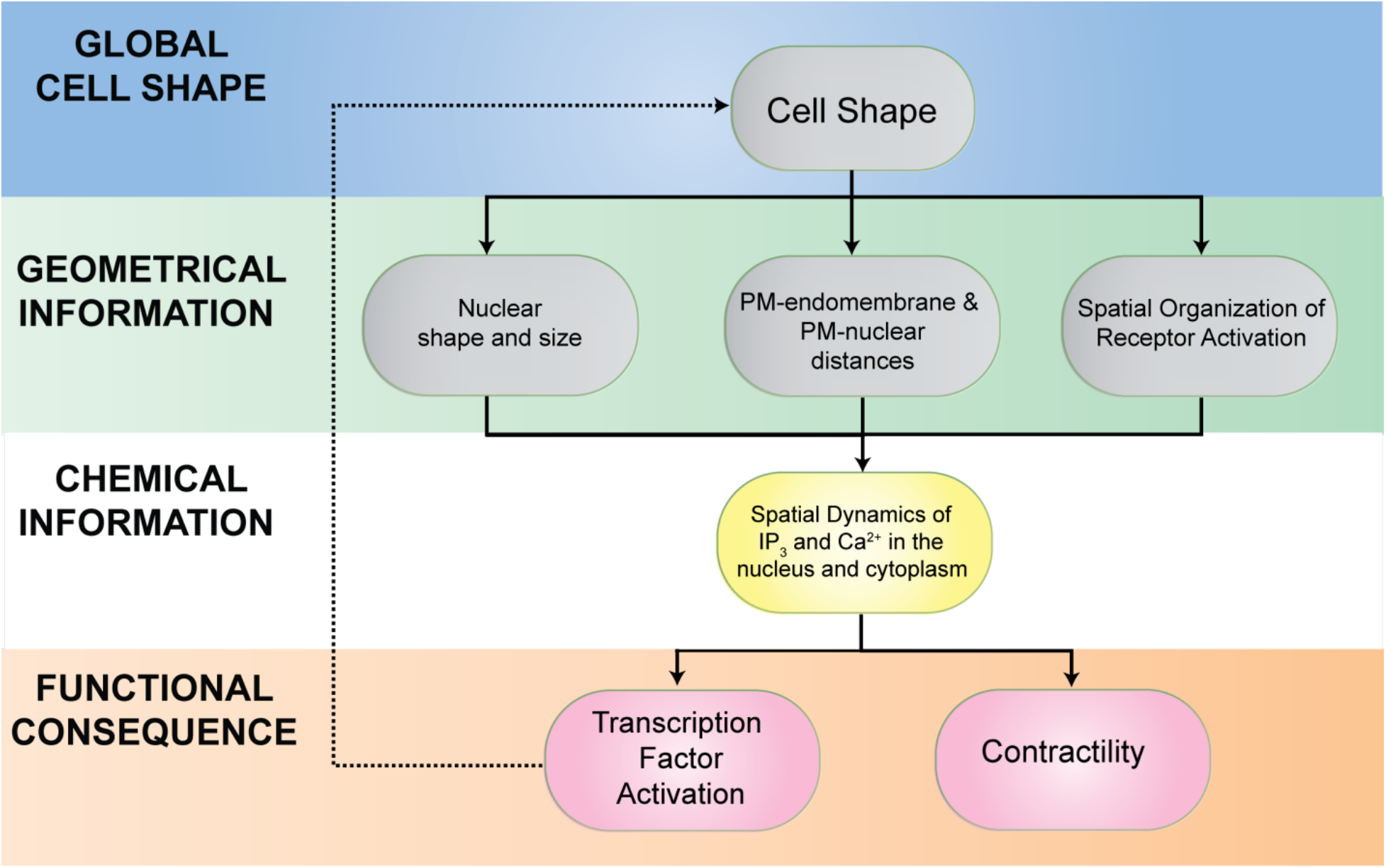
Systems approaches to controlling signaling through subcellular organization hierarchy. Physical determinants such as shape and geometrical information within the cell interact with organelle location and chemical information transfer to impact cellular function and gene expression, which in turn, feedback into cell shape. While each of these effects – shape, organelle location, and chemical information – may separately lead to small changes in signaling, collectively, these effects can lead to altered cellular function including transcription factor activation and contractility.

We used VSMC as a model system to unravel the complex relationship between global cell curvature, signaling, and endomembrane organization. During atherogenesis, disruption of local microenvironment in the medial layer of blood vessels causes VSMC to lose its native spindle shape, and subsequently lose contractile function ^25,78^. It was not clear how loss of shape can lead to a decrease in contractile function. This study provides a mechanistic explanation for the functional role of shape in VSMC contractile function. Cell shape governs membrane curvature, which enables the emergence of systems level properties; cell elongation simultaneously concentrates plasma membrane receptors in the flatter regions of the membrane and reduces the distance between the PM-SR and SR-nucleus in the same region, hence effectively forming a diffusion-restricted domain where receptors, SR, and the nucleus become closer to each other, establishing high effective IP_3_ and Ca^2+^ concentration in the cell center. Given the slow diffusion coefficient of IP_3_ (≤ 10 μm^2^/s) which limits the range of action over which it can exert its signal ^31^ and the dimensions of typical spindle-shaped VSMC (long axis ≥150 μm and short axis length of ≤ 10 μm), control of endomembrane organization by cell shape is a physical, non-genomic mechanism by which IP_3_/Ca^2+^ signals can be locally amplified in order to achieve high concentration of Ca^2+^ globally. These can further regulate contractility through amplification of downstream signaling pathways of MLCK and Ca^2+^-dependent gene expression in the nucleus. Although the effect of cell shape on upstream signaling events and global calcium are small, the progressive amplification during signal flow to physiological effectors through coupled reaction kinetics and spatial organelle organization can produce very different phenotypic effects. This relationship between small changes in levels of second messenger resulting in altered phenotypic response is not unique to calcium signaling pathways. Back in 1994, we had shown that small changes in cAMP levels by activated Gαs could inhibit transformation of NIH-3T3 fibroblasts by oncogenic Ras ^79^.

While we have extensively explored the role of cell shape in signaling here and previously ^3,4^, we have taken an experimental approach using engineered shapes to identify the effects of cell shape by itself. At a conceptual level this is like studying the biochemical activity of a purified protein without the influence of its regulators. In the case of our cell shape studies, features that we have not considered here, such as the forces exerted by the extracellular microenvironment on the cell, and vice versa, also play a critical role in transmitting geometric information ^80,81^, trafficking ^82^, and signal transduction ^83^. Furthermore, the interaction between signaling and cytoskeletal remodeling can lead to changes in cell shape and local curvature ^84,85^. Our observations of increasing anisotropy and robustness in expression of actin myofibrils, along with increased nuclear SRF localization with aspect ratio indicate that cytoskeletal signaling also contributes towards the contractile phenotype of VSMC. Cytoskeletal and Ca^2+^ signaling may act in concert in maintaining the differentiated phenotype of spindle-shaped VSMC. We have able to uncover unique aspects of signal flow regulation in cells based on geometry (cell shape) and chemical reaction cascades alone, and coupling these effects with the effects of extracellular microenvironment and the role of intracellular cytoskeletal interactions is a focus of future studies. We conclude that being at the right place at the right time is critical for information to flow from one cellular compartment to another and for short term signals like Ca^2+^ to have long lasting effects such as contractility and transcription factor activity.

## AUTHOR CONTRIBUTIONS

R.C.C conducted the experiments, data analysis, and initial VCell simulations; M.K.B performed VCell simulations and model development; A. R. and W.G.M.J conducted the experiments; M.H. and S.B. designed and manufactured the biochips; E.U.A. designed and supervised the 3D EM experiments, which were performed by G.P., P.P. and N.J.W..; J.H. oversaw the design of the biochips; S.S. aided with experiment design and data analysis; P.R. worked on model development and 3D simulations; R.I. oversaw the concept, design of the study. All authors worked on writing and revising the drafts of the manuscripts. All authors concur with the contents of the manuscript.

## ACKNOWLEDGEMENTS

We thank Dr. Eric Sobie, Dr. Marc Birtwistle, and Dr. Michael Getz for discussion and reagents. This work was supported by NIH Grants GM072853 and the Systems Biology Center grant P50GM071558. R.C.C. was supported by a NIH postdoctoral fellowship F32GM116415 and research in P.R.’s laboratory is supported by Air Force Office of Scientific Research (AFOSR) Multidisciplinary University Research Initiative (MURI) grant FA9550-18-1-0051. M.B. was supported by the National Defense Science and Engineering Graduate (NDSEG) Fellowship. E.U.A. was supported in part by NIH Grants DK118222 and DK124917.

## SUPPLEMENTARY INFORMATION

### NUMERICAL SOLUTIONS FOR REACTION DIFFUSION EQUATIONS WITH ROBIN BOUNDARY CONDITIONS

We investigate the dynamics of a molecule C_A_, which is formed at the PM, freely diffuses in the cytoplasm where it can be degraded, and is consumed at the SR membrane. This phenomenon can be represented by the following reaction-diffusion equation with the accompanying boundary conditions.

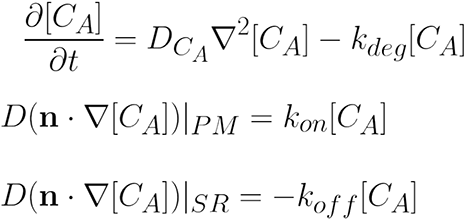

One of the features of this system of equations is that the boundary conditions at both ends are time-dependent and belong to a class of boundary conditions known as Robin boundary conditions ^1^. Furthermore, the boundary conditions depend on the local shape, since the surface normal will change as a function of curvature^2^. If the boundary conditions at both membranes were constant values of the concentration of A, the solution to the partial differential equation (Eq. 1) is a linear distribution from one boundary to the other^1^. However, in the current case, where both boundary conditions have Robin boundary conditions, the solution to these equations depends *both* on the diffusion distance and the reaction rates. The Robin boundary conditions that accompany the reaction-diffusion equation are of the mixed-type and are time dependent. Therefore, writing analytical solutions for them is challenging. Therefore, we use numerical methods to solve this equation. We use finite-element methods to solve the equations in the commercially available software COMSOL Multiphysics®.

Briefly, we used the General Form PDE interface to model the partial differential equations and the flux boundary conditions implemented using the Flux mode with the source term set to the reaction rate. We used six different geometries to test the role of curvature and PM-SR distance. The mesh size was set to ‘Extremely Fine’ and the tolerance was set to relative tolerance was set to 0.01. For the toy model displayed in Fig. 1b, we used parameter values of k_deg_ = 0.5 [1/s], k_on_ = k_off_ = 0.75 [μm/s)], D_Ca_ = 1 [μm^2^/s], and Ca_IC_ = 5 [μM]. Area was conserved across all shapes. Rectangles had heights and widths of 150 nm and 500 nm, 100 nm and 750 nm, and 50 nm and 1500 nm. The circle section had an inner radius of 500 nm, a width of 150 nm, and a sector angle of 50 degrees. The constant elliptical section had a width of 100 nm and sector angle of 73 degrees. The tapering elliptical section had a starting width of 132 nm, an ending height of 25 nm, and a sector angle of 90 degrees. The plot in Fig. 1b is at 50 ms and has a range of values from ~4.7 to 5.9 μM. This toy model served to demonstrate the intricate relationship between curvature and PM-SR distance.

For a rectangle and a circle, solving the PDE with the boundary conditions shown above amounts to solving a 1-dimensional problem (either in the z-direction or in the radial direction). As a result, a gradient is observed only in one direction. For an elliptical cross section (Fig. 1A), a triangle (Supplementary Fig. 1A) and a trapezium (Supplementary Fig. 1B), the geometric variation is in two-dimensions. Therefore, the spatio-temporal dynamics of C_A_ is different in these patterns. Furthermore, in an ellipse, the curvature is varying along the membrane, and therefore, the diffusive flux also varies. This is because the diffusive flux is dependent on the local normal and curvature captures the rate of change of the normal along the curve^2^.

#### Dimensional analysis

Using the PM-SR distance ‘L’ as a characteristic length scale and k_deg_ as the characteristic time scale, we non-dimensionalized the partial differential equation for C_A_ to obtain a dimensionless number (k_deg_L^2^)/D which is the Damkohler number^1^. When the Damkohler number is much greater than 1, then the system is diffusion-dominated and when the Damkohler number is much less than 1, it is reaction-dominated. In cellular environments, various protein interactions and obstacles can lead to diffusion-trapping, which slows diffusion to an effective diffusion given by 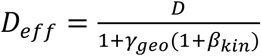, where *γ*_*geo*_ represents the geometric effects of traps and *β*_*kin*_ represents kinetics of binding at traps if applicable ^3,4^. Replacing D in the Damkohler number with this D_eff_ captures the effects of geometric and kinetic barriers to diffusion that can alter signaling dynamics.

#### IP_3_ model implementation in COMSOL

The dynamics of IP_3_ were modeled using the reaction network model presented in Cooling et al.^5^ and implemented in COMSOL using the protocol presented in Vollmer et al.^6^. We modeled multiple compartments – the extracellular space, PM, SR, SR membrane, the nucleus and the nuclear membrane. The various reactions can be found in Supplemental Table 1 from R1-R16. The Robin boundary conditions were implemented using the Flux boundary condition node for the volume and the general boundary form PDE node for the membranes. The mesh size was set to ‘Extremely fine’ such that the minimum element size is 0.1 nm and relative tolerance was set to 0.01. These values of mesh size and tolerance were chosen such that smaller values of tolerance or mesh refinement didn’t alter the numerical results.

### INTEGRATIVE WHOLE-CELL REACTION-DIFFUSION MODEL

To understand how global cell shape can modulate the dynamics of IP_3_ and calcium signaling, we developed mathematical models which analyze the relationships between the binding of extracellular ligand to GPCR, G-protein activation and cycling, PLCβ activation, IP_3_ activation of IP_3_R in the SR, intracellular calcium dynamics, MLCK activation, and NFAT dynamics. Reactions were initially implemented as a system of ordinary differential equation in the *Virtual Cell* environment (http://www.nrcam.uchc.edu/)^7^, and then modeled as a system of partial differential equations (PDEs) in two-dimensions, and then implemented in three dimensions. The complete list of model equations, parameters, units and references of the kinetic parameters are listed in Table 1 and the complete list of initial conditions and diffusion coefficients are listed in Table 2.

#### GPCR Signaling-PLCβ cycling

We adapted the GPCR signaling modules described by Cooling et al^5^ and Eungdamrong et al^8^ with modifications. Reactions R1-R6 represent GPCR-Ligand-G protein interactions on the plasma membrane. The reactions are modeled as kinetic fluxes following the mass action and 9Michaelis-Menten kinetics (see Bhalla and Iyengar). A ligand stimulus of 10 μM was applied from 100s to 150s. The binding of extracellular ligand to cell-surface receptors are represented by reactions R1 and R4. Binding of the ligand to muscarinic acetylcholine receptors causes a conformational change on the Gα_q_ subunit of the heterotrimeric G-protein, causing replacement of GDP with GTP (R5) and dissociation of the Gα subunit from the heterotrimeric G-protein. The binding of activated Gα_q_ to the enzyme PLCβ are represented by R8 and R10. Gα_q_-GTP is hydrolyzed to Gα_q_-GDP by autocatalysis represented by R7. Reactions R14-R16 represent IP3 dynamics, including its production at the plasma membrane, membrane diffusion and degradation at the cytosol. R14 and R15 shows the hydrolysis of PIP_2_ to IP_3_ by PLCβ-Ca^2+^ and PLCβ-Ca^2+^-Gα_q_(GTP) respectively. The degradation of IP_3_, which involves the dephosphorylation and phosphorylation of IP3 to form inositol biphosphate and inositol tetraphosphate are represented by R16. The metabolic recycling of IP_3_ from these products is not considered. PLCβ inactivation is represented by R13.

#### IP_3_/Calcium Dynamics

In vascular smooth muscle cells, agonist-induced calcium release from the SR/ER is the main source of intracellular Ca^2+^ transient. The opening of the inositol triphosphate receptor channels (IP_3_R) from the ER membrane, leads to an increase in cytoplasmic calcium. We used the simplified version of channel kinetics ^10,11^ by De Young and Keizer ^12^ which is represented by R17. Calcium leak from the ER to the cytosol is represented by R20. Calcium is pumped back to the ER by the sarcoplasmic reticulum calcium ATPase (SERCA) pump, represented by R21. Calcium is also pumped into the nucleus through the NPC, represented by R41. We also model downstream signaling dependent on cytosolic calcium including activation of Calmodulin (CaM), Calcineurin (CaN), CaMKII, MLCK, and NFAT (R22-40).

#### Geometries

The spatial geometries in *Virtual Cell* of the cell and organelles were depicted as idealized geometries represented as a series of concentric ellipsoids that represent the whole cell, SR and the nucleus. We assumed that cells complying with shapes of increasing aspect ratios (AR) conserve the cell volume and increase the plasma membrane surface area with increasing AR. Hence, whole cell, cytoplasmic and SR areas were kept constant with increasing aspect ratios for 2D simulations, while whole cell, cytoplasmic and SR volumes were kept constant with increasing aspect ratios for 3D simulations. The cellular geometries were approximated from experimentally observed VSMC confined from AR 1:1 to AR 1:8 (Figure 3a), whereby the SR is closer to the PM in the perinuclear region compared to the cell tips. Furthermore, as the aspect ratio increased, both PM-SR and PM-nuclear distances decreased in the minor axis and increased in the major axis of the ellipse. Nuclear shapes were based on experimentally observed geometries (Fig. 3c-e). Supplementary Fig. 7 show the resulting 3D and 2D geometries, respectively. To solve the PDEs, geometries were discretized into 0.5 μm × 0.5 μm spatial steps. The systems of equations were solved within the Virtual Cell framework using fully-implicit, finite volume, with variable time step solver, with a maximum time step of 0.1 s and an output interval of 5.0 second.

We note that different geometric assumptions were made in simulations for *Virtual Cell* versus COMSOL. COMSOL simulations included disc geometries for the SR that layered on top of each other next to the PM. *Virtual Cell* involved concentric ellipsoids where the SR was modeled as sectors of ellipsoids based on experimental imaging. We note however that real SR structures can be highly complex with layers and folding, which cannot yet be captured fully with either framework (see ^28^). The *Virtual Cell* framework was not able to resolve small distances between PM and SR such as 50-100 nm. This is why we used COMSOL for studying the small length scale effects. Therefore, the COMSOL model was used to implement a simplified biochemical study, but with additional geometric complexity through the layered SR disc stacks.

#### Amplification factors for the SR

Idealized geometries were used for the cell body and internal organelles. However, it is well-known that the SR in particular has complex structures with large surface area. To account for this large area and to investigate its role in regulating the IP_3_-mediated calcium flux at the ER membrane, an amplification factor was included in the membrane flux of IP_3_ activated calcium release from the SR, R17, and effectively scales the magnitude of R17. By varying this parameter, we investigated the effect of different surface areas of the SR in the model, without explicitly simulating the model equations in these complex geometries.

#### Parametric Sensitivity Analysis

A parametric sensitivity analysis for cytoplasmic calcium and activated MLCK was carried out with respect to all model parameters and initial conditions (Supplementary Figs. 18–21). The analysis was carried out in the software COPASI (www.copasi.org) where the model was implemented as a system of ordinary differential equations.

#### Area under the curve Analysis

We used several methods to convey our spatiotemporal signaling results including temporal plots, spatial images, and Area under the Curve (AUC) analysis. AUC is a commonly used measurement of signaling over time. In computational spatial models, it is typically computed by integrating concentration over the whole geometry at each time point, then integrating the temporal plot of moles (or molecules) over time, producing an AUC with units of moles*sec for example. Experimental results usually integrate temporal concentration over time, producing an AUC with units of μM*sec. As AUC is an additive measure, that is it accumulates the signal over time, AUC will always be increasing or constant.

**Supplementary Table 1.**
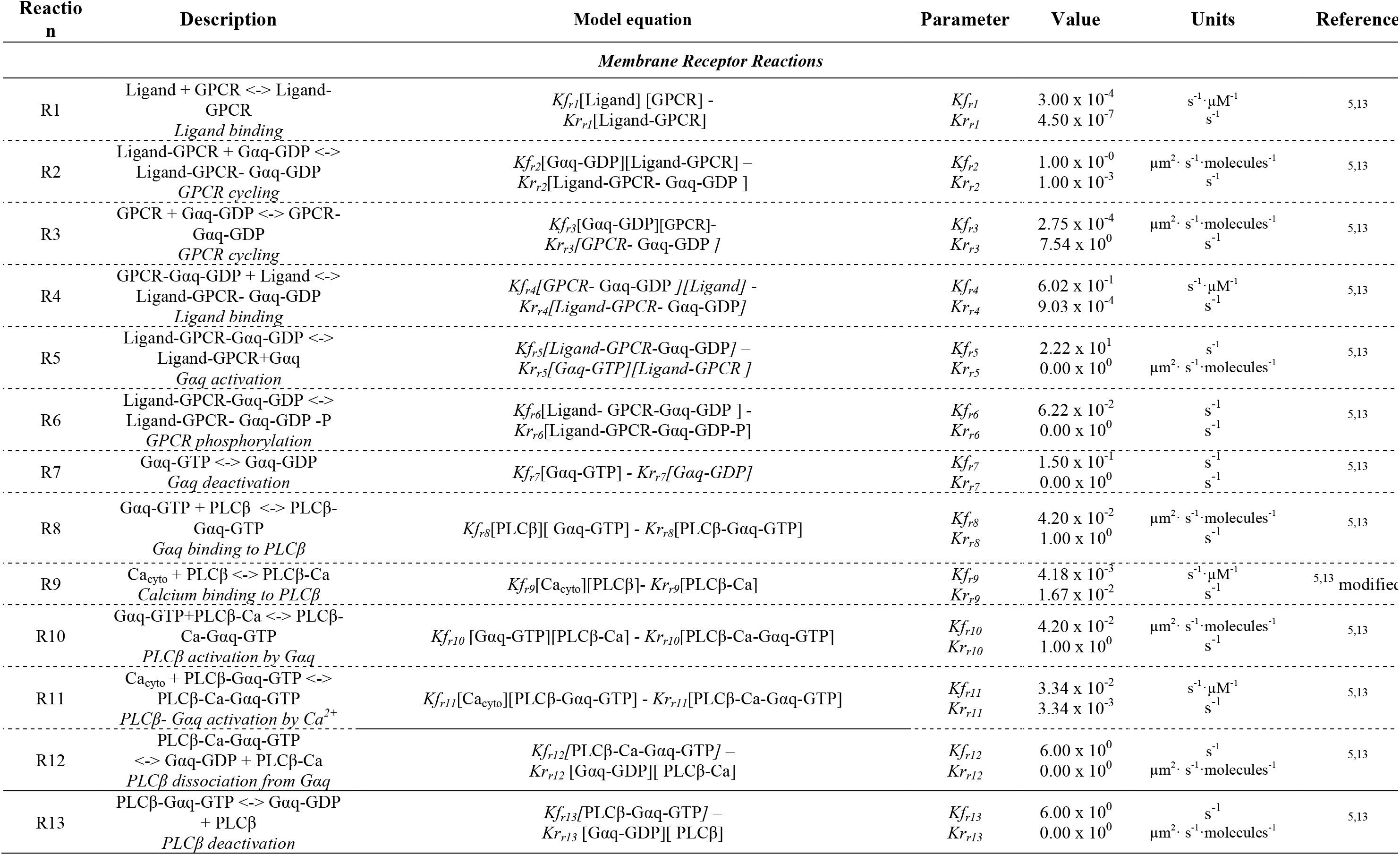
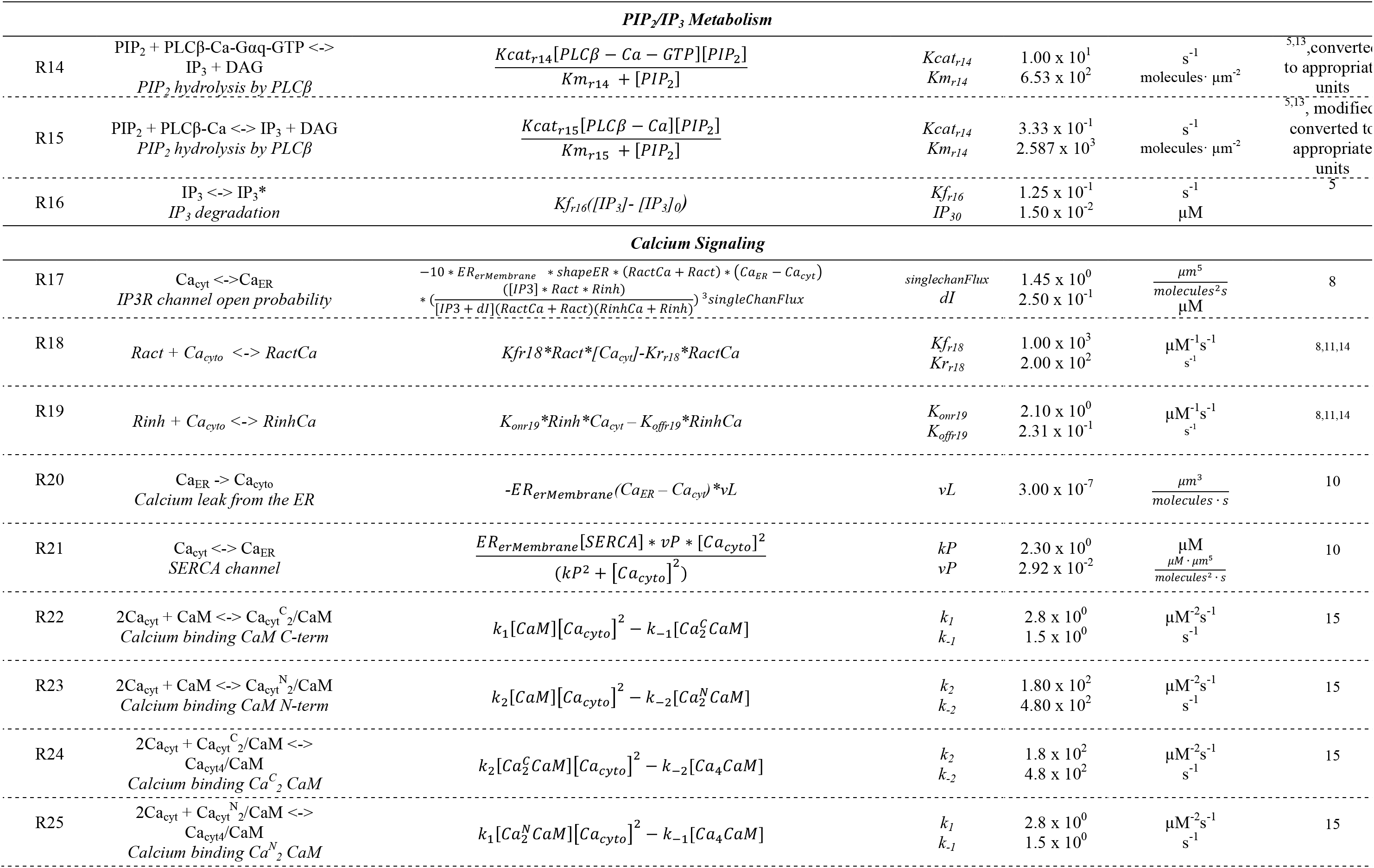
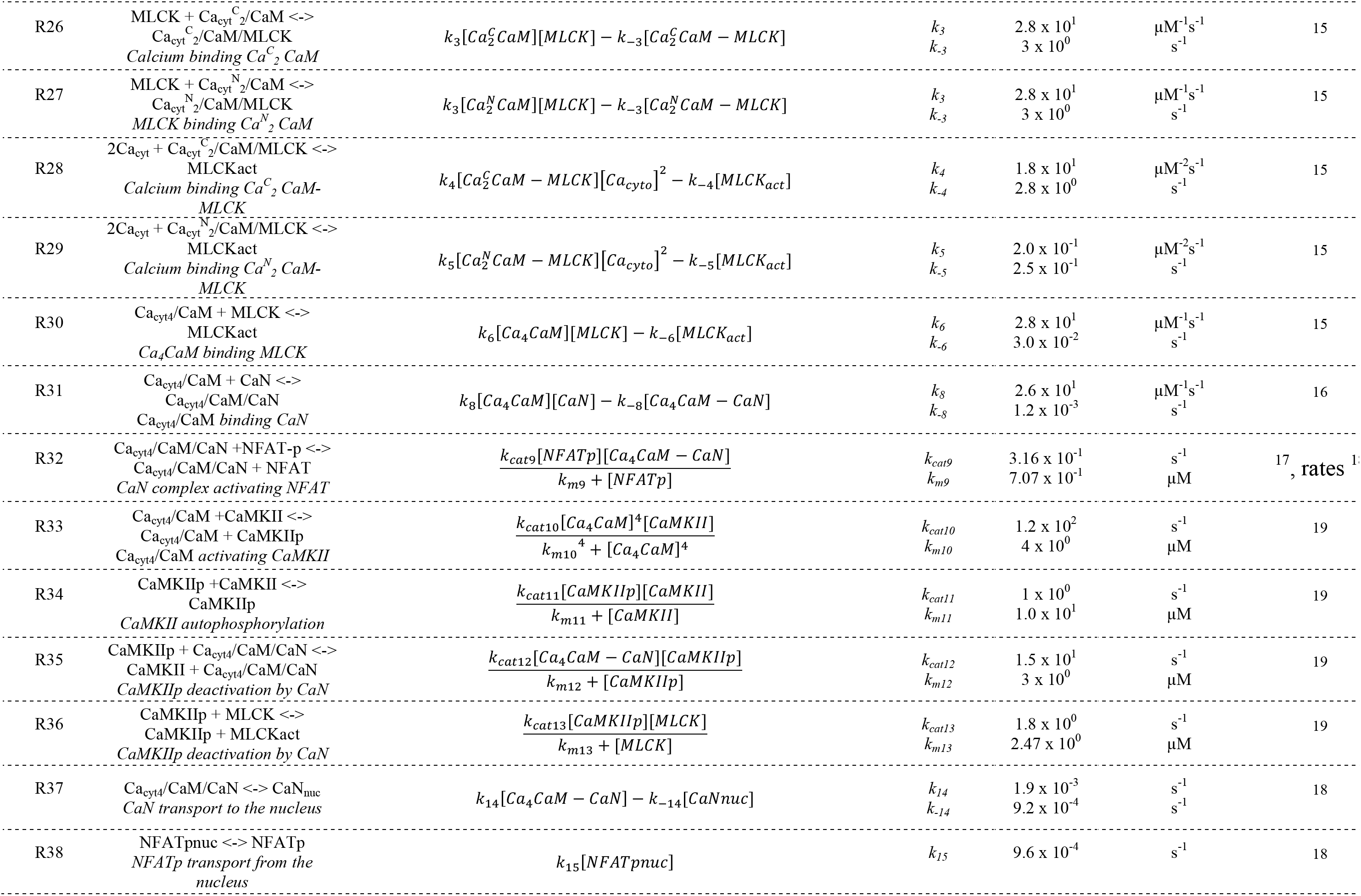
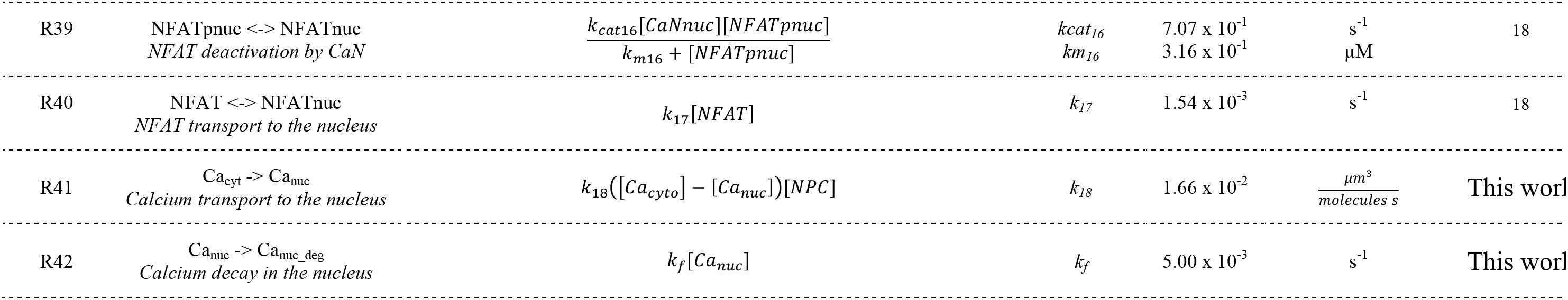
Model reactions and parameter values.

**Supplementary Table 2.** Initial Concentration and Diffusion coefficient of reactants.

| Species | Initial Concentration ($\mu\text{M}$) | Diffusion Coefficient ($\mu\text{m}^2/\text{s}$) | References |
| --- | --- | --- | --- |
| Calcium | 0.1 $\mu\text{M}$ | 220 | Knot, Cooling<sup>5,20</sup> |
| IP3 | 0.015 $\mu\text{M}$ | 10 | Cooling, Dickinson<sup>5,21</sup> |
| Ligand | 10 $\mu\text{M}$ | - | Agonist added |
| Calcium_nuc | 0.122 $\mu\text{M}$ | 220 | Value obtained after equilibration |
| Calcium_ER | 400 $\mu\text{M}$ | 220 | Fink<sup>11,14</sup> |
| Receptor | 12 $\mu\text{m}^{-2}$ *varied in spatial | 0.1 | Cooling, Bers<sup>5,22</sup> |
| L_R_GD | 0 $\mu\text{m}^{-2}$ | 0.1 | Unstimulated state |
| R_GD | 1.07 $\mu\text{m}^{-2}$ | 0.1 | Cooling, Bers<sup>5,22</sup> |
| GT | 0 $\mu\text{m}^{-2}$ | 0.1 | Unstimulated state |
| GD | 10000 $\mu\text{m}^{-2}$ | 0.1 | Cooling, Lukas<sup>5,13</sup> |
| PLC | 90.9 $\mu\text{m}^{-2}$ | 0.1 | Cooling, Lukas<sup>5,13</sup> |
| PLC_GT | 0 $\mu\text{m}^{-2}$ | 0.1 | Unstimulated state |
| PIP2 | 4000 $\mu\text{m}^{-2}$ | 2.5 | Cooling, Xu, Golebiewska<sup>5,23,24</sup> |
| DAG | 0.015 $\mu\text{M}$ | 237 | This work |
| LR | 0 $\mu\text{m}^{-2}$ | 0.1 | Unstimulated state |
| PLC_Ca | 9.09 $\mu\text{m}^{-2}$ | 0.1 | Cooling, Lukas<sup>5,13</sup> |
| PLC_Ca_GT | 0 $\mu\text{m}^{-2}$ | 0.1 | Unstimulated state |
| L_R_GD_P | 0 $\mu\text{m}^{-2}$ | 0.1 | Unstimulated state |
| Rinh | 7.7825 $\mu\text{m}^{-2}$ | 0.1 | Fink, Eungdamrong<sup>8,11,14</sup> |
| Ract | 9.056 $\mu\text{m}^{-2}$ | 0.1 | Fink, Eungdamrong<sup>8,11,14</sup> |
| RinhCa | 3.5375 $\mu\text{m}^{-2}$ | 0.1 | Fink, Eungdamrong<sup>8,11,14</sup> |
| RactCa | 2.264 $\mu\text{m}^{-2}$ | 0.1 | Fink, Eungdamrong<sup>8,11,14</sup> |
| SERCA | 45 $\mu\text{m}^{-2}$ | 0.1 | Fink, Eungdamrong<sup>8,11,14</sup> |
| ER<sub>er</sub>Membrane | 2 $\mu\text{m}^{-2}$ | -- | Fink, Eungdamrong<sup>8,11,14</sup> |
| CaM | 6 $\mu\text{M}$ | 10 | <sup>17,25</sup> |
| CaN | $9.853 \times 10^{-1} \mu\text{M}$ | 10 | <sup>18</sup>, this work |
| CaN* | $1.413 \times 10^{-2} \mu\text{M}$ | 10 | <sup>18</sup>, this work |
| CaN*<sub>nuc</sub> | $2.918 \times 10^{-2} \mu\text{M}$ | 10 | <sup>18</sup>, this work |
| Ca<sub>2</sub>CaM<br>(Ca<sub>2n</sub>CaM+Ca<sub>2c</sub>CaM) | $7.012 \times 10^{-3} \mu\text{M}$ | 10 | <sup>18, 25</sup> |
| Ca<sub>4</sub>CaM | $4.05 \times 10^{-7} \mu\text{M}$ | 10 | <sup>18, 25</sup> |
| CaMKII | 1 $\mu\text{M}$ | 7 | This work, <sup>25</sup> |
| Ca<sub>2</sub>CaMMLCK | 0 $\mu\text{M}$ | 0.05 | <sup>26</sup> |
| MLCK | 5 $\mu\text{M}$ | 0.05 | <sup>15,26</sup> |
| NFATp | $1.617 \times 10^{-2} \mu\text{M}$ | 0.1 | <sup>18</sup>, based on <sup>27</sup> |
| NFAT | $1.17 \times 10^{-4} \mu\text{M}$ | 0.1 | <sup>18</sup>, based on <sup>27</sup> |
| NFATnuc | $3.542 \times 10^{-2} \mu\text{M}$ | 0.1 | <sup>18, 27</sup> |
| NFATpnuc | $1.88 \times 10^{-4} \mu\text{M}$ | 0.1 | <sup>18, 27</sup> |
| NPC | 1 $\mu\text{m}^{-2}$ | 1 | This work |

**Supplementary Figure 1.**
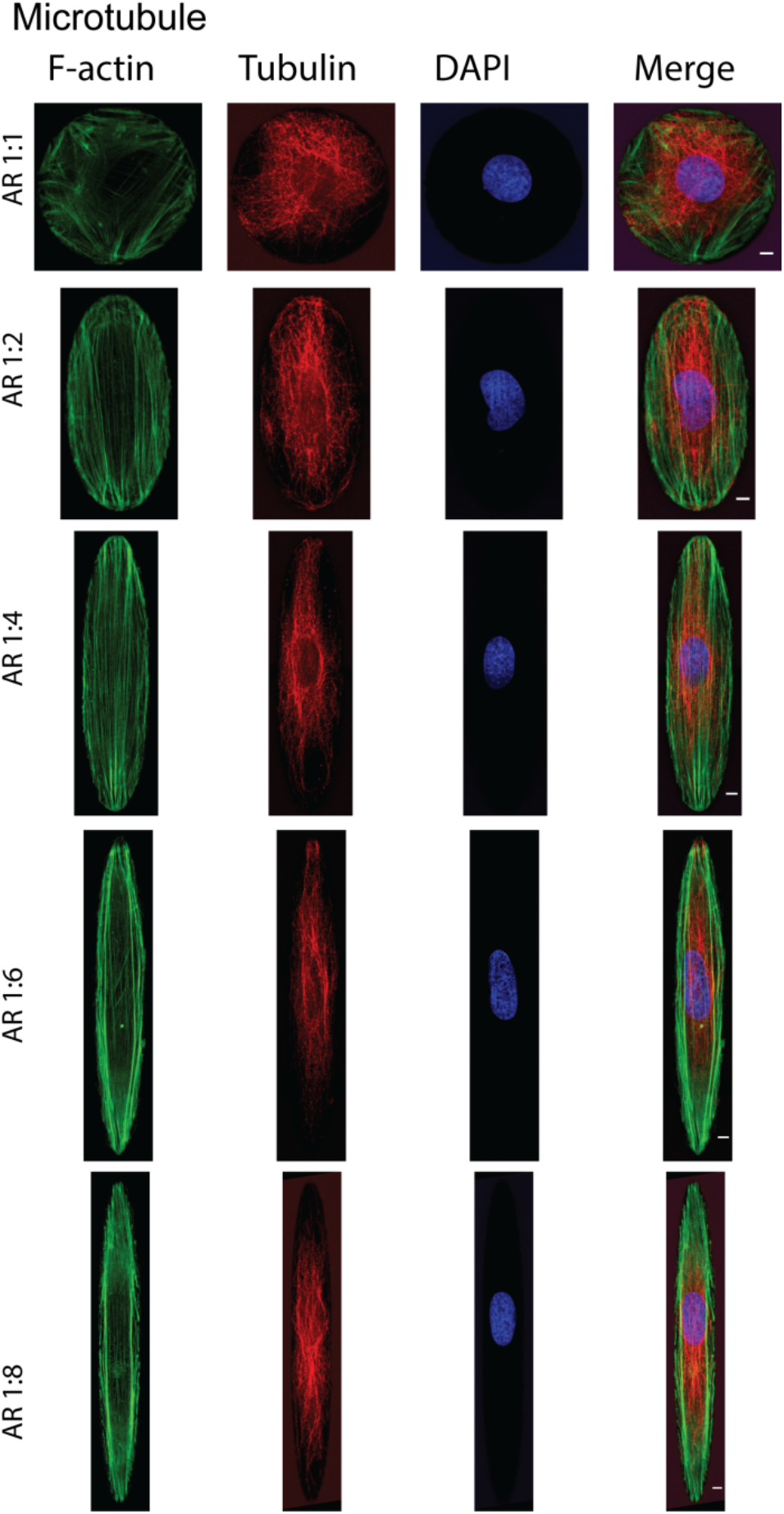
Microtubule organization in VSMC with AR 1:1 to AR 1:8. Shown are staining of F-actin (green), alpha-tubulin (red) and DAPI in VSMC complying with AR 1:8. Scale bars shown in the merged channel are 10 μm.

**Supplementary Figure 2.**
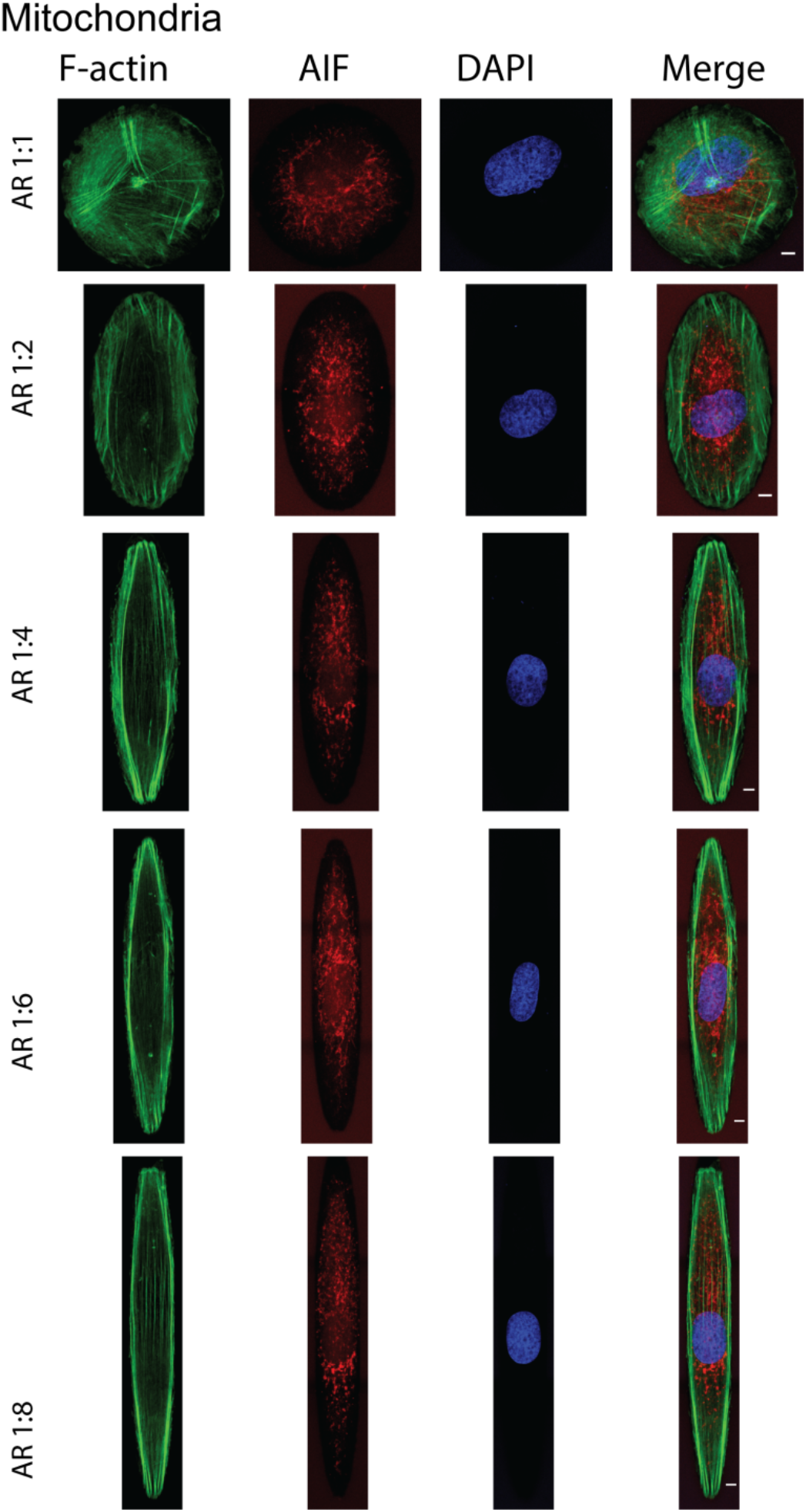
Mitochondrial organization in VSMC with AR 1:1 to AR 1:8. Shown are staining of F-actin (green), mitochondrial marker Apoptosis Inducing Factor (AIF) (red) and DAPI in VSMC complying with AR 1:8. Scale bars shown in the merged channel are 10 μm.

**Supplementary Figure 3.**
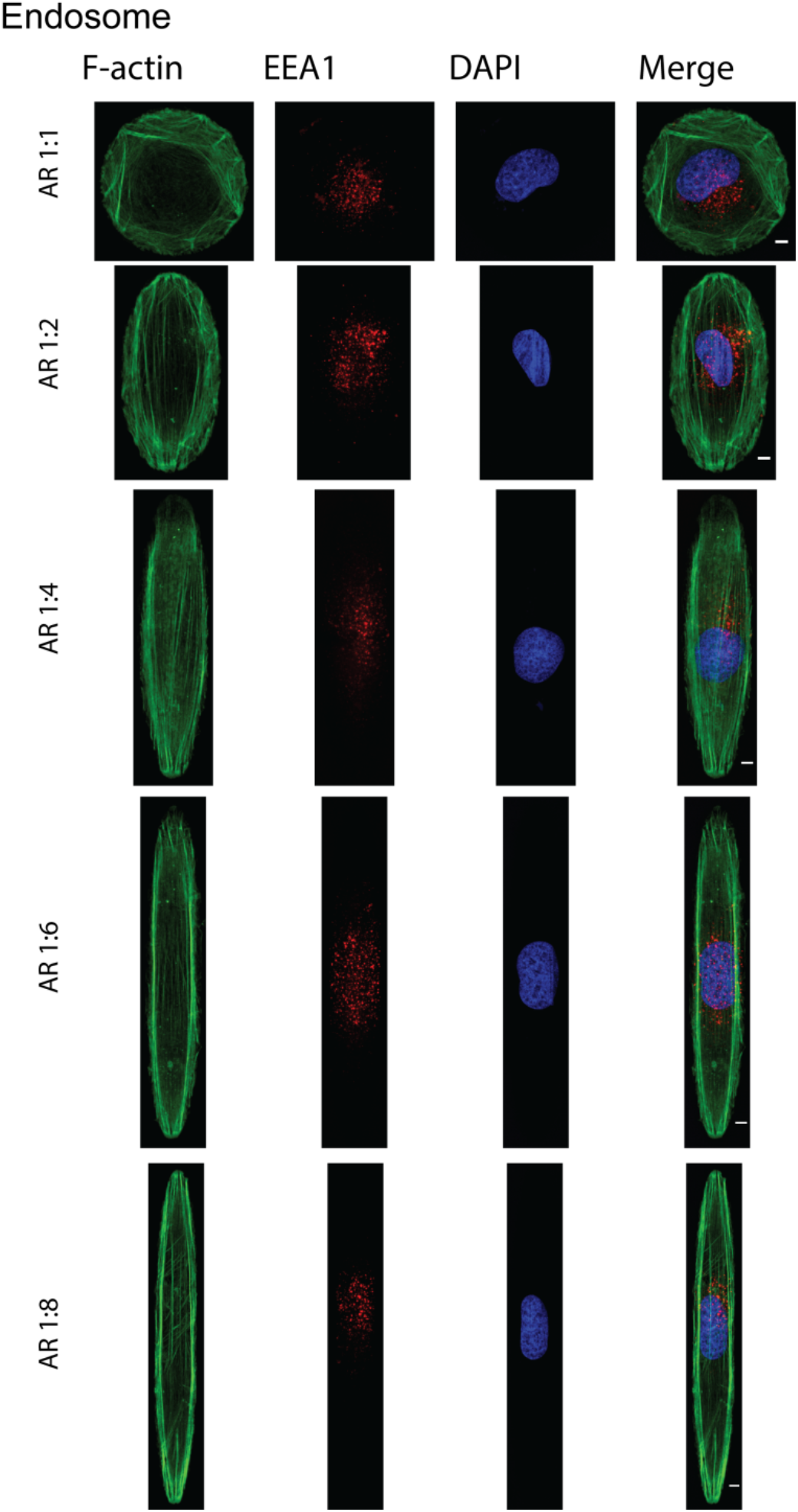
Early endosome organization in VSMC with AR 1:1 to AR 1:8. Shown are staining of F-actin (green), endosomal marker Early Endosome Antigen 1 (EEA) (red) and DAPI in VSMC complying with AR 1:8. Scale bars shown in the merged channel are 10 μm.

**Supplementary Figure 4.**
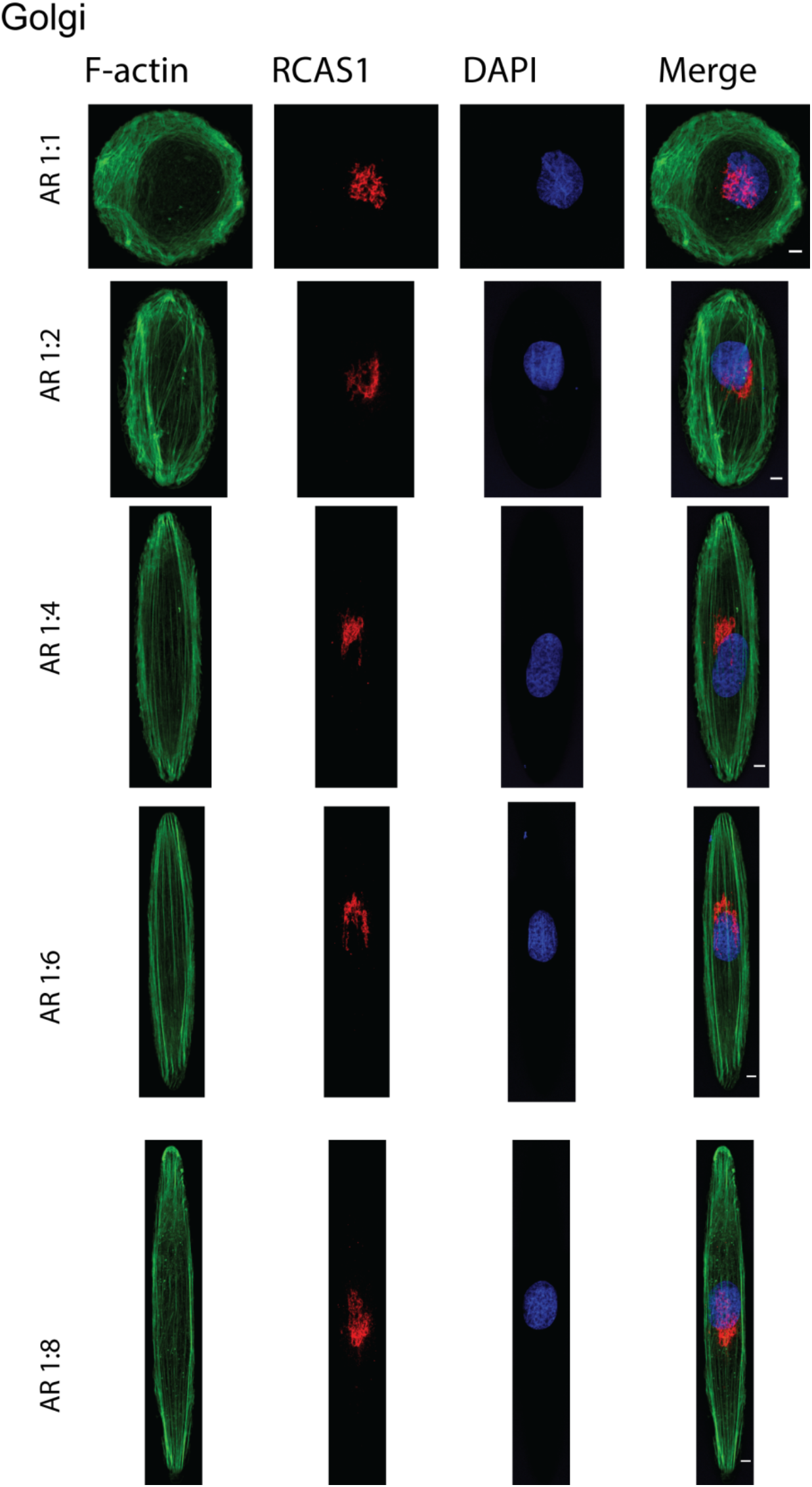
Golgi organization in VSMC with AR 1:1 to AR 1:8. Shown are staining of F-actin (green), Golgi marker, receptor-binding cancer antigen expressed on SiSo cells (RCAS1) (red) and DAPI in VSMC complying with AR 1:8. Scale bars shown in the merged channel are 10 μm.

**Supplementary Figure 5.**
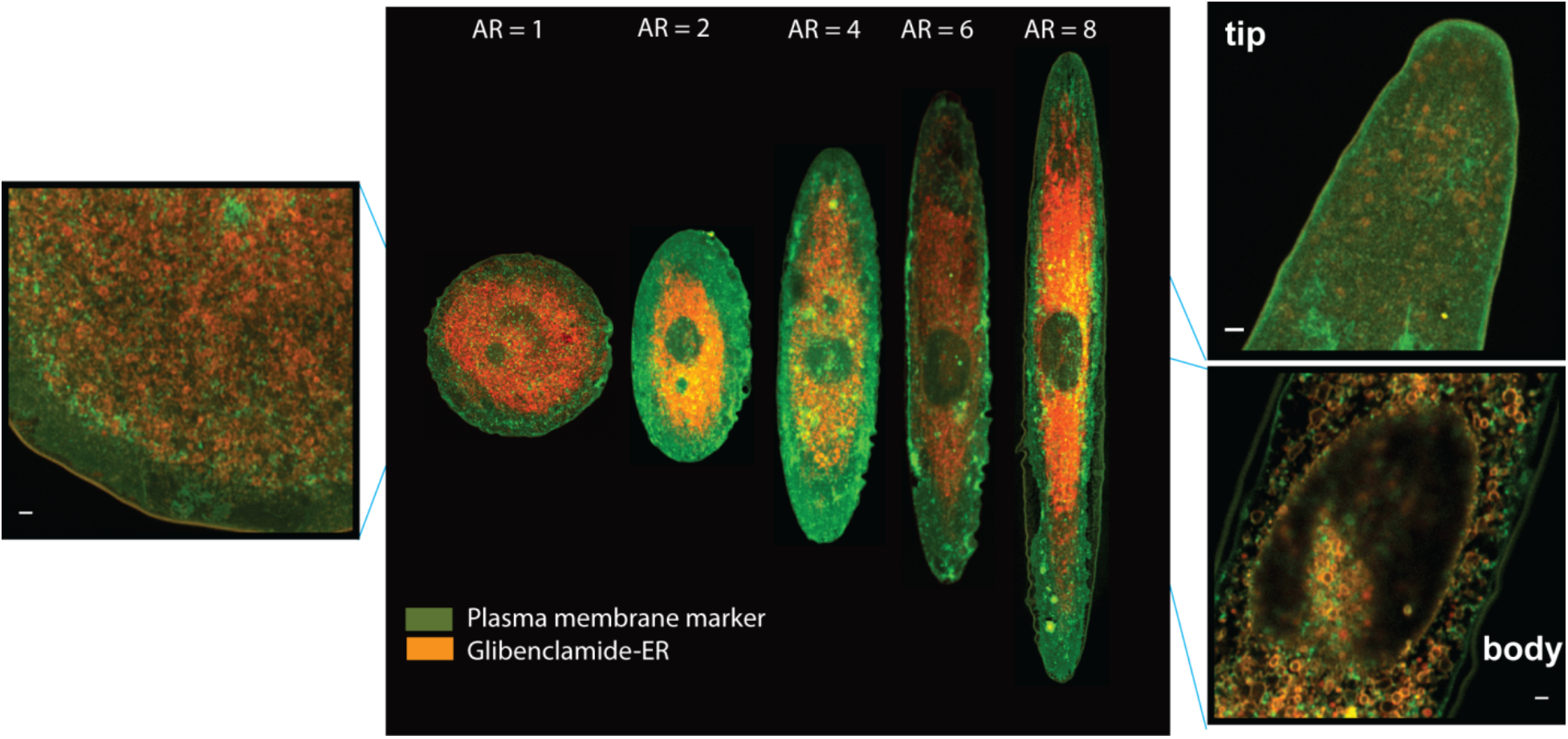
Live cell Airy Scan imaging of SR membrane in VSMC with AR 1:1 to AR 1:8. Shown are confocal and Airy scan images (inset) of plasma membrane marker (CellMask Plasma Membrane marker, Invitrogen), green and SR marker (Bodipy-Glibenclamide ER) in red. Scale bars are 0.5 μm.

**Supplementary Figure 6.**
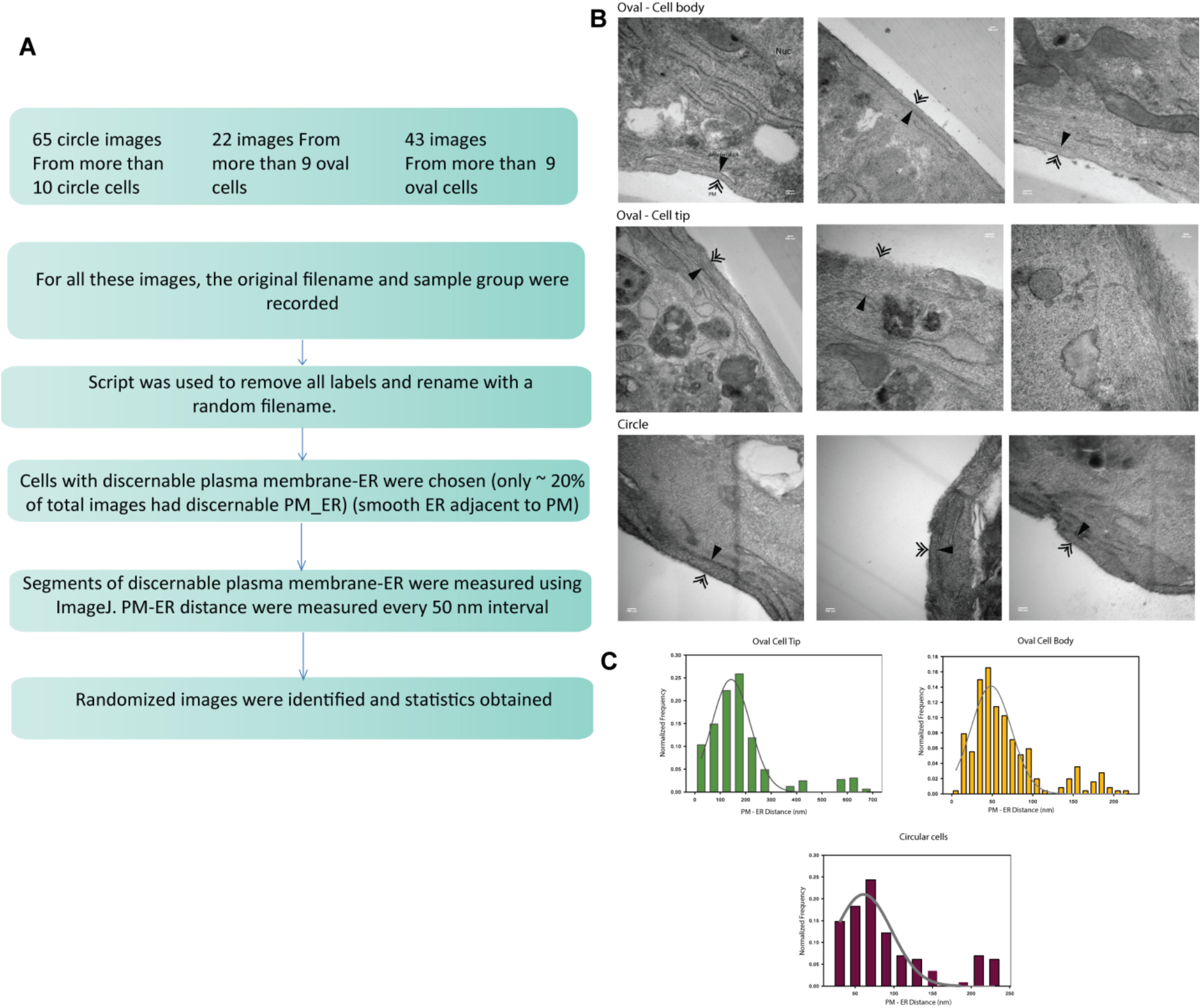
TEM quantification of PM-SR distances from electron micrographs. Peripheral SR was identified by its close apposition to the plasma membrane and the absence of attached ribosomes. PM-peripheral SR was measured at 50-nm intervals. A) Post-imaging processing of EM micrographs to quantify PM-SR distances B) Representative images of cells from elliptical cell body, elliptical cell tip and circular cell tip. C) Histograms of distances obtained. Scale bars shown are 50 nm.

**Supplementary Figure 7.**
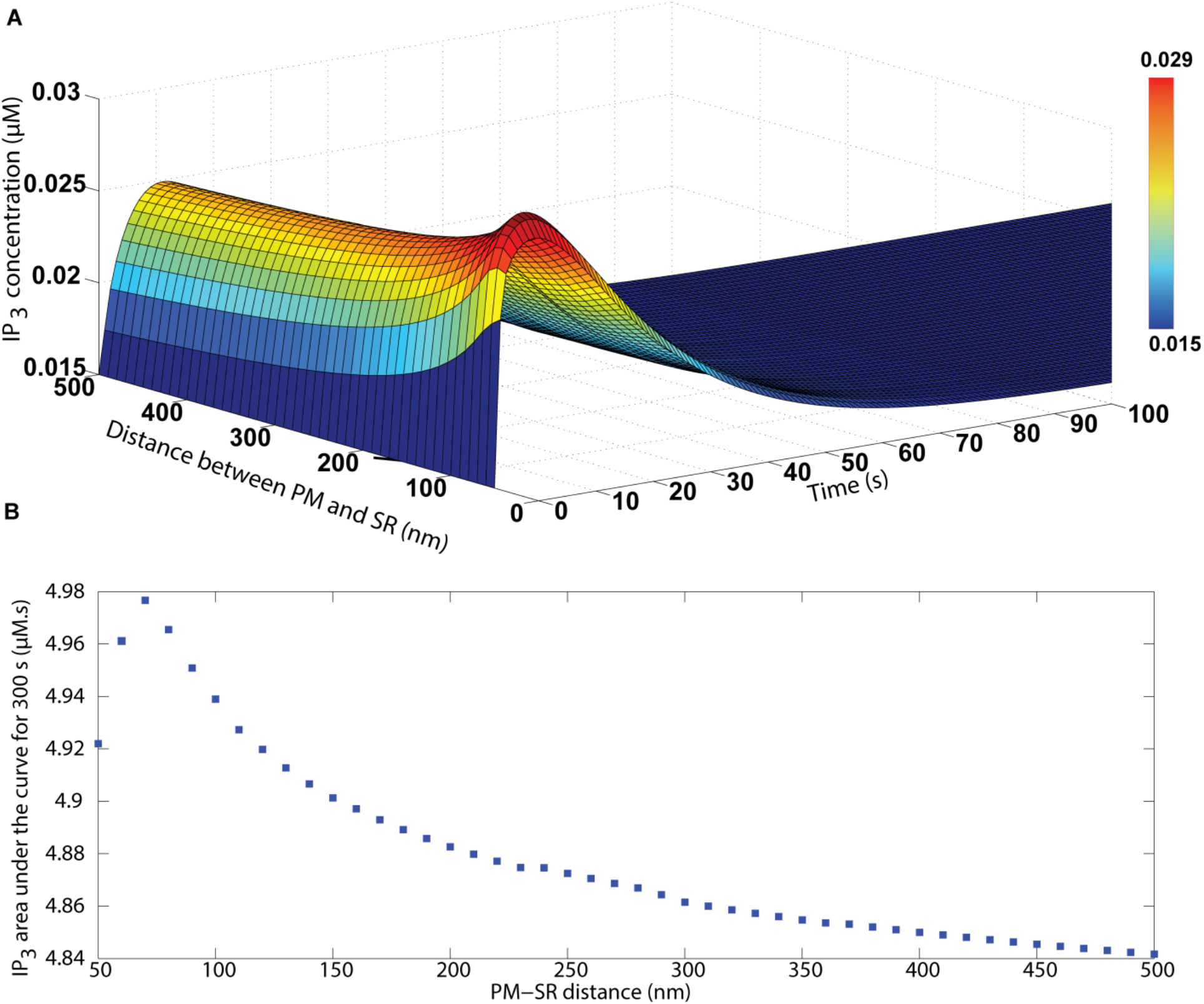
The spatio-temporal profile of IP_3_ at different PM-SR distances and resulting AUC from the COMSOL model. IP_3_ concentration, at a location between the PM and SR, as a function of time for different PM-SR distances in rectangular geometries shown in Figure 3B. (B) AUC for IP_3_ shows an increase between 50 and 80 nm with a peak at 70 nm and decreases for increasing PM-SR distances from there on, suggesting that the small PM-SR distances are best suited for IP_3_ signaling.

**Supplementary Figure 8.**
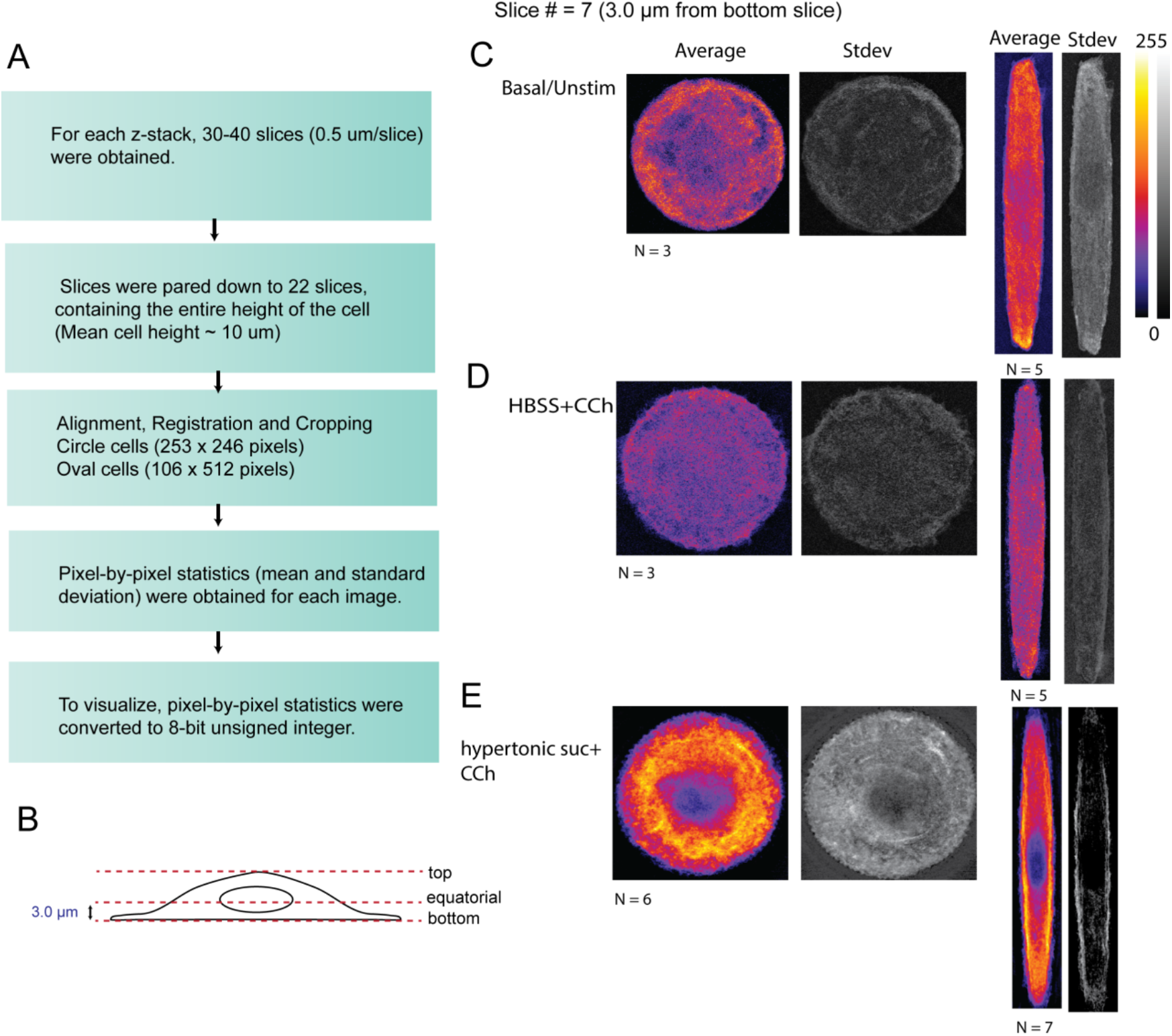
Quantitative Immunofluorescence workflow to determine distribution of M_3_R under different conditions. A) Processing of Z-stacks obtained from individual cells B) Images were obtained at z = 3.0 μm from the bottom of the cell. Representative images obtained at C) basal/unstimulated D) stimulated with 10 μM Carbachol in HBSS and E) Stimulated with 10 μM Carbachol in the presence of hypertonic sucrose at 4 °C to inhibit receptor endocytosis. Scalebars shown are 10 μm.

**Supplementary Figure 9.**
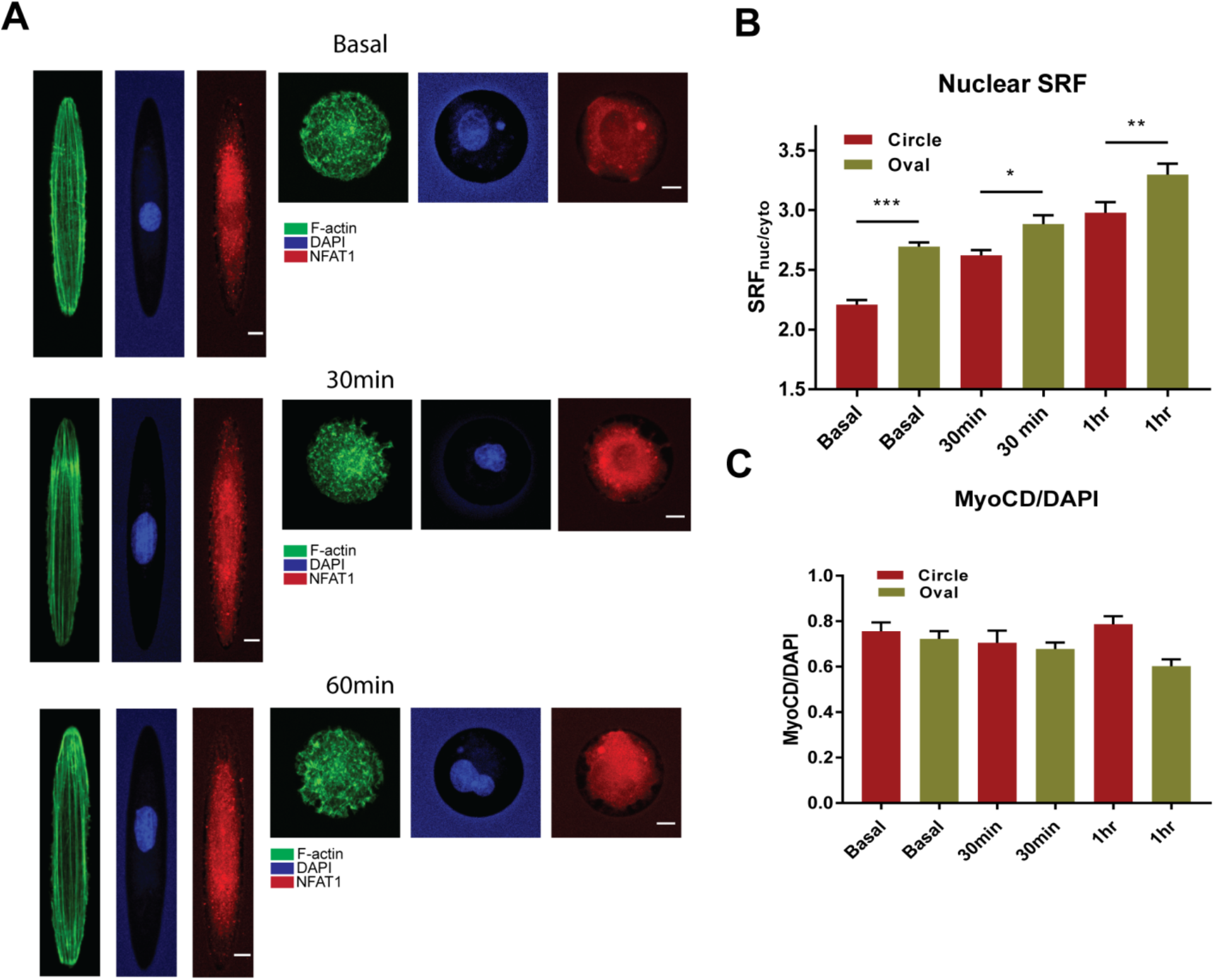
NFAT and SRF are impacted by changes in shape while myocardin is not. A) Representative images of NFAT after stimulation with carbachol showing nuclear translocation of NFAT. Scale bars shown in NFAT (red) are 10 μm. B) SRF localization into nucleus is impacted by cell shape (shown are mean ± SEM, N_circle_,^basal^ = 118, N_oval_,^basal^ = 120, N_circle_,^30min^ = 152, N_oval_,^30min^ = 133, N_circle_,^1hr^ = 99, N_oval_,^1hr^ = 132), C) MyoCD was constitutively in the nucleus. No change in myocd nuclear increase was seen with carbachol stimulation (shown are mean ± SEM, N_circle_,^basal^ = 17, N_oval_,^basal^ = 5, N_circle_,^30min^ = 9, N_oval_,^15min^ = 17, N_circle_,^1hr^ = 26, N_oval_,^1hr^ = 13). (*, **, *** denote p < 0.05, p < 0.001 and p < 0.0001 respectively, denoting statistical significance between the two cell shape, circle in red and oval in green, using two-tailed Student’s t-test).

**Supplementary Figure 10.**
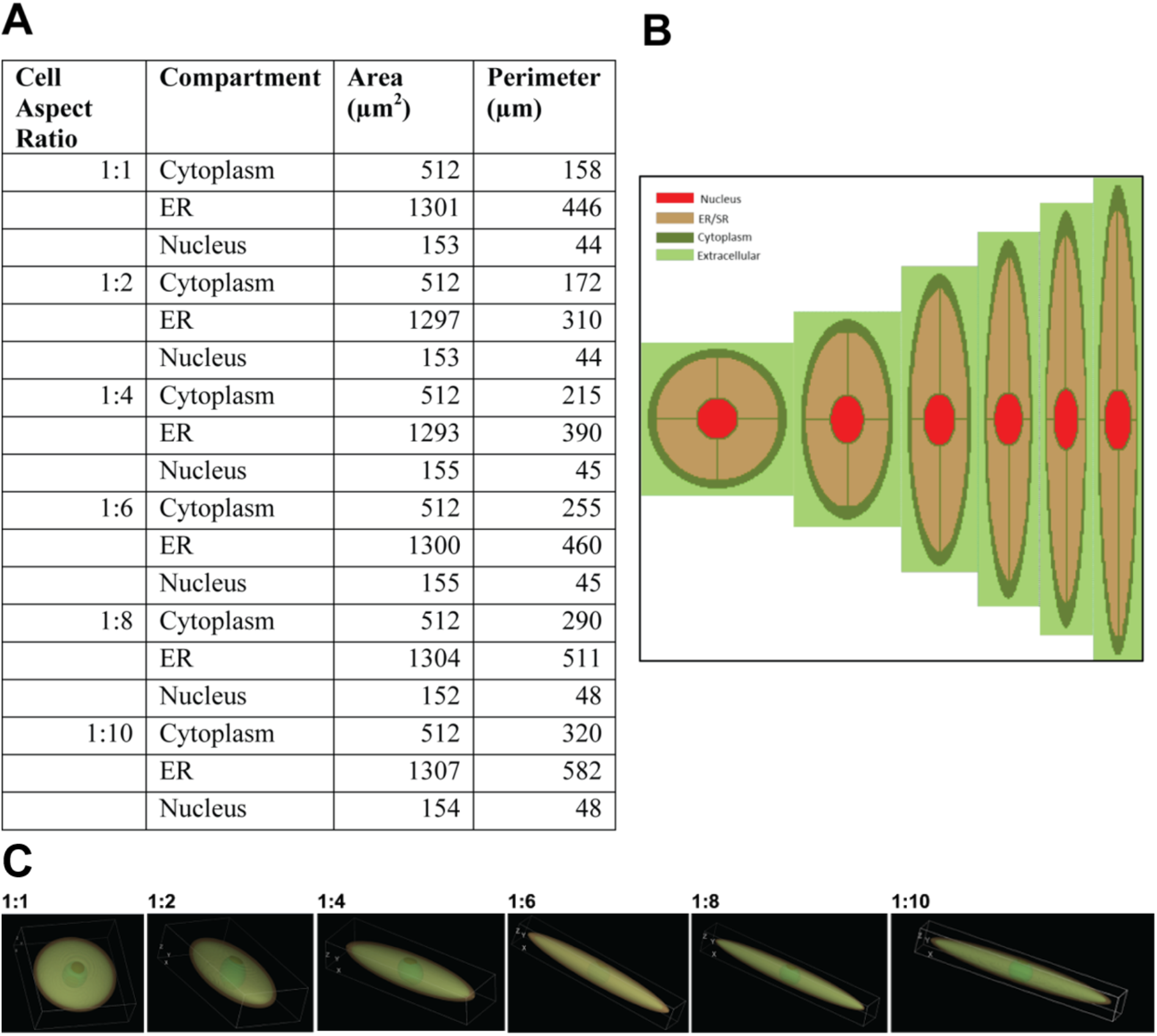
A) Geometric region details used for the spatial model using *Virtual Cell* and B) resulting 2D geometries with increasing aspect ratio. C) 3D geometries of cells with constant volume and increasing surface area to volume ratio representing plasma membrane, SR membrane and nucleus in increasing elliptical aspect ratios.

**Supplementary Figure 11.**
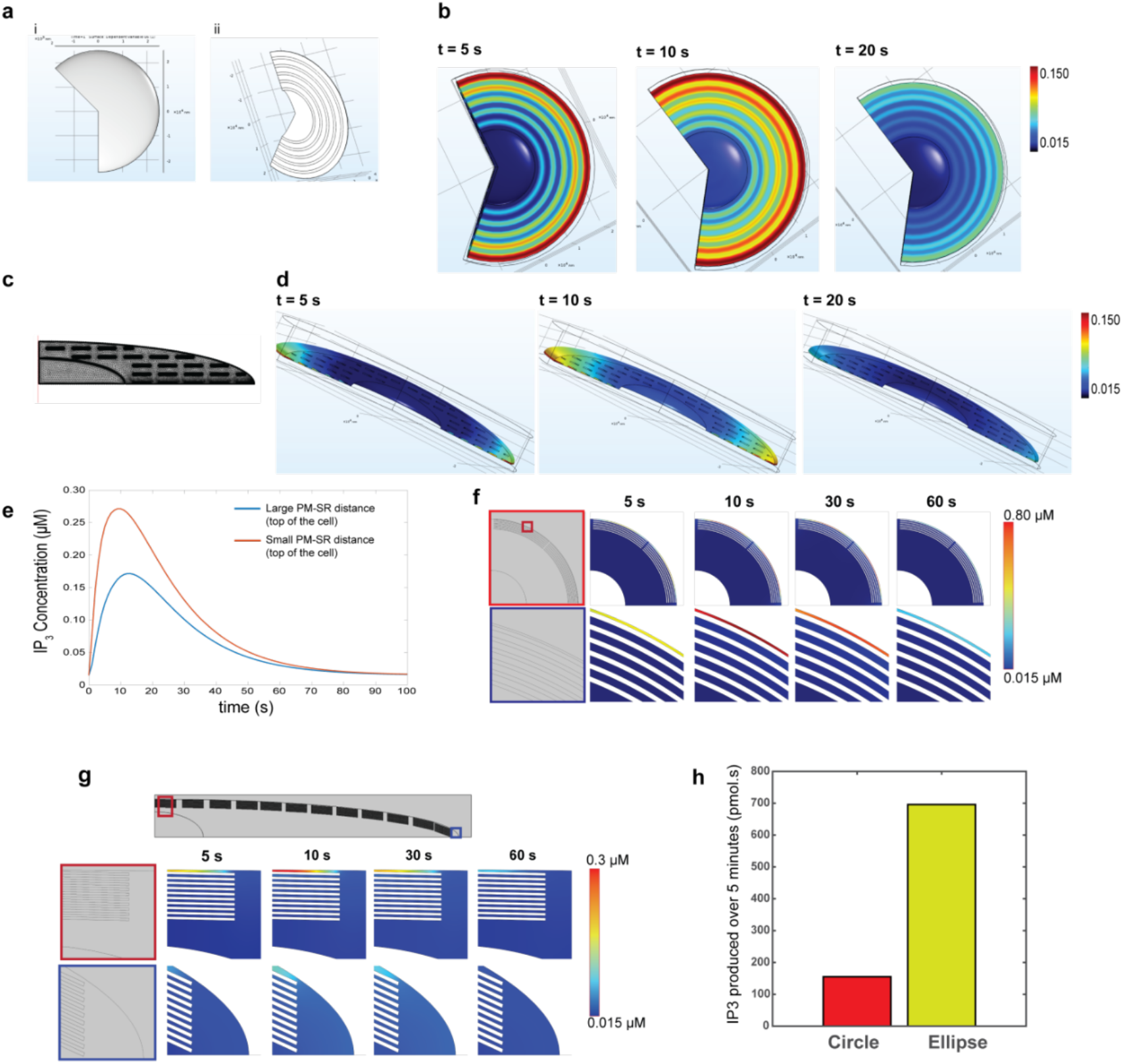
3D spatial models provide mechanistic understanding of cell elongation-induced Ca^2+^ potentiation. 3D simulations show the effect of cell elongation on inter-organelle distances on IP_3_/Ca^2+^ potentiation using COMSOL a) 3D geometry to represent a cell with a circular 2D cross section and the ER as tori placed at different distances from the PM with the finite element mesh. b) IP_3_ spatial dynamics at 5, 10, and 20s in the geometry shown in panel a. c) 3D geometry to represent a cell with an elliptical cross section with rings of ER. d) IP_3_ spatial dynamics at 5, 10, and 20 s in the ellipsoidal geometry. e) IP_3_ dynamics at two different points near and far from the PM showing the PM-SR distance affects IP_3_ transience. f -h) IP_3_ concentrations in 2D circular geometries (f) and 2D elliptical geometries (g). h) IP_3_ concentrations over space (the entire domain) and time were integrated, since integration of signal over time provides a buffer for time scale and damps out the effect of temporal fluctuations and spatial integration allows for conversion from concentration (moles per unit volume) to amount (moles). Shown are graphs comparing integrated IP_3_ in circular and elliptical geometries.

**Supplementary Figure 12.**
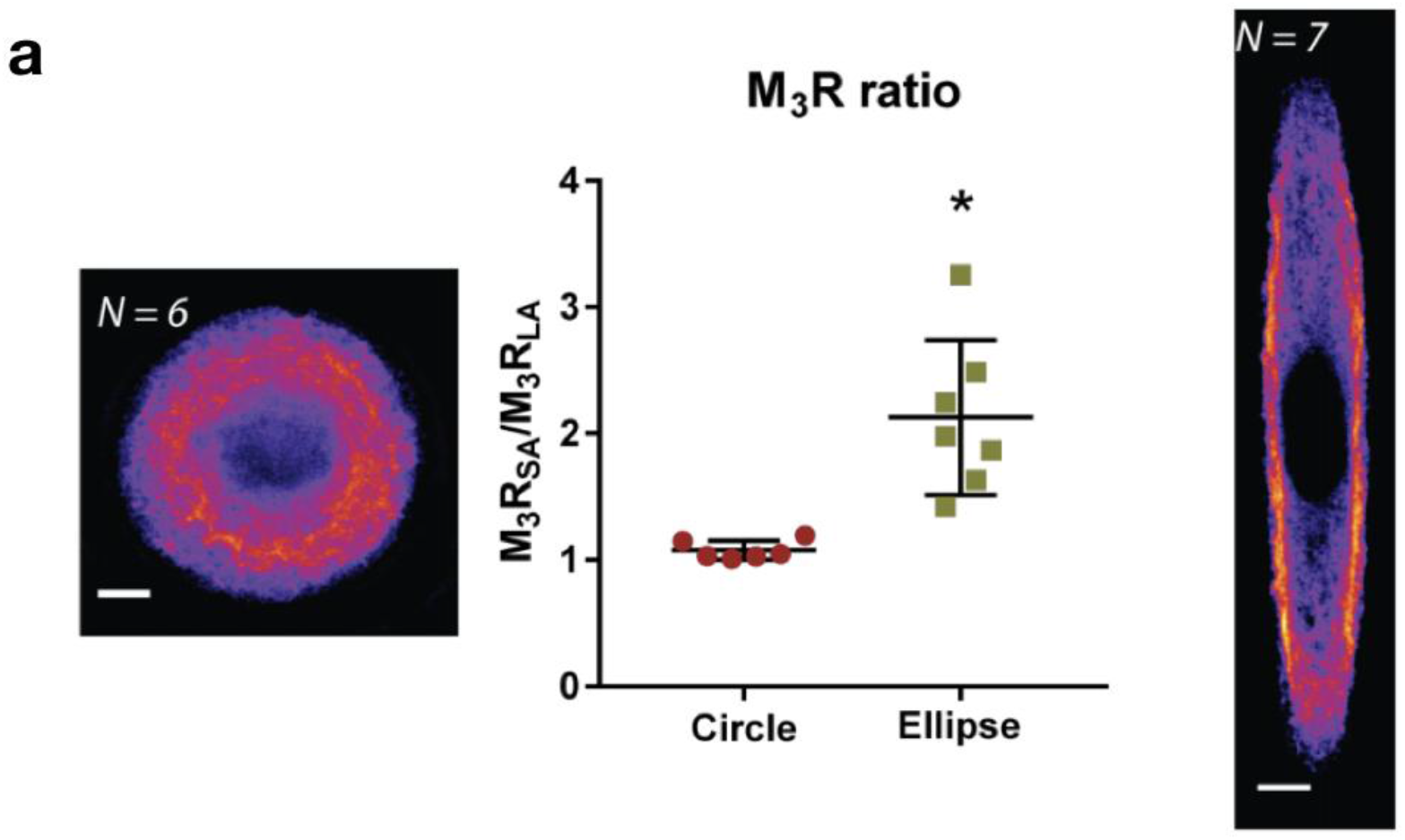
Shape impacts M_3_R/Ca^2+^ signaling in VSMC. (a) Average intensity of M3R along the length of the cell in circular (*N=6)* and oval *(N=7)* VSMC. Differences in anisotropic distribution of M_3_R were quantified by obtaining the ratio of M_3_R staining intensities on the short and long axes (M_3_R_SA_/M_3_R_LA_, mean±SEM ellipse=2.1 ± 0.6 A.U. *N = 7*, circle= 1.1 ± 0.1 A.U., *N=6*, *P=0.0016, two-tailed t-test). Scale bars shown are 10 μm.

**Supplementary Figure 13.**
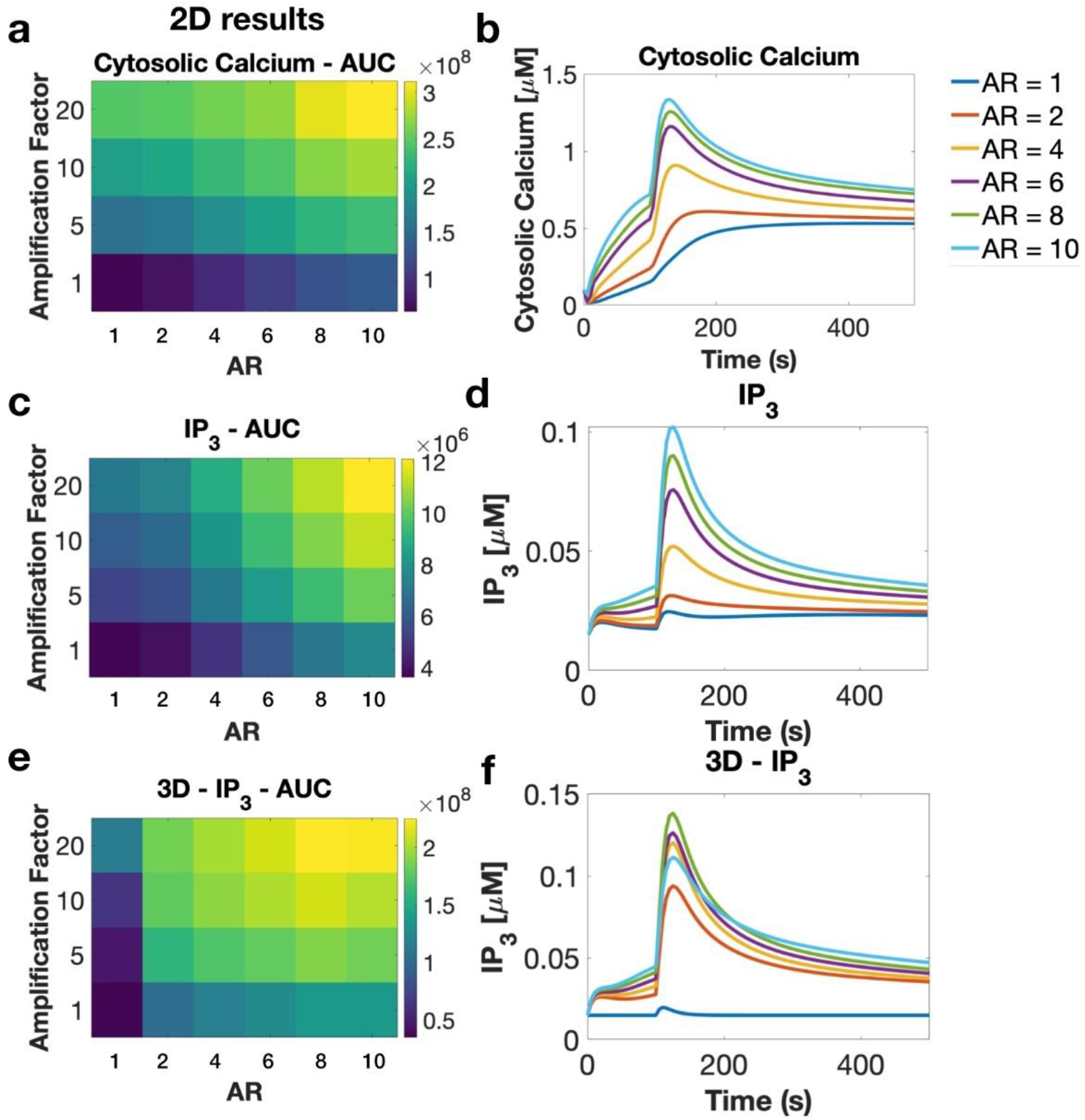
Simulation results for 2D and 3D geometries. a-d) 2D simulation results for cytosolic calcium and IP_3_ dynamics. AUC and temporal dynamics for different ARs and SR flux amplification factors. e-f) 3D simulation results for IP_3_ dynamics for geometries of different ARs and SR flux amplification factors. Inset: legend for all temporal simulation results.

**Supplementary Figure 14.**
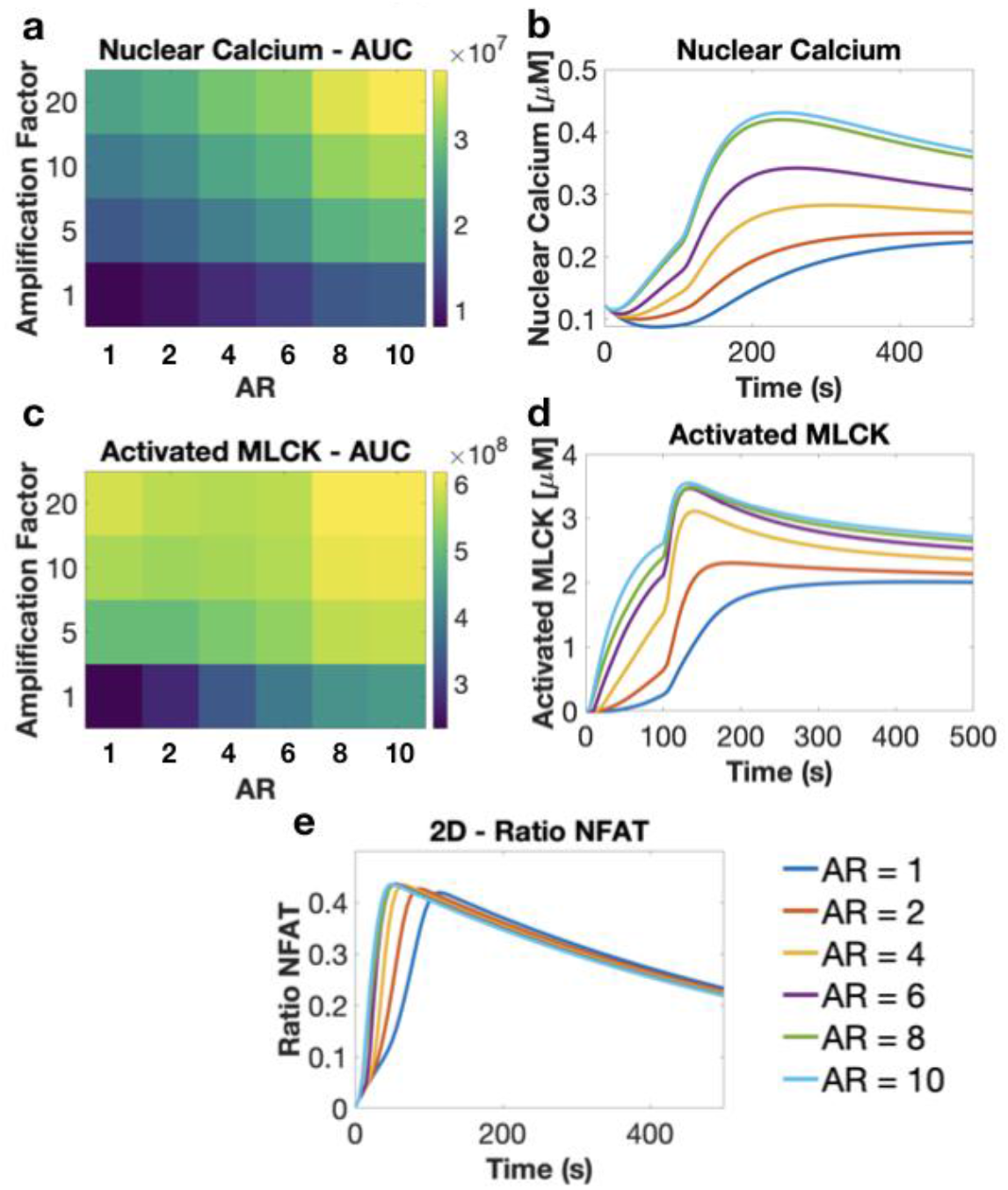
2D simulation results for downstream signaling. a-e) 2D simulation results for nuclear calcium, activated MLCK, and NFAT dynamics. AUC and temporal dynamics for different ARs and SR flux amplification factors. Inset: legend for all temporal simulation results.

**Supplementary Figure 15.**
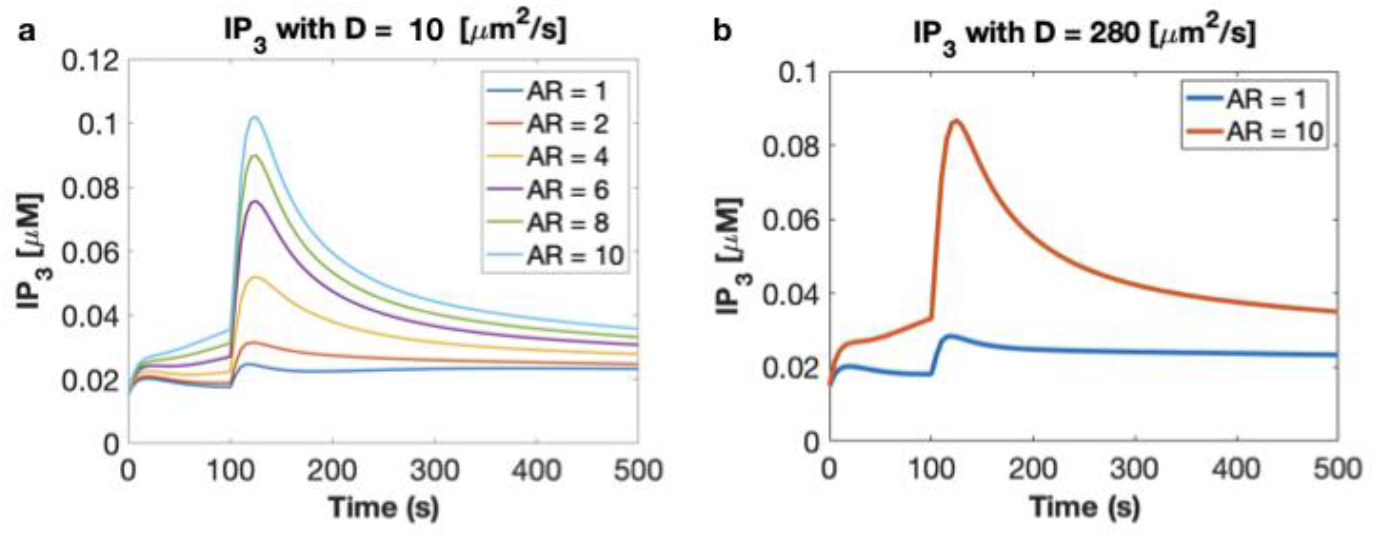
Effect of IP_3_ diffusion coefficient. a-b) IP_3_ dynamics for different IP_3_ diffusion coefficients 10 versus 280. We see the same trend regardless of diffusion coefficient.

**Supplementary Figure 16.**
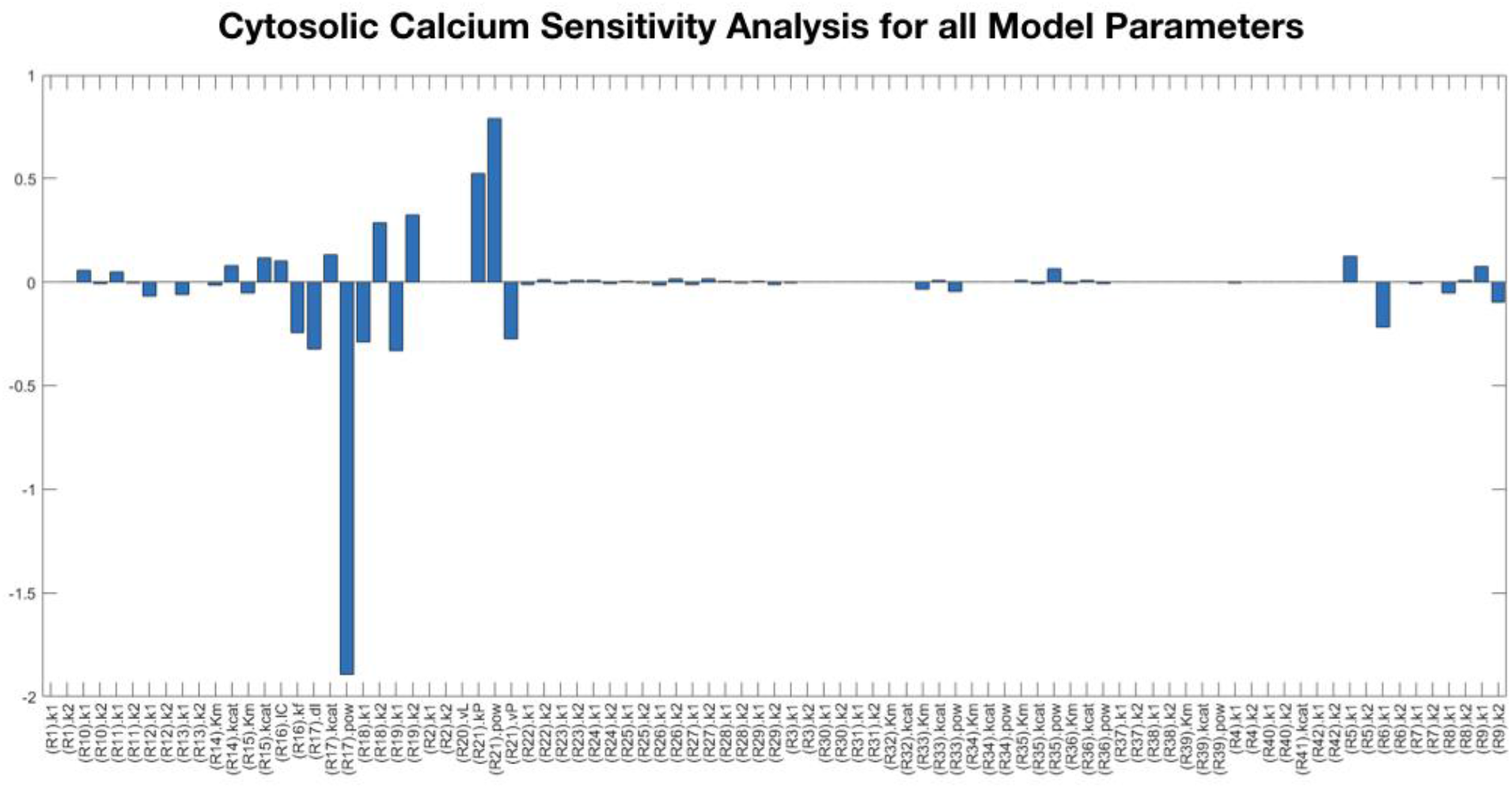
Sensitivity Analysis shows how different model parameters affect cytoplasmic calcium transients. The model is particularly sensitive to R17, which means that calcium release from the SR due to IP_3_R is particularly important to the model.

**Supplementary Figure 17.**
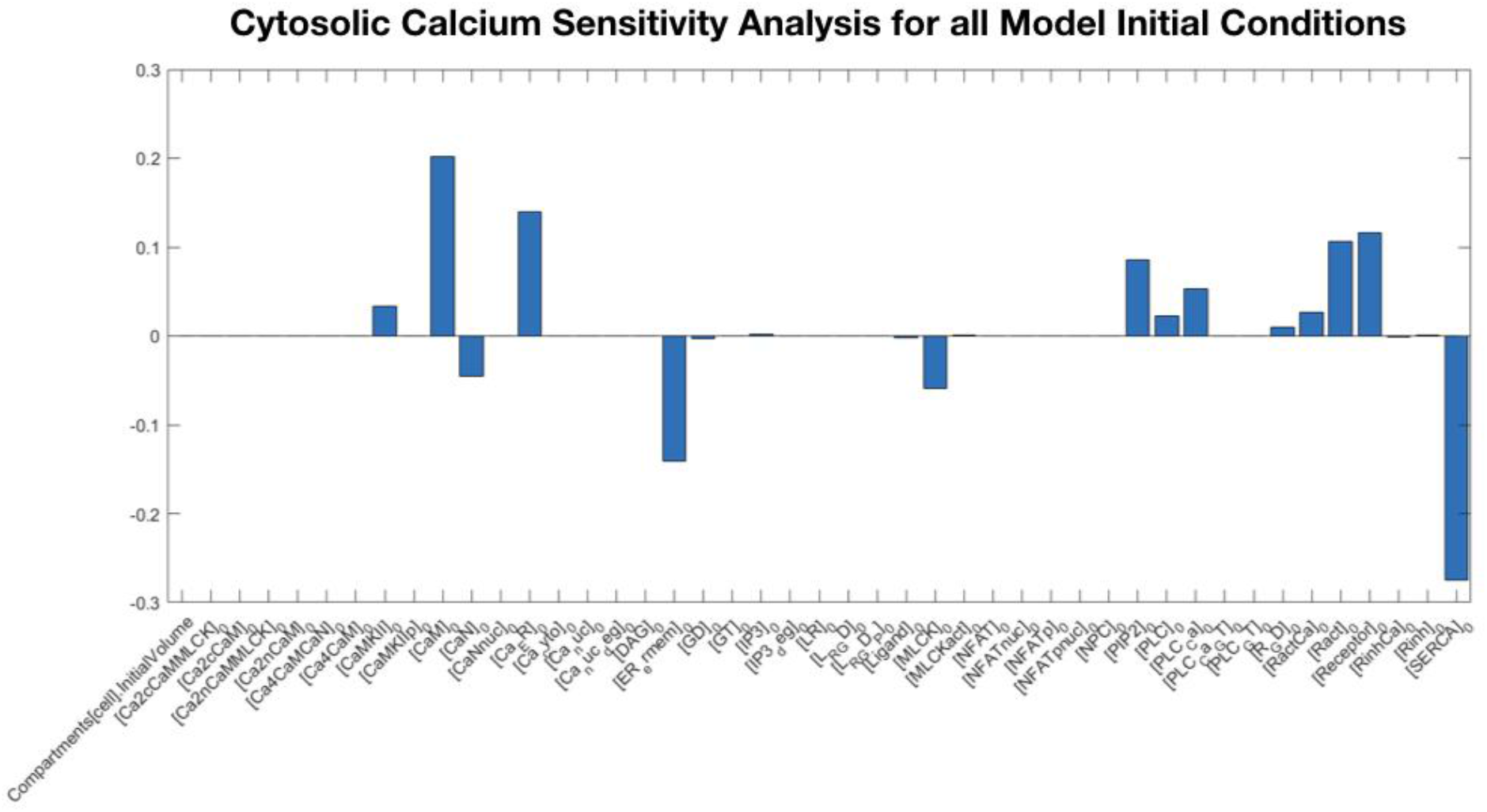
Sensitivity Analysis for cytoplasmic calcium transients with respect to model initial conditions. We see that several species have a larger impact than others. In particular, cytoplasmic calcium is sensitive to SERCA and ER_ermem_ density, and CaM and Ca_ER_ initial concentrations.

**Supplementary Figure 18.**
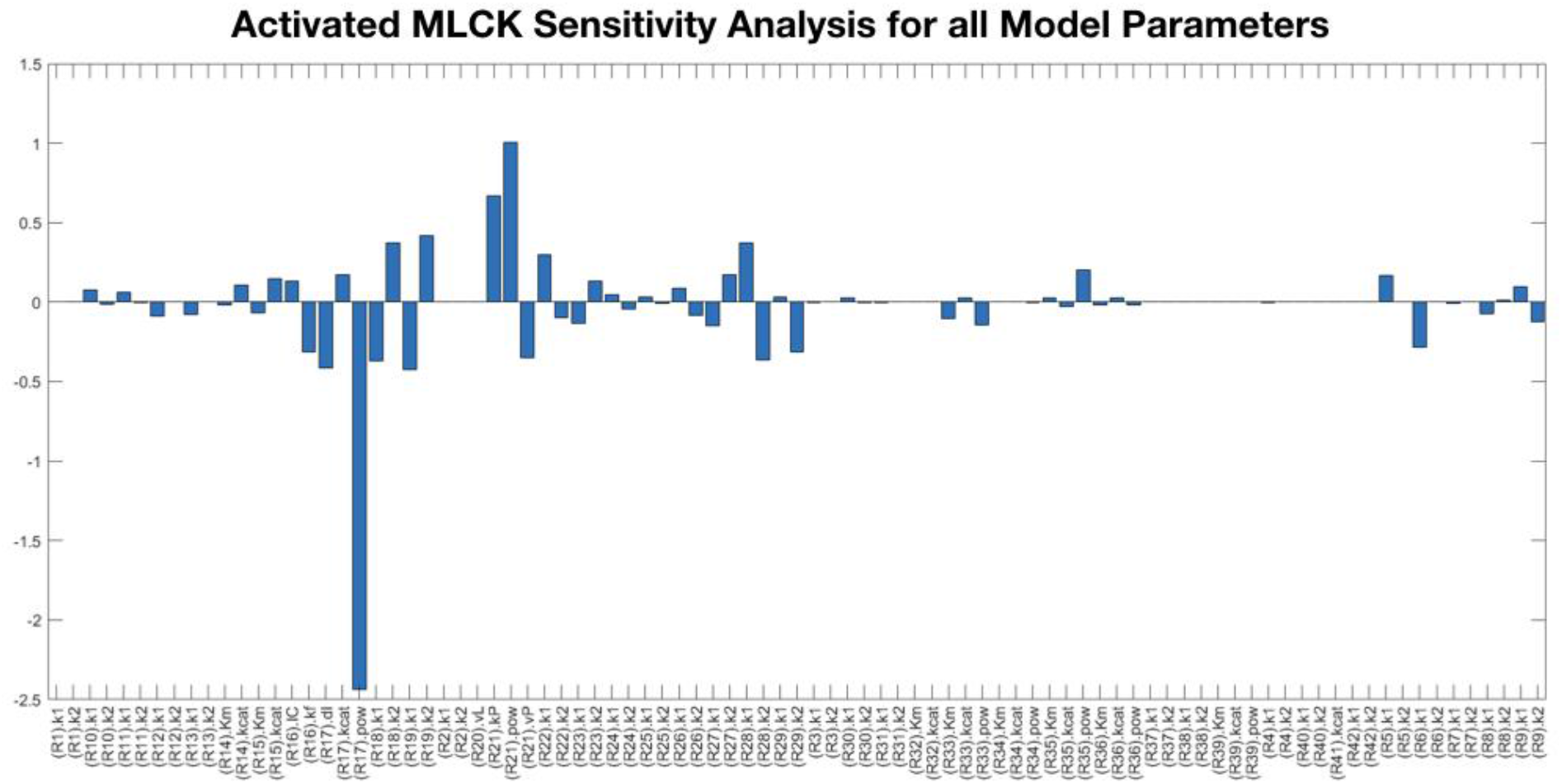
Sensitivity Analysis shows how all model parameters affect activated MLCK transients. We see that similarly to cytoplasmic calcium, activated MLCK depends on calcium dynamics associated with the SR, R17 and R21 in particular.

**Supplementary Figure 19.**
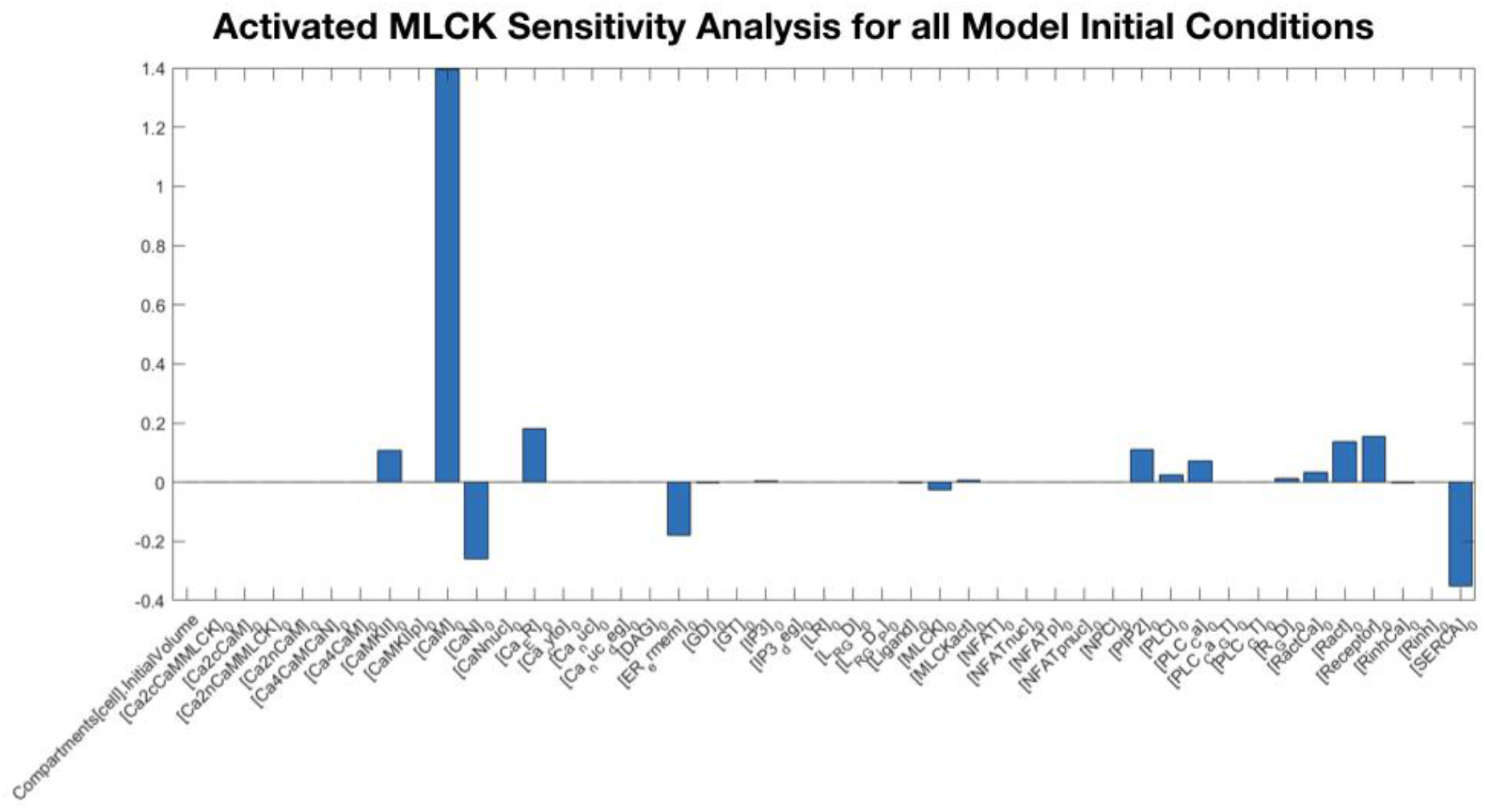
Sensitivity Analysis for activated MLCK transients with respect to model initial conditions. Activated MLCK is particularly sensitive to CaM initial concentration which makes sense because MLCK activation directly depends on CaM.

**Supplementary Figure 20.**
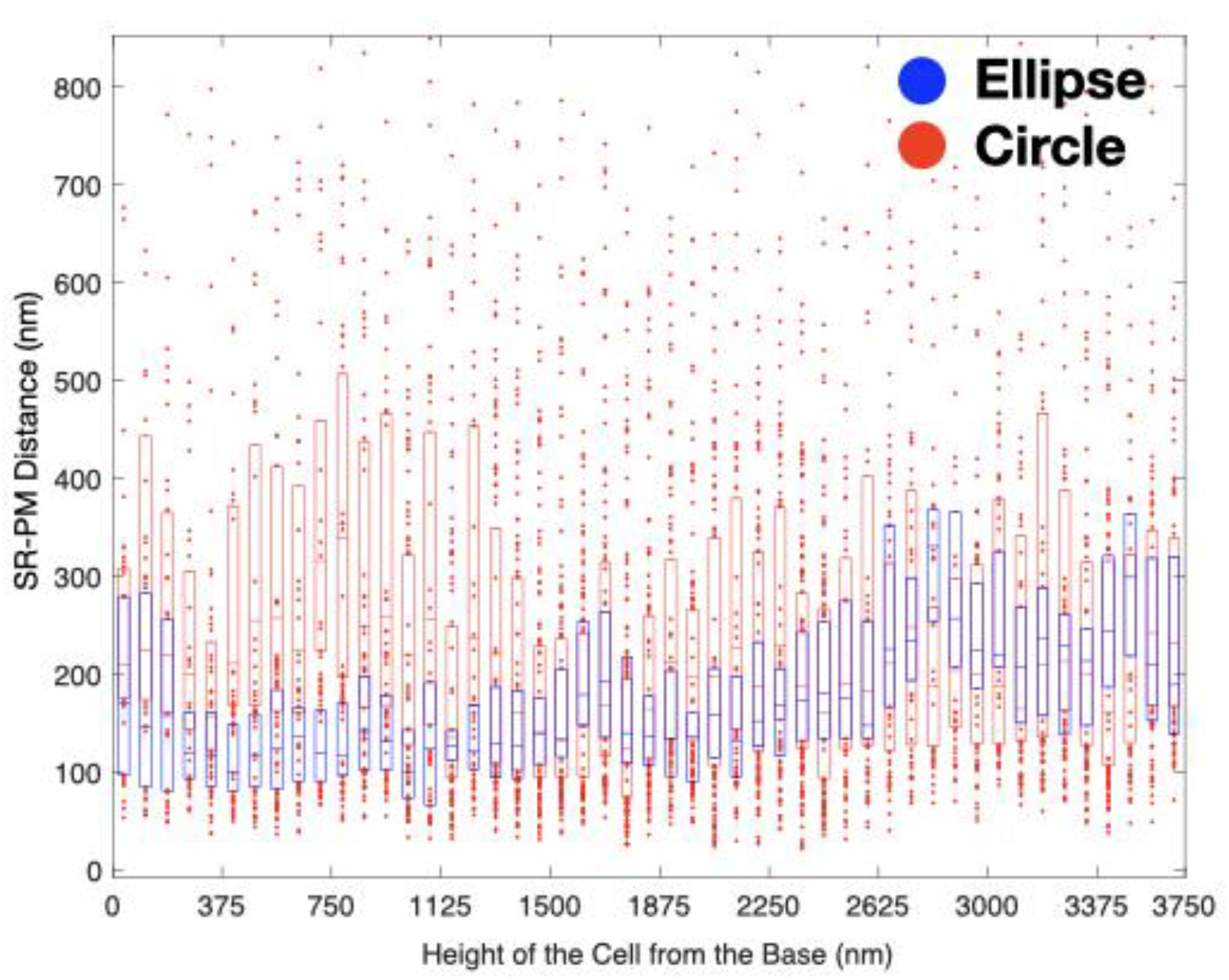
3D serial block face scanning electron microscopy was used to quantify the PM-SR distance in circular versus elliptical cells. The PM-SR distance per dyad was computed at each axial position and plotted as a function of representative circular and elliptical cells. 5309 and 1692 PM-SR distances were taken for the circular and elliptical cells, respectively.

### RAW CALCIUM DATA AND ANALYSES

Experimental calcium traces, their raw data, and the MATLAB script used to calculate their AUCs are listed below. Results from the data are found in Figures 5 and 6.

Each plot represents a cell conforming to a micropattern.

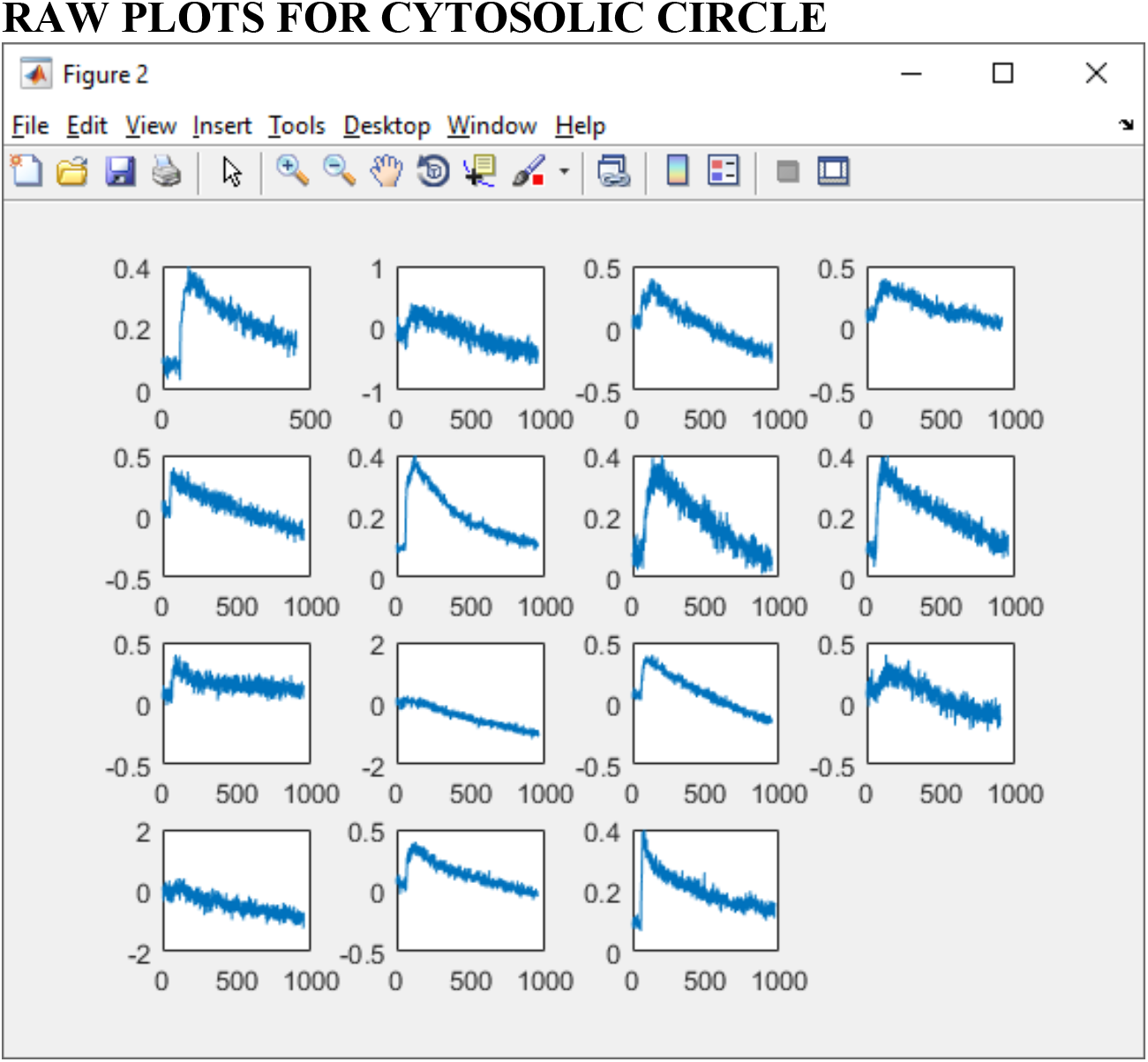

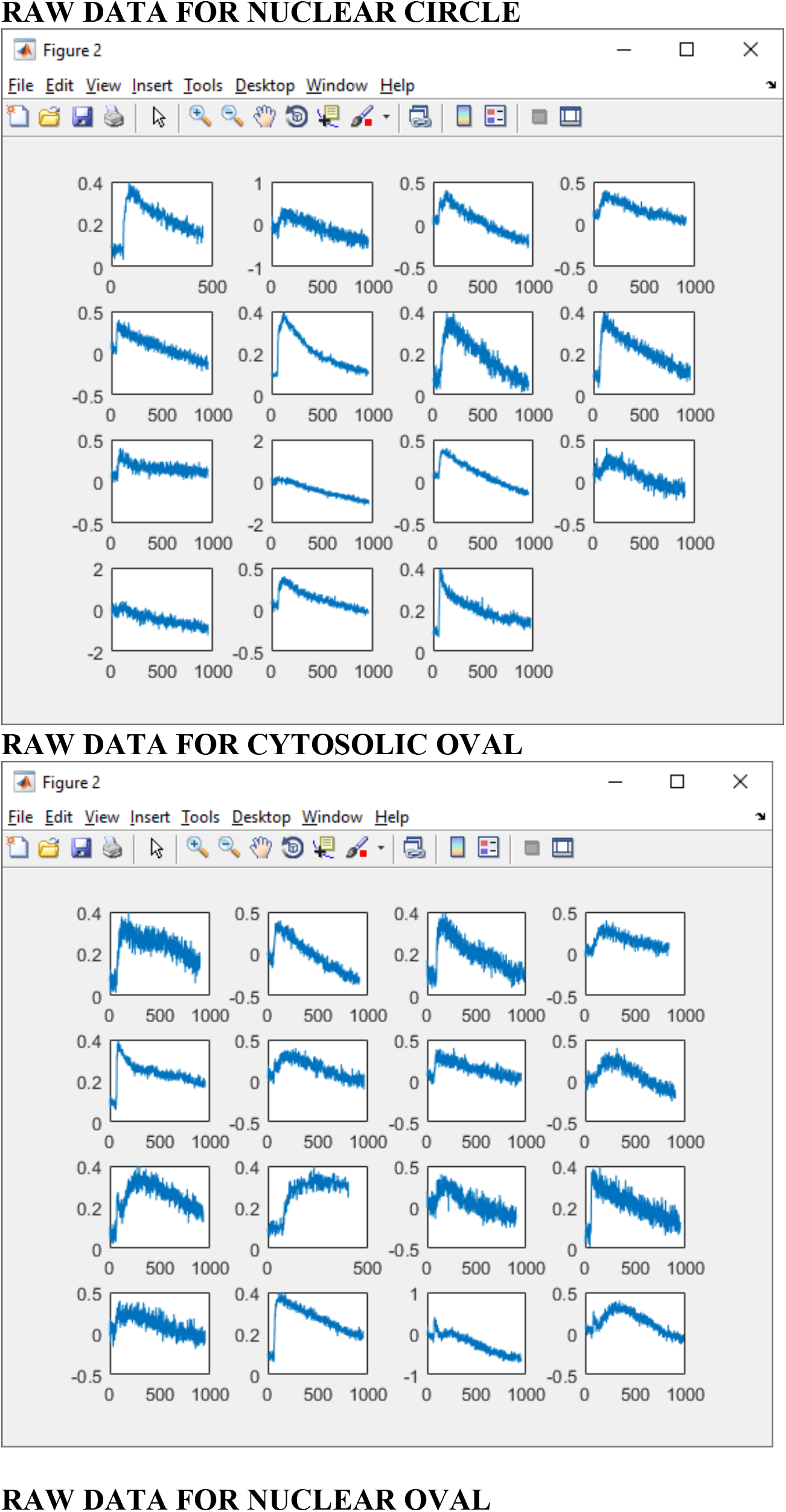

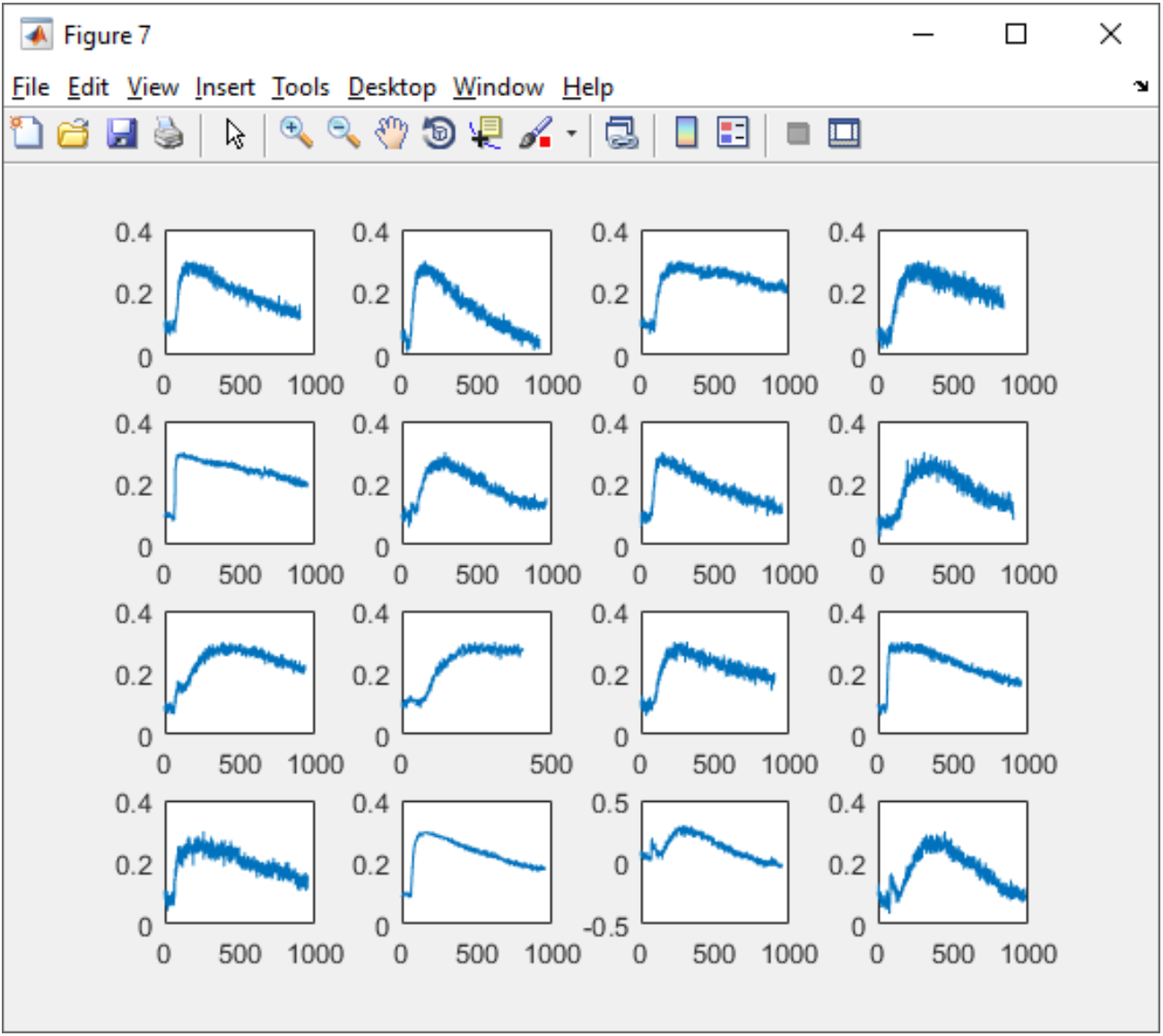

### RAW DATA OF EXPERIMENTAL AUC OBAINED USING THE CUSTOM-MADE AUC FUNCTION

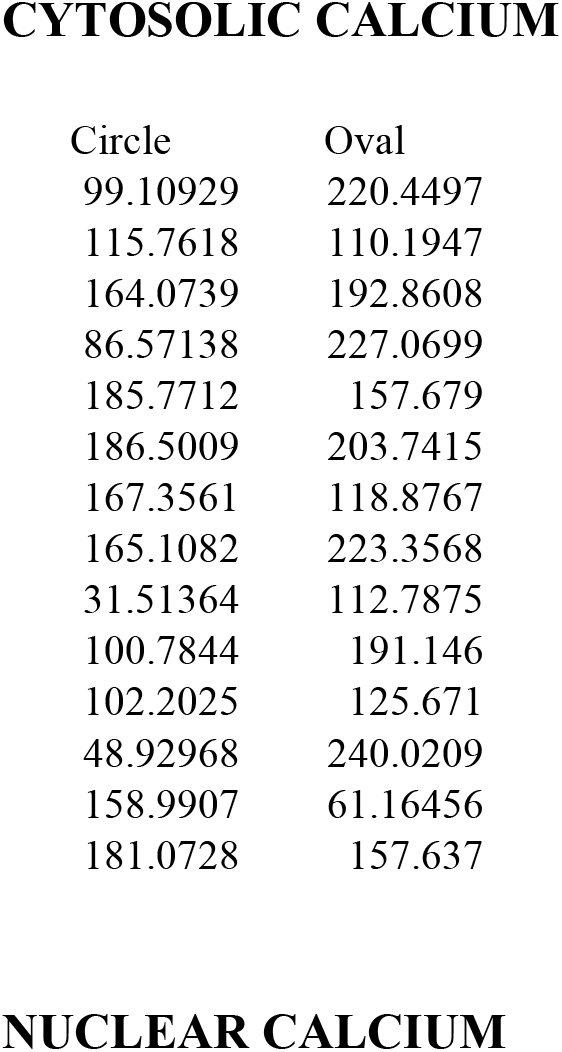

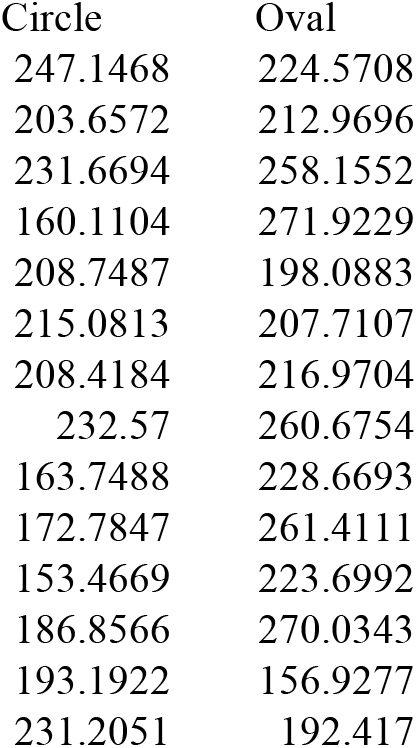

### SCRIPTS FOR ANALYZING IMAGING DATA FOR ELLIPSE AND CIRCULAR CELLS

#### Brief Description of the scripts

**Raw movie data were first imported using imageJ. An ROI with a specified size was used to specify nuclear and cytosolic regions. Raw intensity data was then normalized based on BAPTA and 100 uM A23187. AUC was calculated using the trapz function, as described by the following:**

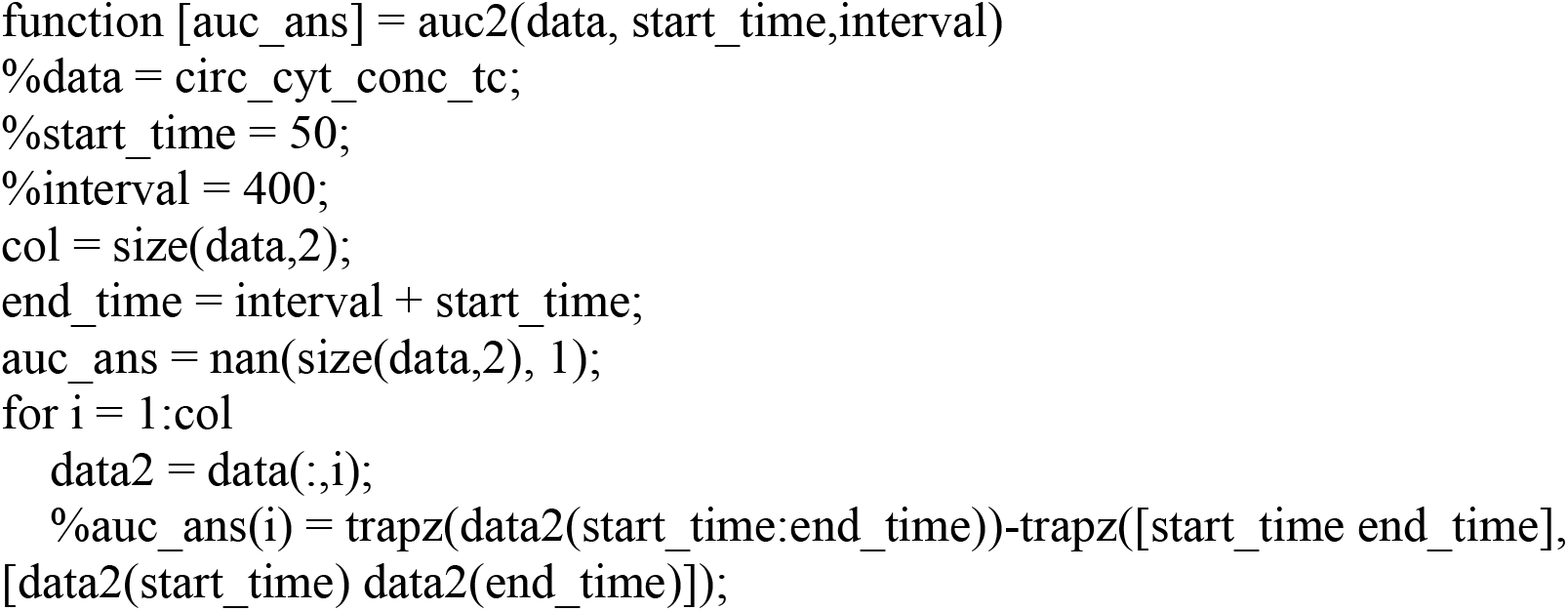

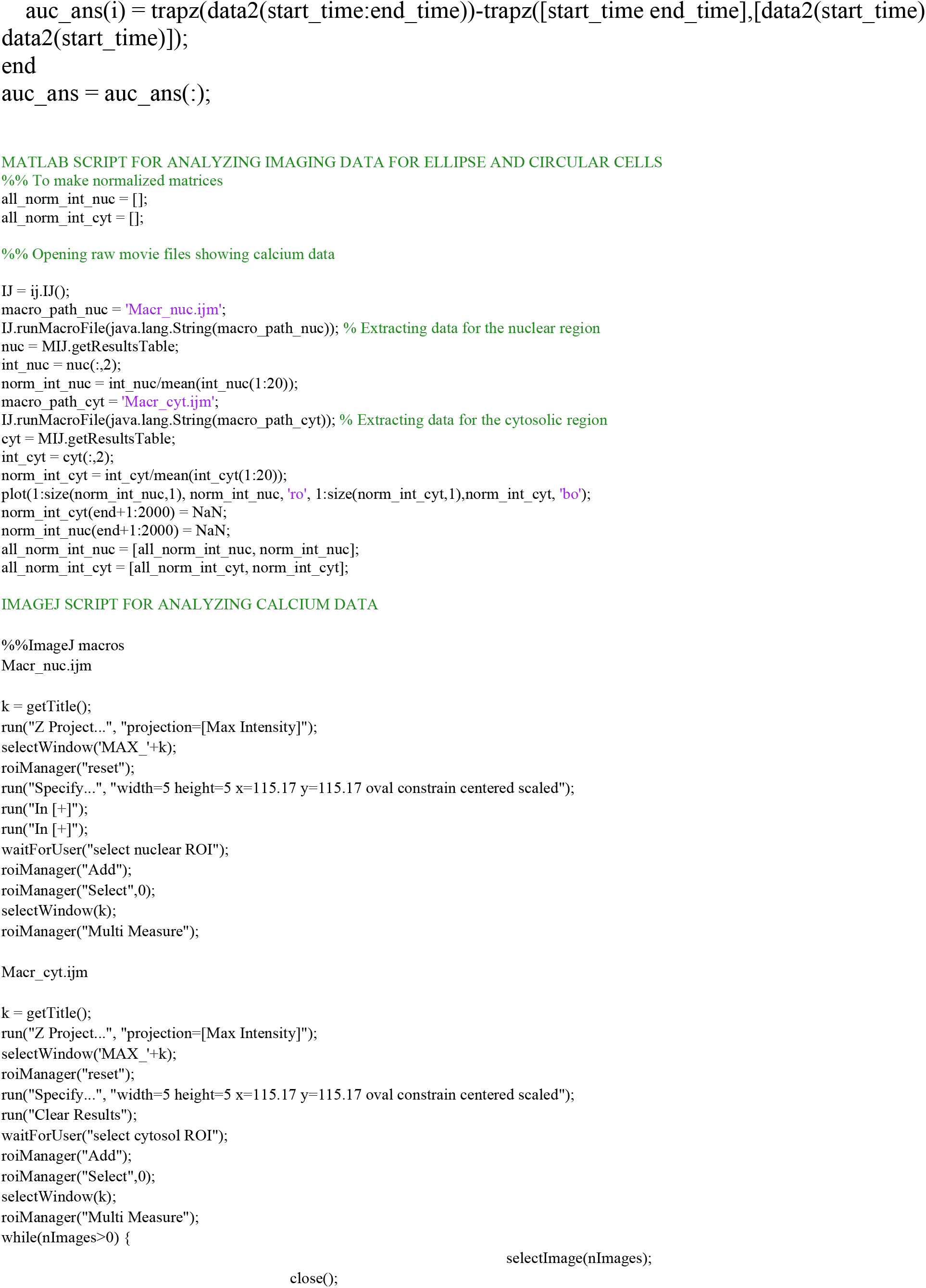

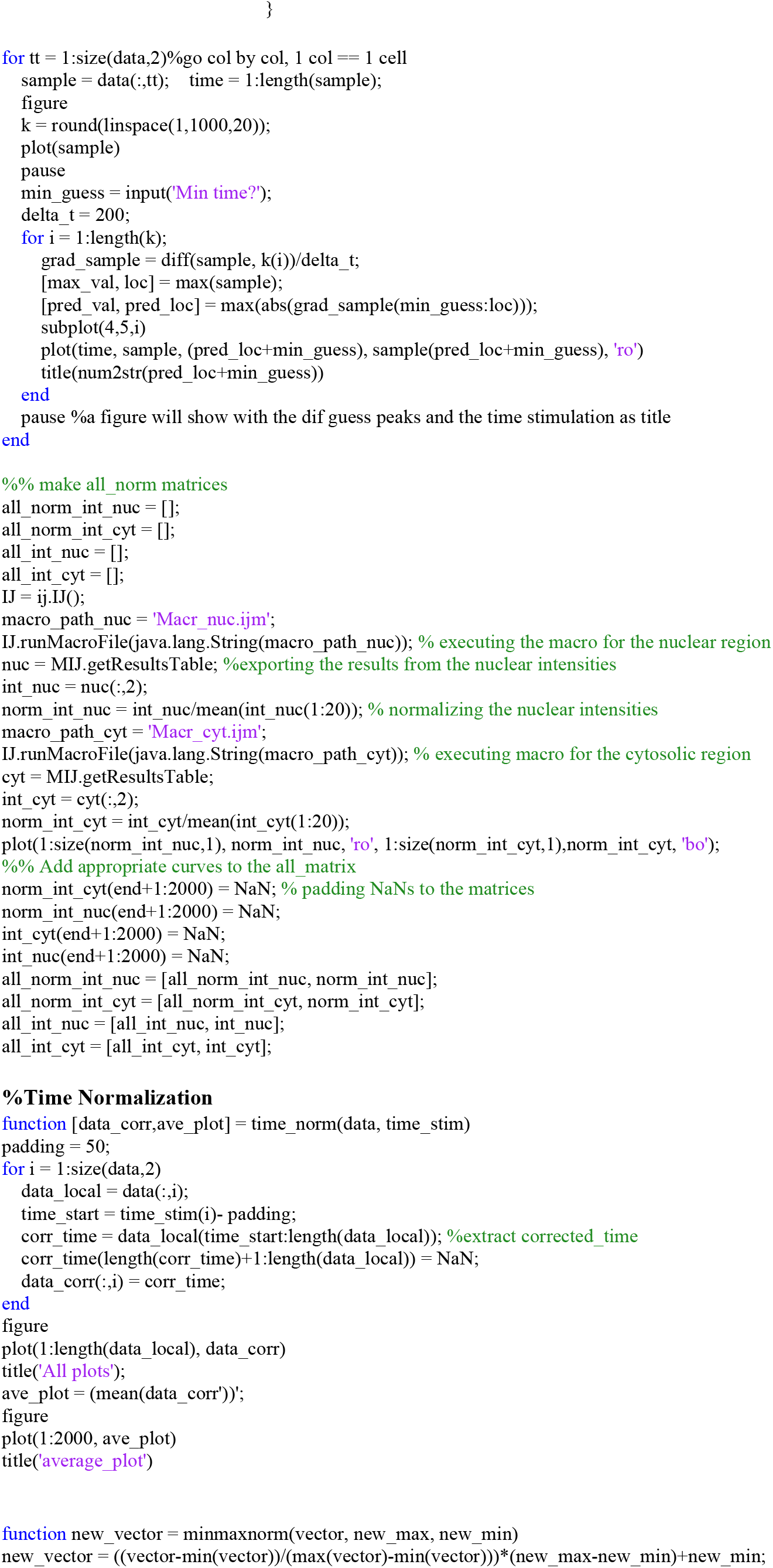

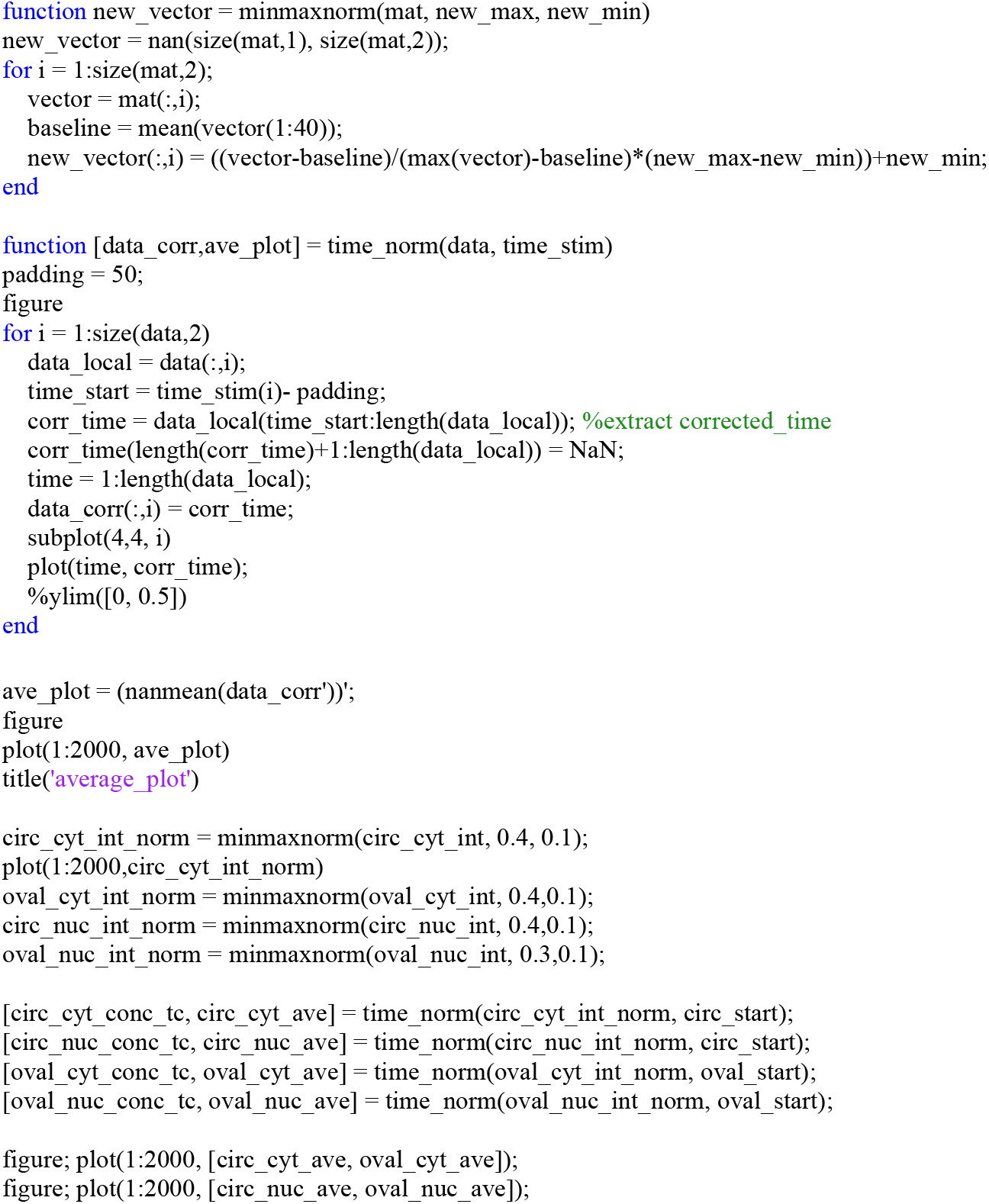

#### CONVERSION OF RATIOMETRIC TO CONCENTRATION

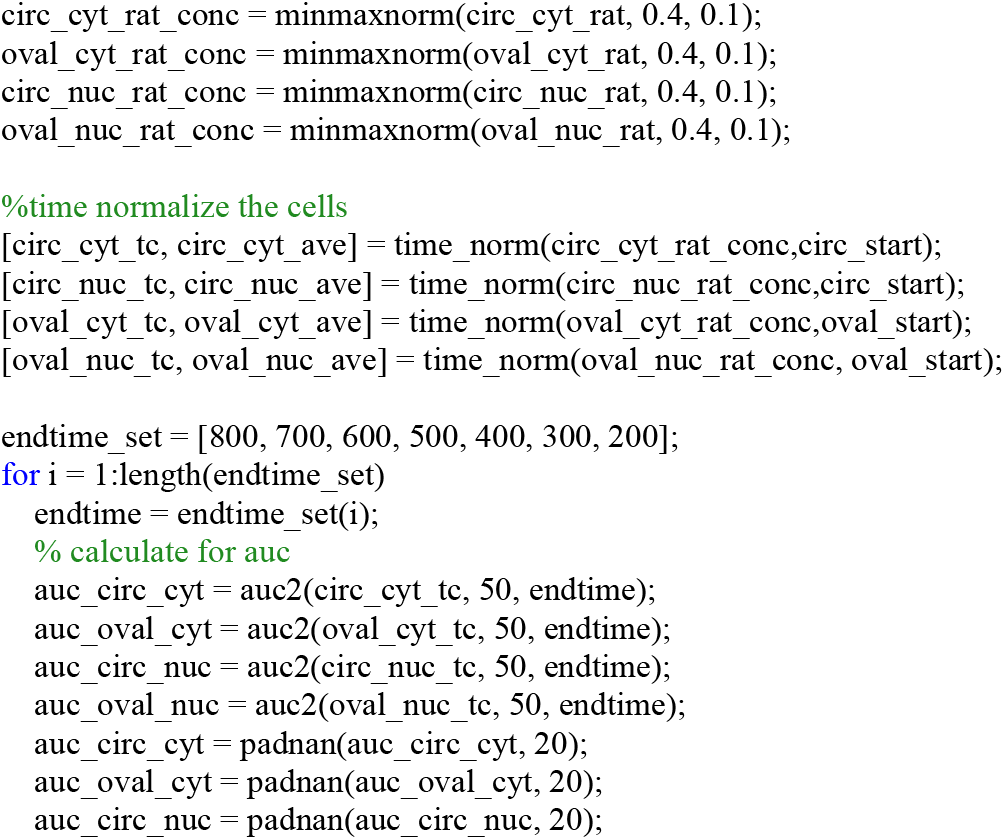

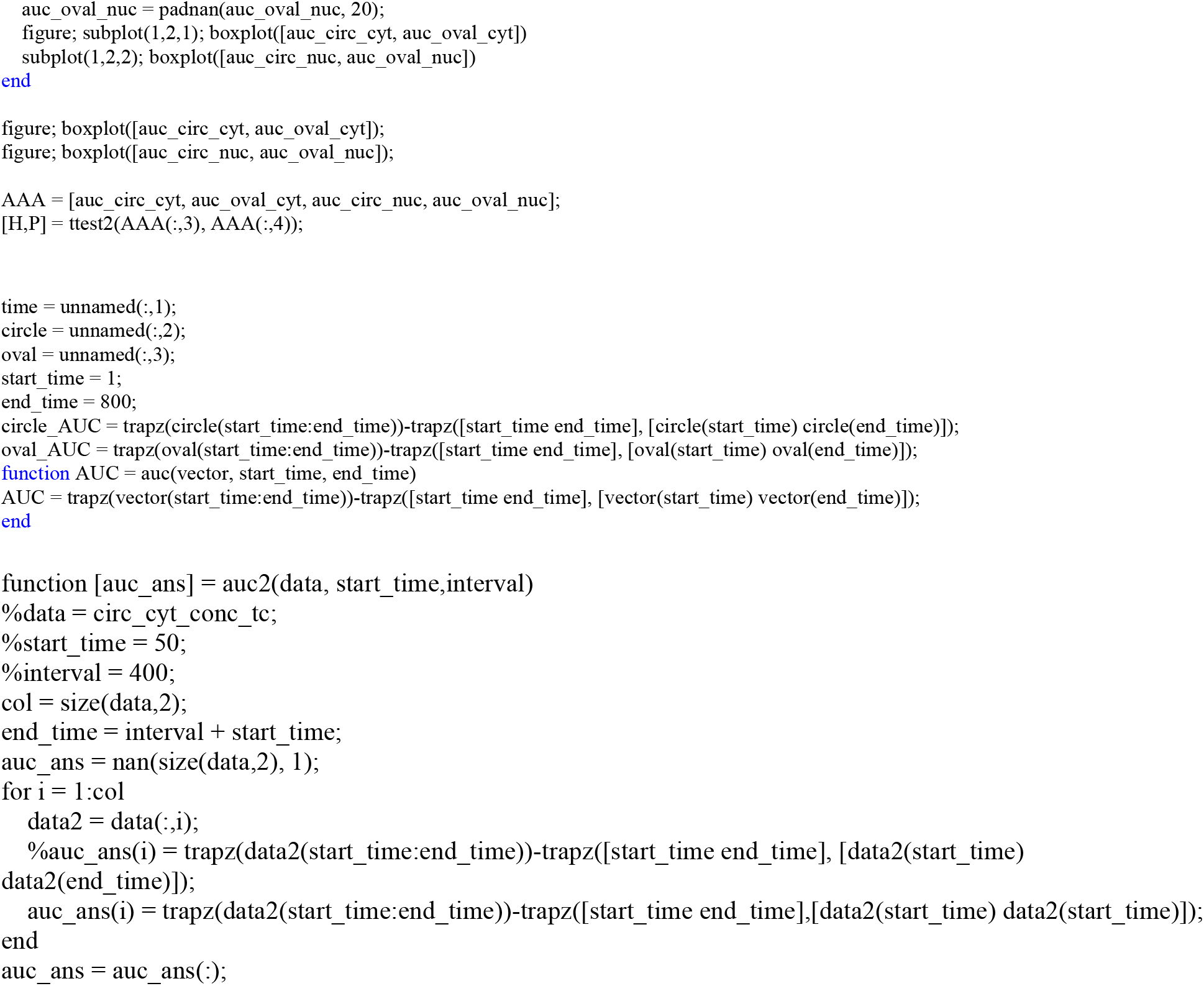

### FOR CALCULATING SIMULATION AREA UNDER THE CURVE

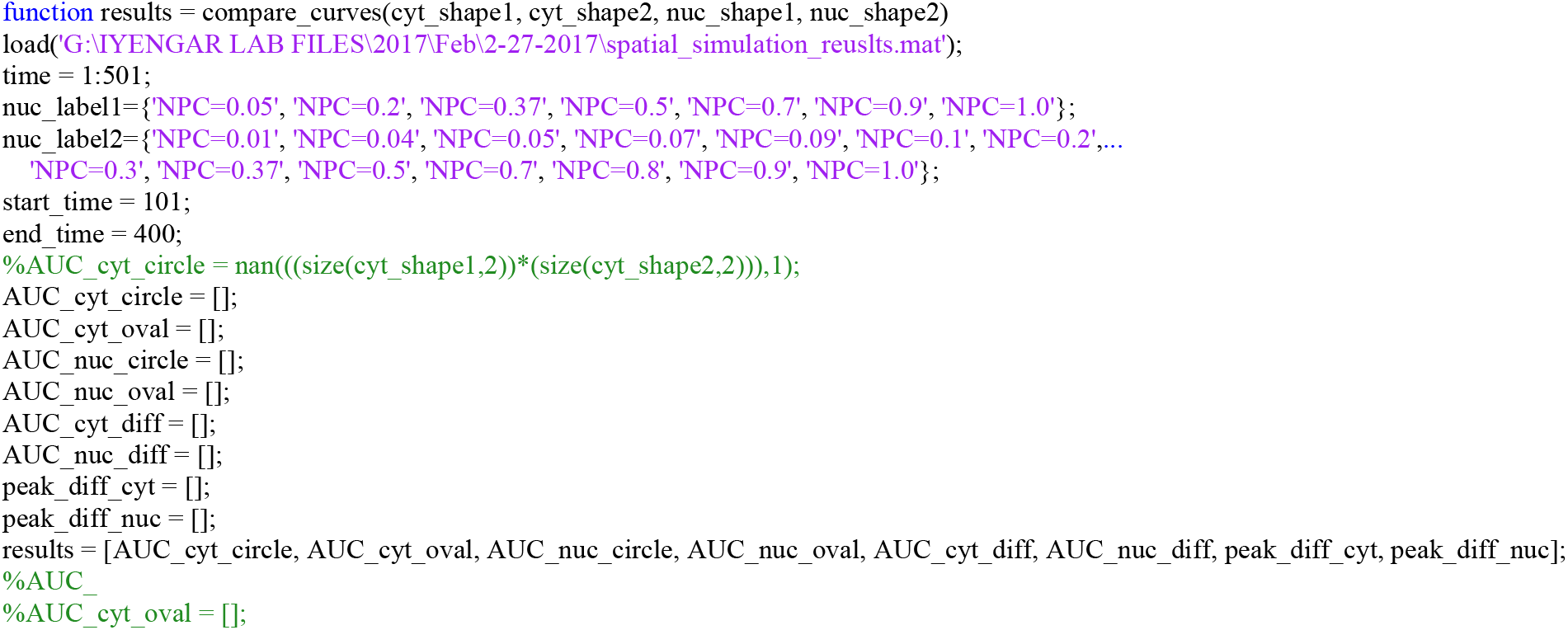

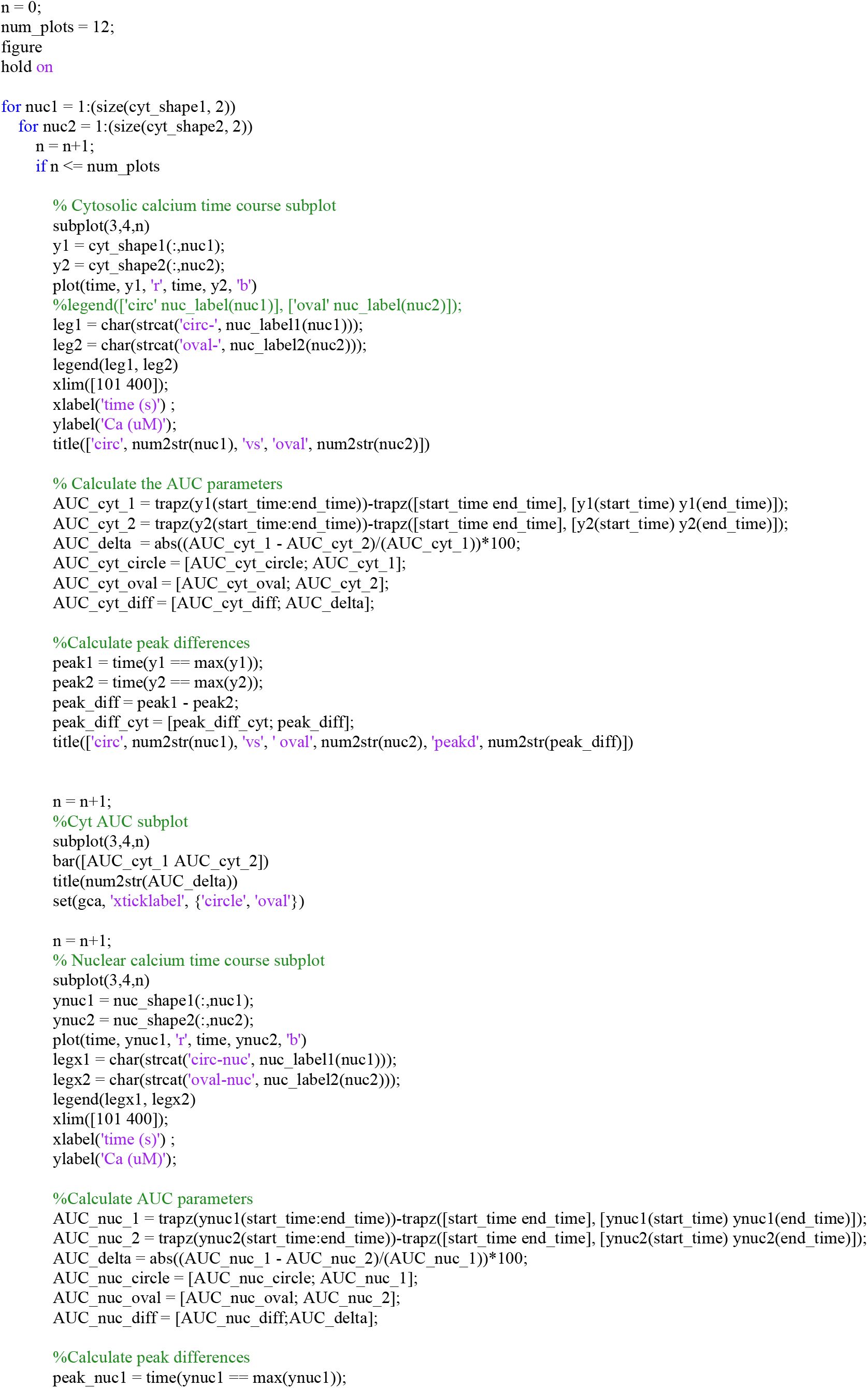

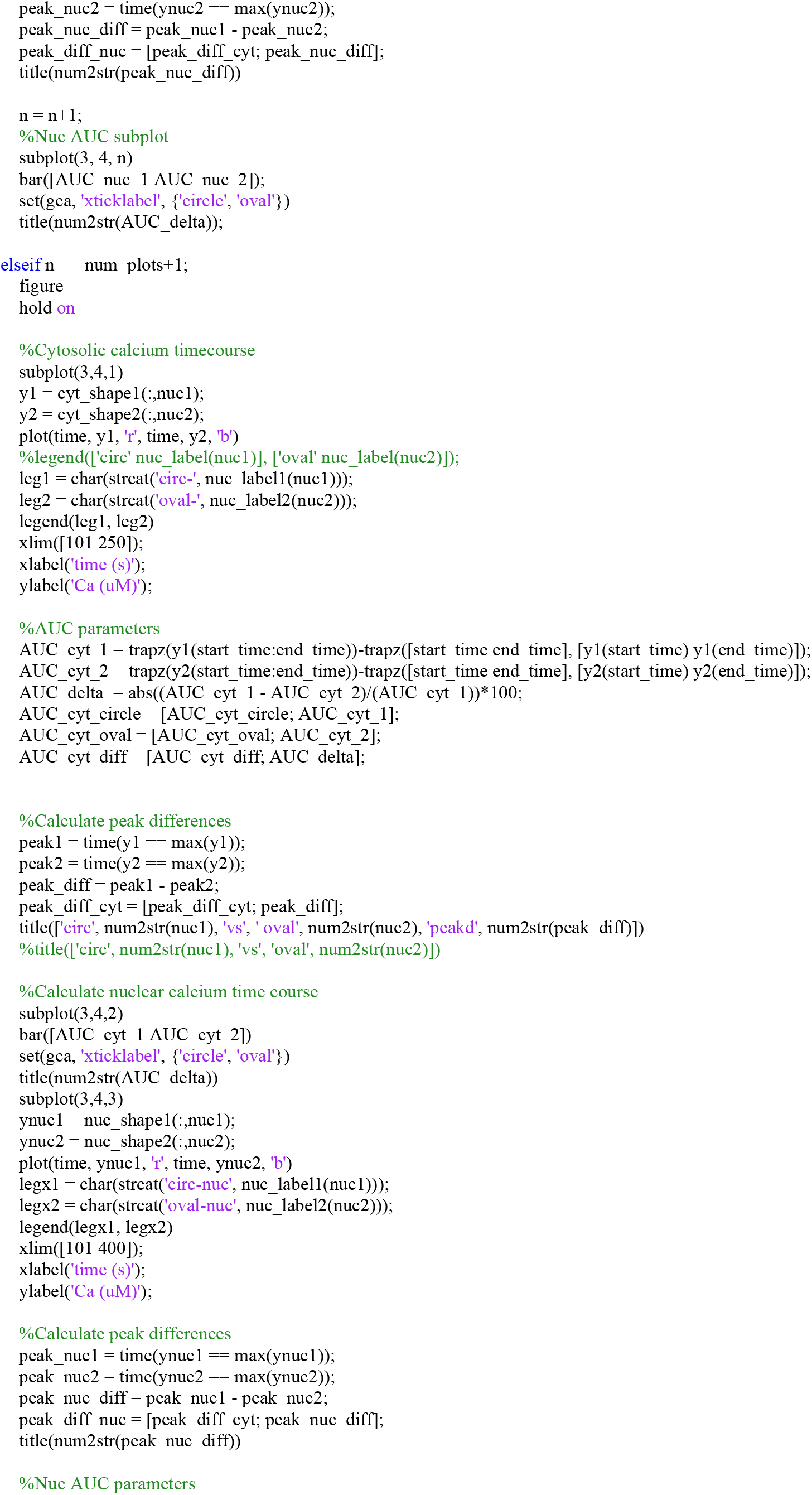

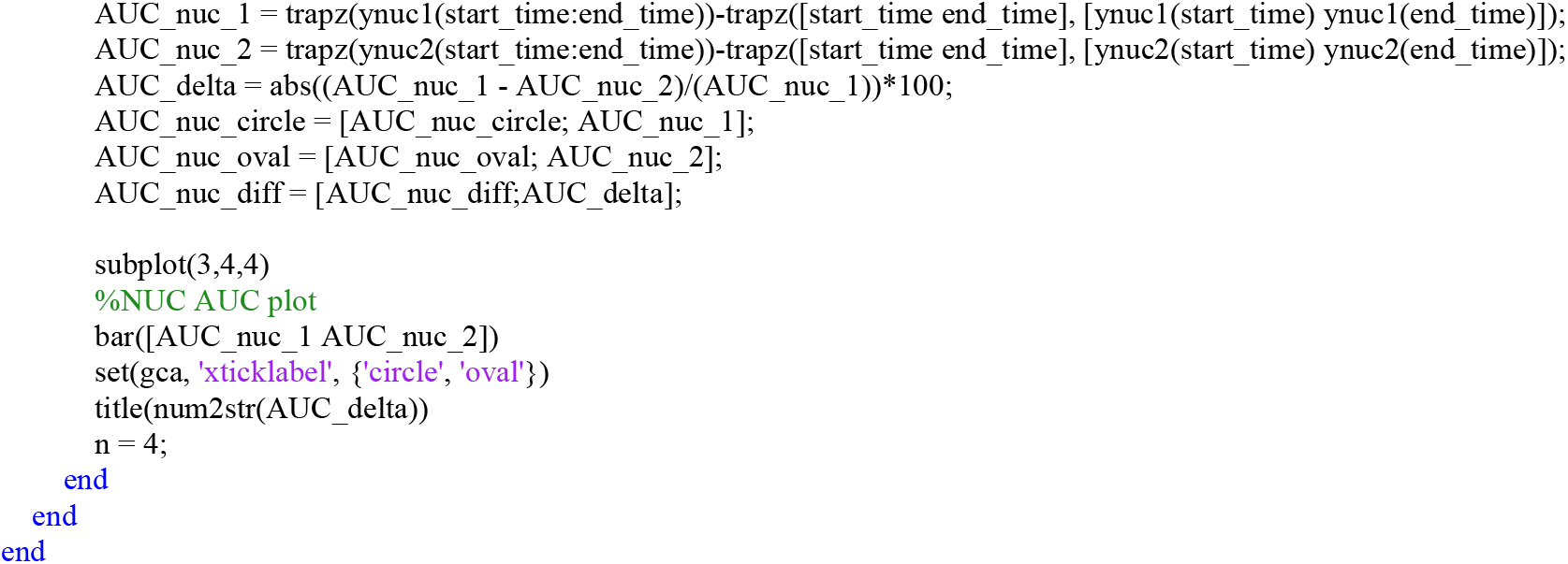

## REFERENCES

1. Neves, S. R., Ram, P. T. & Iyengar, R. G protein pathways. Science 296, 1636–1639 (2002).

2. Bhalla, U. S. & Iyengar, R. Emergent properties of networks of biological signaling pathways. Science 283, 381–387 (1999).

3. Rangamani, P. et al. Decoding information in cell shape. Cell 154, 1356–1369 (2013).

4. Neves, S. R. et al. Cell Shape and Negative Links in Regulatory Motifs Together Control Spatial Information Flow in Signaling Networks. Cell 133, 666–680 (2008).

5. Meyers, J., Craig, J. & Odde, D. J. Potential for Control of Signaling Pathways via Cell Size and Shape. Curr. Biol. 16, 1685–1693 (2006).

6. Kholodenko, B. N., Hancock, J. F. & Kolch, W. Signalling ballet in space and time. Nature Publishing Group 11, 414–426 (2010).

7. Kholodenko, B. N. Spatially distributed cell signalling. FEBS Lett. 583, 4006–4012 (2009).

8. Vilardaga, J.-P., Jean-Alphonse, F. G. & Gardella, T. J. Endosomal generation of cAMP in GPCR signaling. Nature Publishing Group 10, 700–706 (2014).

9. Villaseñor, R., Nonaka, H. & Del Conte-Zerial, P. Regulation of EGFR signal transduction by analogue-to-digital conversion in endosomes. Elife (2015).

10. Whitaker, M. Calcium at fertilization and in early development. Physiol. Rev. 86, 25–88 (2006).

11. Pinton, P., Giorgi, C., Siviero, R., Zecchini, E. & Rizzuto, R. Calcium and apoptosis: ER-mitochondria Ca2+ transfer in the control of apoptosis. Oncogene 27, 6407–6418 (2008).

12. Berridge, M. J. Calcium signalling and cell proliferation. Bioessays 17, 491–500 (1995).

13. Wei, C. et al. Calcium flickers steer cell migration. Nature 457, 901–905 (2009).

14. Berridge, M. J. & Taylor, C. W. Cold Spring Harbor Symp. Quant. Biol 53, 927–933 (1988).

15. Hirose, K., Kadowaki, S., Tanabe, M., Takeshima, H. & Iino, M. Spatiotemporal dynamics of inositol 1,4,5-trisphosphate that underlies complex Ca2+ mobilization patterns. Science 284, 1527–1530 (1999).

16. Berridge, M. J. Inositol trisphosphate and calcium signalling mechanisms. Biochim. Biophys. Acta 1793, 933–940 (2009).

17. Brozovich, F. V. et al. Mechanisms of Vascular Smooth Muscle Contraction and the Basis for Pharmacologic Treatment of Smooth Muscle Disorders. Pharmacol. Rev. 68, 476–532 (2016).

18. Kudryavtseva, O., Aalkjær, C. & Matchkov, V. V. Vascular smooth muscle cell phenotype is defined by C a2+-dependent transcription factors. FEBS J. 280, 5488–5499 (2013).

19. Hogan, P. G., Chen, L., Nardone, J. & Rao, A. Transcriptional regulation by calcium, calcineurin, and NFAT. Genes Dev. 17, 2205–2232 (2003).

20. Miranti, C. K., Ginty, D. D., Huang, G., Chatila, T. & Greenberg, M. E. Calcium activates serum response factor-dependent transcription by a Ras- and Elk-1-independent mechanism that involves a Ca2+/calmodulin-dependent kinase. Mol. Cell. Biol. 15, 3672–3684 (1995).

21. Wang, D. et al. Activation of cardiac gene expression by myocardin, a transcriptional cofactor for serum response factor. Cell 105, 851–862 (2001).

22. Rensen, S. S. M., Doevendans, P. A. F. M. & van Eys, G. J. J. M. Regulation and characteristics of vascular smooth muscle cell phenotypic diversity. Neth. Heart J. 15, 100–108 (2007).

23. Rzucidlo, E. M., Martin, K. A. & Powell, R. J. Regulation of vascular smooth muscle cell differentiation. J. Vasc. Surg. 45 Suppl A, A25–32 (2007).

24. Gomez, D. & Owens, G. K. Smooth muscle cell phenotypic switching in atherosclerosis. Cardiovasc. Res. 95, 156–164 (2012).

25. Chamley-Campbell, J., Campbell, G. R. & Ross, R. The smooth muscle cell in culture. Physiol. Rev. 59, 1–61 (1979).

26. Owens, G. K., Kumar, M. S. & Wamhoff, B. R. Molecular regulation of vascular smooth muscle cell differentiation in development and disease. Physiol. Rev. 84, 767–801 (2004).

27. Owens, G. K. Regulation of differentiation of vascular smooth muscle cells. Physiol. Rev. 75, 487–517 (1995).

28. Pitaval, A., Tseng, Q., Bornens, M. & Théry, M. Cell shape and contractility regulate ciliogenesis in cell cycle–arrested cells. J. Cell Biol. 191, 303–312 (2010).

29. Alford, P. W., Nesmith, A. P., Seywerd, J. N., Grosberg, A. & Parker, K. K. Vascular smooth muscle contractility depends on cell shape. Integr. Biol. 3, 1063–1070 (2011).

30. Polte, T. R., Eichler, G. S., Wang, N. & Ingber, D. E. Extracellular matrix controls myosin light chain phosphorylation and cell contractility through modulation of cell shape and cytoskeletal prestress. Am. J. Physiol. Cell Physiol. 286, C518–28 (2004).

31. Dickinson, G. D., Ellefsen, K. L., Dawson, S. P., Pearson, J. E. & Parker, I. Hindered cytoplasmic diffusion of inositol trisphosphate restricts its cellular range of action. Sci. Signal. 9, ra108 (2016).

32. Lock, J. T., Smith, I. F. & Parker, I. Spatial-temporal patterning of Ca2+ signals by the subcellular distribution of IP3 and IP3 receptors. Semin. Cell Dev. Biol. (2019) doi:10.1016/j.semcdb.2019.01.012.

33. Eglen, R. M., Reddy, H., Watson, N. & Challiss, R. A. Muscarinic acetylcholine receptor subtypes in smooth muscle. Trends Pharmacol. Sci. 15, 114–119 (1994).

34. Caulfield, M. P. Muscarinic Receptors—Characterization, coupling and function. Pharmacology & Therapeutics vol. 58 319–379 (1993).

35. Qin, D., Xia, Y. & Whitesides, G. M. Soft lithography for micro- and nanoscale patterning. Nat. Protoc. 5, 491–502 (2010).

36. Calizo, R. C. & Scarlata, S. A Role for G-Proteins in Directing G-Protein-Coupled Receptor–Caveolae Localization. Biochemistry 51, 9513–9523 (2012).

37. Isotani, E. et al. Real-time evaluation of myosin light chain kinase activation in smooth muscle tissues from a transgenic calmodulin-biosensor mouse. Proc. Natl. Acad. Sci. U. S. A. 101, 6279–6284 (2004).

38. Hanson, H. H., Reilly, J. E., Lee, R., Janssen, W. G. & Phillips, G. R. Streamlined embedding of cell monolayers on gridded glass-bottom imaging dishes for correlative light and electron microscopy. Microsc. Microanal. 16, 747–754 (2010).

39. Bell, M., Bartol, T., Sejnowski, T. & Rangamani, P. Dendritic spine geometry and spine apparatus organization govern the spatiotemporal dynamics of calcium. J. Gen. Physiol. 1017–1034 (2019) doi:10.1085/jgp.201812261.

40. Cugno, A., Bartol, T. M., Sejnowski, T. J., Iyengar, R. & Rangamani, P. Geometric principles of second messenger dynamics in dendritic spines. Sci. Rep. 444489 (2019) doi:10.1038/s41598-019-48028-0.

41. Bird, R. B., Stewart, W. E. & Lightfoot, E. N. Transport Phenomena. vol. 7 (Wiley, 2006).

42. Bressloff, P. C. & Newby, J. M. Stochastic models of intracellular transport. Rev. Mod. Phys. 85, 135–196 (2013).

43. Bressloff, P. C. & Earnshaw, B. A. Diffusion-trapping model of receptor trafficking in dendrites. Phys. Rev. E Stat. Nonlin. Soft Matter Phys. 75, 041915 (2007).

44. Pollard, T. D. & Cooper, J. A. Actin, a central player in cell shape and movement. Science 326, 1208–1212 (2009).

45. McBeath, R., Pirone, D. M., Nelson, C. M., Bhadriraju, K. & Chen, C. S. Cell shape, cytoskeletal tension, and RhoA regulate stem cell lineage commitment. Dev. Cell 6, 483–495 (2004).

46. Vogel, V. & Sheetz, M. Local force and geometry sensing regulate cell functions. Nat. Rev. Mol. Cell Biol. 7, 265–275 (2006).

47. Storm, C., Pastore, J. J., MacKintosh, F. C., Lubensky, T. C. & Janmey, P. A. Nonlinear elasticity in biological gels. Nature 435, 191–194 (2005).

48. Versaevel, M., Grevesse, T. & Gabriele, S. Spatial coordination between cell and nuclear shape within micropatterned endothelial cells. Nat. Commun. 3, 671 (2012).

49. Valm, A. M. et al. Applying systems-level spectral imaging and analysis to reveal the organelle interactome. Nature 546, 162–167 (2017).

50. Waterman-Storer, C. M. & Salmon, E. D. Endoplasmic reticulum membrane tubules are distributed by microtubules in living cells using three distinct mechanisms. Curr. Biol. 8, 798–806 (1998).

51. Wray, S. & Burdyga, T. Sarcoplasmic reticulum function in smooth muscle. Physiol. Rev. 90, 113–178 (2010).

52. Wu, Y. et al. Contacts between the endoplasmic reticulum and other membranes in neurons. Proc. Natl. Acad. Sci. U. S. A. 114, E4859–E4867 (2017).

53. Schauer, K. et al. Probabilistic density maps to study global endomembrane organization. Nat. Methods 7, 560–566 (2010).

54. Ehlert, F. J. Contractile role of M2 and M3 muscarinic receptors in gastrointestinal, airway and urinary bladder smooth muscle. Life Sci. 74, 355–366 (2003).

55. Mundell, S. J. & Kelly, E. The effect of inhibitors of receptor internalization on the desensitization and resensitization of three Gs-coupled receptor responses. Br. J. Pharmacol. 125, 1594–1600 (1998).

56. Hansen, S. H., Sandvig, K. & van Deurs, B. Clathrin and HA2 adaptors: effects of potassium depletion, hypertonic medium, and cytosol acidification. J. Cell Biol. 121, 61–72 (1993).

57. Heuser, J. E. & Anderson, R. G. Hypertonic media inhibit receptor-mediated endocytosis by blocking clathrin-coated pit formation. J. Cell Biol. 108, 389–400 (1989).

58. Voeltz, G. K., Rolls, M. M. & Rapoport, T. A. Structural organization of the endoplasmic reticulum. EMBO Rep. 3, 944–950 (2002).

59. Vasan, R. et al. Applications and Challenges of Machine Learning to Enable Realistic Cellular Simulations. Frontiers in Physics vol. 7 (2020).

60. Lee, C. T. et al. 3D mesh processing using GAMer 2 to enable reaction-diffusion simulations in realistic cellular geometries. PLoS Comput. Biol. 16, e1007756 (2020).

61. Toescu, E. C. & Verkhratsky, A. The importance of being subtle: small changes in calcium homeostasis control cognitive decline in normal aging. Aging Cell 6, 267–273 (2007).

62. Herring, B. P., El-Mounayri, O., Gallagher, P. J., Yin, F. & Zhou, J. Regulation of myosin light chain kinase and telokin expression in smooth muscle tissues. Am. J. Physiol. Cell Physiol. 291, C817–27 (2006).

63. Takashima, S. Phosphorylation of myosin regulatory light chain by myosin light chain kinase, and muscle contraction. Circ. J. 73, 208–213 (2009).

64. Mizuno, Y. et al. Myosin light chain kinase activation and calcium sensitization in smooth muscle in vivo. Am. J. Physiol. Cell Physiol. 295, C358–64 (2008).

65. Ye, G. J. C. et al. The contractile strength of vascular smooth muscle myocytes is shape dependent. Integr. Biol. 6, 152–163 (2014).

66. Crabtree, G. R. Calcium, calcineurin, and the control of transcription. J. Biol. Chem. 276, 2313–2316 (2001).

67. Sotiropoulos, A., Gineitis, D., Copeland, J. & Treisman, R. Signal-regulated activation of serum response factor is mediated by changes in actin dynamics. Cell 98, 159–169 (1999).

68. Liu, H. W. et al. The RhoA/Rho kinase pathway regulates nuclear localization of serum response factor. Am. J. Respir. Cell Mol. Biol. 29, 39–47 (2003).

69. Creemers, E. E., Sutherland, L. B., Oh, J., Barbosa, A. C. & Olson, E. N. Coactivation of MEF2 by the SAP domain proteins myocardin and MASTR. Mol. Cell 23, 83–96 (2006).

70. Ruhle, B. & Trebak, M. Emerging roles for native Orai Ca2+ channels in cardiovascular disease. Curr. Top. Membr. 71, 209–235 (2013).

71. Trebak, M. STIM/Orai signalling complexes in vascular smooth muscle. J. Physiol. 590, 4201–4208 (2012).

72. Croisier, H. et al. Activation of store-operated calcium entry in airway smooth muscle cells: insight from a mathematical model. PLoS One 8, e69598 (2013).

73. Bers, D. M. Cardiac excitation–contraction coupling. Nature 415, 198–205 (2002).

74. Scriven, D. R., Dan, P. & Moore, E. D. Distribution of proteins implicated in excitation-contraction coupling in rat ventricular myocytes. Biophys. J. 79, 2682–2691 (2000).

75. Sala, C. & Segal, M. Dendritic spines: the locus of structural and functional plasticity. Physiol. Rev. 94, 141–188 (2014).

76. Bakal, C., Aach, J., Church, G. & Perrimon, N. Quantitative morphological signatures define local signaling networks regulating cell morphology. Science 316, 1753–1756 (2007).

77. Sailem, H., Bousgouni, V., Cooper, S. & Bakal, C. Cross-talk between Rho and Rac GTPases drives deterministic exploration of cellular shape space and morphological heterogeneity. Open Biology vol. 4 130132 (2014).

78. Hansson, G. K. Molecular Cell Biology of Atherosclerosis. PanVascular Medicine 1–17 (2014) doi:10.1007/978-3-642-37393-0_6-1.

79. Chen, J. & Iyengar, R. Suppression of Ras-induced transformation of NIH 3T3 cells by activated G alpha s. Science vol. 263 1278–1281 (1994).

80. Mitra, A. et al. Cell geometry dictates TNFα-induced genome response. Proc. Natl. Acad. Sci. U. S. A. 114, E3882–E3891 (2017).

81. Jain, N., Iyer, K. V., Kumar, A. & Shivashankar, G. V. Cell geometric constraints induce modular gene-expression patterns via redistribution of HDAC3 regulated by actomyosin contractility. Proc. Natl. Acad. Sci. U. S. A. 110, 11349–11354 (2013).

82. Hassinger, J. E., Oster, G., Drubin, D. G. & Rangamani, P. Design principles for robust vesiculation in clathrin-mediated endocytosis. Proc. Natl. Acad. Sci. U. S. A. 114, E1118–E1127 (2017).

83. Sriram, K., Laughlin, J. G., Rangamani, P. & Tartakovsky, D. M. Shear-Induced Nitric Oxide Production by Endothelial Cells. Biophys. J. 111, 208–221 (2016).

84. Chabanon, M., Stachowiak, J. C. & Rangamani, P. Systems biology of cellular membranes: a convergence with biophysics. Wiley Interdiscip. Rev. Syst. Biol. Med. 9, (2017).

85. Rangamani, P., Levy, M. G., Khan, S. & Oster, G. Paradoxical signaling regulates structural plasticity in dendritic spines. Proceedings of the National Academy of Sciences 113, E5298–307 (2016).

86. Ron, A. et al. Cell shape information is transduced through tension-independent mechanisms. Nat. Commun. 8, 2145 (2017).

87. Heinrich, R., Neel, B. G. & Rapoport, T. A. Mathematical models of protein kinase signal transduction. Mol. Cell 9, 957–970 (2002).

## REFERENCES

1. Bird, R. B., Stewart, W. E. & Lightfoot, E. N. Transport Phenomena. vol. 7 (Wiley, 2006).

2. Kreyszig, E. Introduction to differential geometry and Riemannian geometry. (University of Toronto Press, 1969).

3. Bressloff, P. C. & Newby, J. M. Stochastic models of intracellular transport. Rev. Mod. Phys. 85, 135–196 (2013).

4. Bressloff, P. C. & Earnshaw, B. A. Diffusion-trapping model of receptor trafficking in dendrites. Phys. Rev. E Stat. Nonlin. Soft Matter Phys. 75, 041915 (2007).

5. Cooling, M., Hunter, P. & Crampin, E. J. Modeling Hypertrophic IP_3_ Transients in the Cardiac Myocyte. Biophys. J. 93, 3421–3433 (2007).

6. Vollmer, J., Menshykau, D. & Iber, D. Simulating organogenesis in COMSOL: Cell-based signaling models. arXiv [q-bio.QM] (2013).

7. Loew, L. M. & Schaff, J. C. The Virtual Cell: A software environment for computational cell biology. Trends in Biotechnology vol. 19 401–406 (2001).

8. Eungdamrong, N. J. & Iyengar, R. Compartment-specific feedback loop and regulated trafficking can result in sustained activation of Ras at the Golgi. Biophys. J. 92, 808–815 (2007).

9. Bhalla, U. S. & Iyengar, R. Emergent properties of networks of biological signaling pathways. Science 283, 381–387 (1999).

10. Li, Y. X. & Rinzel, J. Equations for InsP3 receptor-mediated [Ca2+]i oscillations derived from a detailed kinetic model: a Hodgkin-Huxley like formalism. J. Theor. Biol. 166, 461–473 (1994).

11. Fink, C. C. et al. An image-based model of calcium waves in differentiated neuroblastoma cells. Biophys. J. 79, 163–183 (2000).

12. De Young, G. W. & Keizer, J. A single-pool inositol 1,4,5-trisphosphate-receptor-based model for agonist-stimulated oscillations in Ca2+ concentration. Proc. Natl. Acad. Sci. U. S. A. 89, 9895–9899 (1992).

13. Lukas, T. J. A signal transduction pathway model prototype I: From agonist to cellular endpoint. Biophys. J. 87, 1406–1416 (2004).

14. Fink, C. C. et al. Morphological control of inositol-1,4,5-trisphosphate-dependent signals. J. Cell Biol. 147, 929–936 (1999).

15. Kapustina, M. et al. Mechanical and biochemical modeling of cortical oscillations in spreading cells. Biophys. J. 94, 4605–4620 (2008).

16. Quintana, A. R., Wang, D., Forbes, J. E. & Waxham, M. N. Kinetics of calmodulin binding to calcineurin. Biochem. Biophys. Res. Commun. 334, 674–680 (2005).

17. Lee, M. & Park, J. Regulation of NFAT activation: a potential therapeutic target for immunosuppression. Mol. Cells 22, 1–7 (2006).

18. Khalilimeybodi, A., Daneshmehr, A. & Sharif Kashani, B. Ca2+-dependent calcineurin/NFAT signaling in β-adrenergic-induced cardiac hypertrophy. Gen. Physiol. Biophys. 37, 41–56 (2018).

19. Rangamani, P., Levy, M. G., Khan, S. & Oster, G. Paradoxical signaling regulates structural plasticity in dendritic spines. Proceedings of the National Academy of Sciences 113, E5298–307 (2016).

20. Knot, H. J. & Nelson, M. T. Regulation of arterial diameter and wall [Ca2+] in cerebral arteries of rat by membrane potential and intravascular pressure. J. Physiol. 508 (Pt 1), 199–209 (1998).

21. Dickinson, G. D., Ellefsen, K. L., Dawson, S. P., Pearson, J. E. & Parker, I. Hindered cytoplasmic diffusion of inositol trisphosphate restricts its cellular range of action. Sci. Signal. 9, ra108 (2016).

22. Bers, D. M. Cardiac excitation-contraction coupling. Nature 415, 198–205 (2002).

23. Xu, C., Watras, J. & Loew, L. M. Kinetic analysis of receptor-activated phosphoinositide turnover. J. Cell Biol. 161, 779–791 (2003).

24. Golebiewska, U. et al. Membrane-bound basic peptides sequester multivalent (PIP2), but not monovalent (PS), acidic lipids. Biophys. J. 91, 588–599 (2006).

25. Sanabria, H., Digman, M. A., Gratton, E. & Waxham, M. N. Spatial diffusivity and availability of intracellular calmodulin. Biophys. J. 95, 6002–6015 (2008).

26. Hong, F. et al. Diffusion of myosin light chain kinase on actin: A mechanism to enhance myosin phosphorylation rates in smooth muscle. J. Cell Biol. 211, 2111OIA229 (2015).

27. Gorski, S. A., Dundr, M. & Misteli, T. The road much traveled: trafficking in the cell nucleus. Curr. Opin. Cell Biol. 18, 284–290 (2006).

28. Lee, Christopher T., et al. “3D mesh processing using GAMer 2 to enable reaction-diffusion simulations in realistic cellular geometries.” PLoS computational biology 16.4 (2020): e1007756.

